# A *cis*-regulatory element promoting increased transcription at low temperature in cultured ectothermic *Drosophila* cells

**DOI:** 10.1101/2020.10.15.340596

**Authors:** Yu Bai, Emmanuel Caussinus, Stefano Leo, Fritz Bosshardt, Faina Myachina, Gregor Rot, Mark D. Robinson, Christian F. Lehner

## Abstract

Cells of many ectothermic species, including *Drosophila melanogaster*, maintain homeostatic function within a considerable temperature range. The cellular mechanisms enabling temperature acclimation are still poorly understood. At the transcriptional level, the heat shock response has been extensively analyzed. The opposite has received less attention. Here, using cultured *Drosophila* cells, we have identified genes with increased transcript levels at the lower end of the readily tolerated temperature range, as well as chromatin regions with increased DNA accessibility. Candidate *cis*-regulatory elements (CREs) for transcriptional upregulation at low temperature were selected and evaluated with a novel reporter assay for accurate assessment of their temperature-dependency. Robust transcriptional upregulation at low temperature could be demonstrated for a fragment from the *pastrel* gene, which expresses more transcript and protein at reduced temperatures. The CRE is controlled by the JAK/STAT signaling pathway and antagonizing activities of the transcription factors Pointed and Ets97D.

## Introduction

Many habitats on earth experience temperature fluctuations of variable scale and temporal dynamics. Life in these thermally unstable habitats is a major challenge because temperature change affects all biological processes, but importantly not in a uniform manner. In general, diffusion is less temperature-dependent than enzymatic reactions, and the latter have individual temperature profiles. Thus, temperature acclimation is predicted to require complex regulation of cellular processes. Endothermic organisms, like humans, largely circumvent this challenge by keeping the core body temperature constant at the expense of metabolic energy. However, the large majority of organisms (microorganisms, plants and most animals including the model organism *Drosophila melanogaster*) are ectothermic. Their cells function over an often surprisingly wide range of ambient temperature. The cellular mechanisms enabling temperature acclimation are still poorly understood.

At the transcriptional level, considerable progress has been made in case of the heat shock response (HSR), which was discovered early on. Polytene chromosomes from salivary glands of *Drosophila busckii* larvae were observed to display a distinct chromosome puffing pattern in response to elevated temperature (Ritossa 1962). This evidence for transcriptional induction eventually led to the molecular isolation and characterization of the highly conserved *Hsp* genes (Vihervaara et al. 2018; Gomez-Pastor et al. 2018). Their products, the heat shock proteins (HSPs), function primarily as chaperones, which prevent or reverse protein misfolding and provide an environment for proper protein folding. The transcriptional activation of *Hsp* genes in response to heat shocks has served as an important experimental paradigm for research on the molecular mechanisms of transcriptional control. Heat shocks activate the transcription factor HSF1 and thereby cause a release of paused RNA polymerase II from sites downstream of the *Hsp* promoters into productive elongation. The HSR can be induced by stressors other than heat shock (like oxidative or heavy metal stress, glucose depletion). *Hsp* genes are also induced during recovery from severe cold shock (Liu et al. 1994; Colinet and Hoffmann 2010; Štětina et al. 2015; Heckel et al. 2016; Königer and Grath 2018). However, HSF1 is not responsible for the transcriptional induction of all of the many heat-shock induced genes in mammals and yeast (Mahat et al. 2016; Solís et al. 2016). More than half of the heat shock-induced genes are activated in a HSF1-independent manner in mammalian cells (Mahat et al. 2016). In addition, heat shock represses more genes than it induces, and the heat-induced repression is entirely HSF1-independent (Mahat et al. 2016).

In comparison to acclimation to elevated temperature, the opposite, i.e., the cellular response to temperature decrease has received less attention, in particular in animal organisms. For the relatively immotile microbial and plant organisms, cold is less avoidable and transcriptional responses have been characterized more extensively (Abduljalil 2018; Weber and Marahiel 2003; Ding et al. 2020; Ritonga and Chen 2020). In plants, considerable insight concerning crucial transcription factors (TFs) and their regulation by upstream thermosensors for responses to both extreme cold and more modest cool temperatures has been obtained. The ICE-CBF TFs regulate cold response (COR) genes important for cold tolerance in many plant species (Ritonga and Chen 2020). In case of vernalization, the process by which wintertime chill stimulates springtime flowering, epigenetic repression of the *FLC* locus by Polycomb factors is central in *Arabidopsis thaliana* (Whittaker and Dean 2017; Zhao et al. 2020). In thermomorphogenesis, i.e., the morphological changes according to ambient temperature, transcription factors (TFs) of the PIF family, which are controlled by both light and temperature, are crucial (Quint et al. 2016; Chung et al. 2020). Beyond PIFs, HSF family proteins also induce a large part of the warm transcriptome via eviction of +1 nucleosomes containing the histone H2A variant H2A.Z in temperature responsive genes (Cortijo et al. 2017; Kumar and Wigge 2010).

To characterize transcriptional responses to low temperature within the readily tolerated range in an ectothermic animal, we have chosen *Drosophila melanogaster*. For this fly, the temperature range for successful completion of the entire life cycle is commonly reported to be 14 to 29°C, and 25°C is considered to be optimal (Petavy et al. 2001). *D. melanogaster* and closely related species have already been used extensively for research on effects and responses to low temperature (Denlinger and Lee 2010; Hoffmann et al. 2003). In regard to transcriptional control, a majority of the published literature concerns exposure to severe cold, but a few studies have also reported transcriptome analyses within the readily tolerated range of 14-29°C (Chen et al. 2015; Jakšić and Schlötterer 2016; Fast et al. 2017). These studies performed with adult females or ovaries have detected extensive transcriptome changes. Analysis of whole animals and tissues likely augments the complexity of transcriptional responses. Diverse organs and cell types might express specific or even opposite responses to low temperature in case of particular pathways.

To reduce response complexity, we decided to use cultured cells of the *D. melanogaster* S2R+ cell line (Yanagawa et al. 1998) for analysis of transcriptional responses to temperatures at the lower end of the readily tolerated range. Temperature-dependence of the transcriptome was analyzed using DNA microarrays and 3’ RNA-Seq, and compared with that in adult male flies and another cell line. The temporal dynamics of the S2R+ cell transcriptome after a shift to 14°C was also analyzed. Combined with our data on temperature-dependence of DNA accessibility in chromatin acquired with ATAC-Seq (Buenrostro et al. 2013), we identified candidate *cis*-regulatory elements (CREs) driving transcriptional upregulation at low temperature. These CREs were further analyzed with a novel reporter assay for accurate evaluation of their temperature-dependency. Robust transcriptional upregulation at low temperature could be demonstrated in particular for a fragment from the *pastrel* (*pst*) gene, and its activity was found to be controlled by the JAK/STAT signaling pathway and the antagonizing activities of the transcription factors Pointed and Ets97D. Our work provides data resources and initial mechanistic insights into transcriptional control of acclimation to low temperature.

## Results

### The temperature range from 14 to 29°C is readily tolerated by *Drosophila* S2R+ cells

*Drosophila* cell lines, including S2R+ cells, are usually cultured around the presumed optimal temperature of 25°C. To assess the range of suboptimal temperatures permissive for proliferation of S2R+ cells (Fig. 1A), replicate cultures were plated, followed by incubation at different temperatures (17, 15, 13, 11, and 9°C). To monitor cell proliferation after the temperature shift, phase contrast micrographs of the same culture regions were taken at intervals (day 0, 1, 3, 5, 10 and 15). At 9, 11 and 13°C, a marginal increase in cell numbers was apparent. However, in parallel a clear increase in cell debris was observed in particular at 9°C, indicating substantial cell death (Fig. 1A). At 13°C, cells were increasingly more spindle shaped and a noticeable fraction of cells had vacuoles at the latest time point (day 15). At 15°C, cell proliferation was evident with only mild effects on cell morphology. Even stronger proliferation was obtained at 17°C, without obvious effects on cell morphology.

**Fig. 1.**
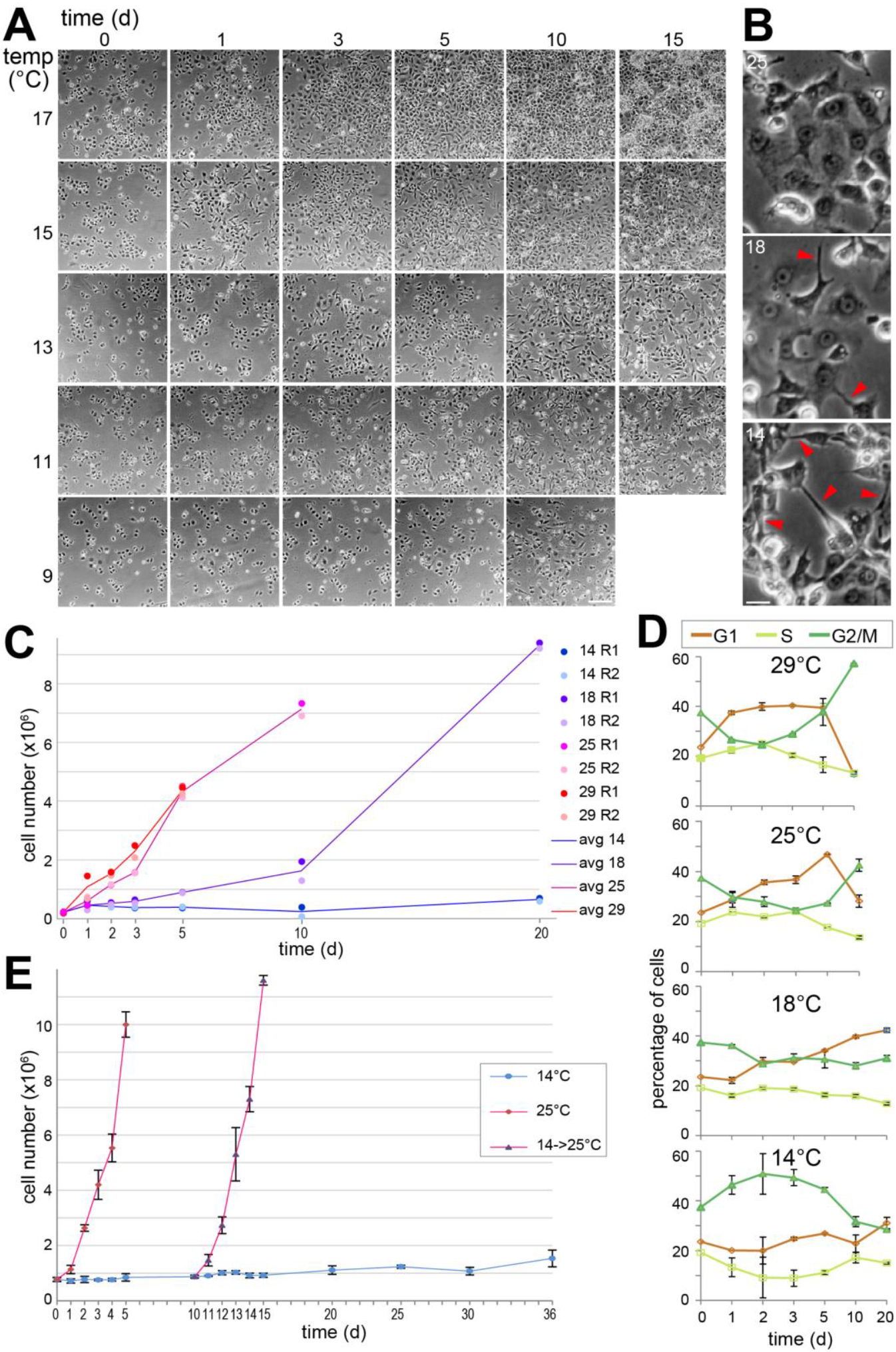
Proliferation of S2R+ cells at suboptimal temperatures. (**A,B**) For microscopic evaluation of cell proliferation, cells were plated in aliquots and shifted to the indicated suboptimal temperatures. Phase contrast images of the same regions were acquired at the indicated times after downshift. (**B**) High magnification views of cells grown to comparable cell density at the indicated temperatures illustrate increased cell aggregation and longer extensions (arrowheads) at low temperature. Scale bar = 50 μm (A) and 10 μm (B). (**C**) Cells were counted to assess cell proliferation at different temperatures (14, 18, 25 and 29°C). Culture aliquots were shifted to the different target temperatures, followed by counting at the indicated times after the shift. Two replicates (R1 and R2) were analyzed for each temperature and time point. Counts of viable cells in the two replicates and their mean are displayed. (**D**) Temperature effects on the cell cycle profile. Additional culture aliquots beyond those used for the counting shown in panel (C) were analyzed by flow cytometry after DNA staining for identification of cells in the G1, S and G2/M phase. Mean and s.d. are displayed (n = 2). (**E**) Immediate recovery of cell proliferation rate at 25°C after prolonged incubation at 14°C. Culture aliquots were shifted to either 14 or 25°C. Moreover, after 10 days of incubation at 14°C, some aliquots were transferred back to 25°C. Mean and s.d. of the counts of viable cells are displayed (n = 3).

For further analysis of temperature effects on S2R+ cell proliferation, we analyzed replicate cultures after shifts to different temperatures (29, 25, 18 and 14°C) apart from phase contrast microscopy by counting the number of live and dead cells. Moreover, flow cytometry was used for evaluation of cell cycle profiles, i.e. the fraction of cells in the G1-, S- and G2/M phase, respectively. Again, S2R+ cells were observed to be more spindle shaped overall with a greater cell aggregation tendency at the lowest temperature (14°C) (Fig. 1B). The number of live cells did not increase steadily at this lowest temperature (Fig. 1C). After an initial doubling within the first day, the number of live cells slowly returned to about the initial value until day 10, followed by another minor increase until day 20. The eventual increase in live cell number was observed in three independent experiments (with a fold change of 3.2, 1.9 and 1.4, respectively, between day 5 and 20). Flow cytometric analysis of cell cycle profiles over time indicated that S2R+ cells arrest in the G2 phase when entering the stationary phase after growth at 29 or 25°C (Fig. 1D). At 14°C, however, G2 cell enrichment was observed early after the temperature shift and the cell cycle profiles at the latest time points (10 and 20 days after the shift) were more similar to those observed during the proliferative phase at the higher temperatures (18, 25, and 29°C). For a further clarification whether S2R+ cells tolerate 14°C, we analyzed the dynamics of cell proliferation when returning cultures back to 25°C after 10 days of incubation at 14°C (Fig. 1E). Cell numbers increased rapidly after the transfer back to the optimal temperature, as fast as in those cells that had never been exposed to low temperature, confirming that 14°C does not result in substantial irreversible damage (Fig. 1E). Moreover, the percentage of dead cells at 14°C, which appeared to be slightly higher compared to 25°C (11.9% +/− 5.5 and 8.5% +/− 5.0, respectively, averaged over all replicates and time points analyzed in 4 independent experiments, i.e., n = 56 and n = 97, respectively) remained constant over time. Overall these findings suggest that S2R+ cells acclimate after a transfer from 25 to 14°C and resume cell proliferation at a very low rate eventually.

### Temperature effects on the S2R+ cell transcriptome

To analyze the effects of suboptimal temperatures on the S2R+ cell transcriptome, we used DNA microarrays for initial analyses. Aliquots of cells were plated at different temperatures (11, 14, 25, and 30°C) (Fig. 2A). Twenty-four hours after the temperature shift, RNA was isolated and analyzed. Three replicate experiments were preformed, resulting in a total of 12 samples. Comparison of the results obtained (S1 Table) revealed minimal differences between replicates of the same temperature treatment, as well as clear temperature effects (Fig. 2A). For a first estimate of the temperature regulated transcriptome, the number of differentially expressed (de) genes (fold change ≥ 2, false discovery rate (FDR) < 0.01) was determined by comparing the average expression level at the high temperatures (25 and 30°C) with that at the low temperatures (11 and 14°C) (S2 Table). Among 7350 clearly expressed genes, 698 (9%) had increased (CoolUp genes) and 1287 (18%) reduced (CoolDown genes) transcript levels in the cool conditions (Fig. 2B).

**Fig. 2.**
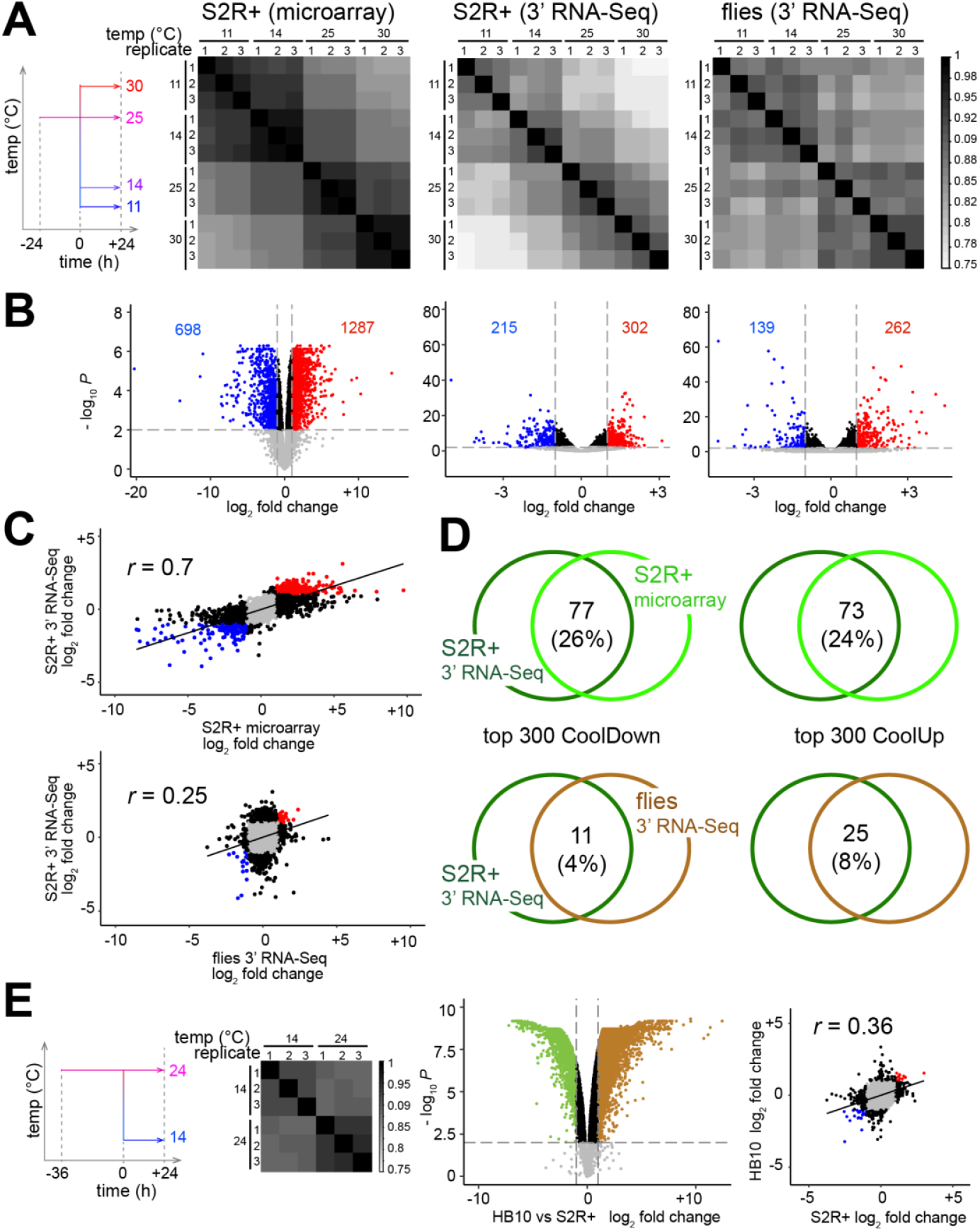
Temperature dependence of the transcriptome in different cell types (S2R+ and HB10) and adult male flies. (**A,B**) Aliquots of S2R+ cells and adult male flies were shifted for 24 hours to the indicated temperatures (11, 14, 25 and 30°C) before RNA isolation. Three replicates were analyzed in each experiment. A first experiment with S2R+ cells was analyzed with DNA microarrays, a second experiment with S2R+ cells and adult male flies with 3’ RNA-Seq. (**A**) Plots displaying the matrix of Pearson’s correlation coefficients, obtained after pairwise comparison of transcript levels in the different samples of a given experiment, revealed maximal similarities between replicates from the same temperature, more clearly for cells compared to flies. In addition, a greater similarity of the transcriptomes at the lower temperatures (11 and 14°C) compared to that at the higher temperatures (25 and 30°C) was evident in all experiments. (**B**) Volcano plots illustrate the number of CoolUp genes (blue dots) and CoolDown genes (red dots) with significantly different expression in a comparison between the lower and higher temperatures (FDR < 0.01; fold change ≥ 2). (**C,D**) Limited similarity of the transcriptome response to temperature change in S2R+cells and adult male flies. (**C**) Scatter plots of the fold changes of transcript levels (lower versus higher temperatures) observed in either the two experiments with S2R+ cells (top) or in the two 3’ RNA-Seq experiments with S2R+ cells and adult male flies (bottom). The temperature response of the transcriptome in the two S2R+ cell experiments was highly correlated, in contrast to poor correlation between S2R+ cells and adult males. *r* = Pearson’s correlation coefficient. (**D**) The extent of overlap among the top 300 CoolUp (left side) and CoolDown (right side) genes detected in either the two experiments with S2R+ cells (top) or in the two 3’ RNA-Seq experiments with S2R+ cells and adult male flies (bottom) is illustrated with Venn diagrams. The overlaps between the two S2R+ cell experiments were 3-6 fold larger than those between S2R+ cells and adult male flies. (**E**) Transcriptome changes in response to low temperature in HB10 cells. Culture aliquots were shifted to the indicated temperatures (14 and 24°C) before expression profiling with DNA microarrays. The Pearson’s correlation coefficients obtained after pairwise comparison of the different samples revealed maximal similarities between the three replicates from the same temperature. The volcano plots illustrates substantial differences between the transcriptomes of the different cell types HB10 and S2R+ at optimal temperature. Moreover, also the temperature response in the two cell types was poorly correlated, as indicated by the scatter plot of the fold changes in transcript levels (14°C versus optimal temperature) observed in the two cell types.

For validation of the microarray data, we repeated shifting S2R+ cells to different temperatures (11, 25, 30°C) and analyzed transcript levels of selected genes by quantitative reverse transcriptase polymerase chain reactions (qRT-PCR) (S1 Fig.). For validation we selected five CoolUp genes (*Lip4*, *Orct2*, *CG6321*, *CG13694*, *CG32944*), three CoolDown genes (*Hsp22*, *Hsp23*, *DNApol-α50*), and two genes (*Atg1*, *Pten*) with transcript levels at most marginally affected by temperature according to the microarray data. The results of the qRT-PCR analyses were well correlated with the microarray data (S1 Fig.), indicating their reliability.

When screening the microarray data visually for strong CoolUp genes with functions of interest concerning acclimation, we noticed that temperature appeared to affect polyadenylation in ten out of the first 35 such genes. In the ten genes, only those transcripts that were terminating at the most distal polyadenylation site (PAS) were CoolUp, while transcript isoforms resulting from more proximal polyadenylation were not (S2 Fig.). This observation raised the possibility that polyadenylation might be inefficient globally at low temperature, resulting in a bias towards more distal PASs. To clarify effects on alternative polyadenylation (APA), we applied 3’ RNA-Seq (Moll et al. 2014) after exposure of S2R+ cells to the same temperatures as before in the DNA microarray analysis (11, 14, 25, and 30°C). Moreover, for comparisons between cultured cells and flies, we also analyzed RNA samples isolated from aliquots of adult male flies after a 24 hour exposure at these temperatures (11, 14, 25, and 30°C). Before describing temperature effects on APA below, however, we will comment on the comparison of the results obtained with DNA microarrays and 3’ RNA-Seq for further illumination of reproducibility and cell-type specificity of the transcriptome response to temperature change.

Similar as with the microarray data, 3’ RNA-Seq data (S3 Table) from the three replicates at a given temperature was generally more strongly correlated than the data from different temperatures (Fig. 2A). For an initial comparison of the S2R+ cell data from microarray and 3’ RNA-Seq analyses, we first determined the number of de genes (fold change ≥ 2, FDR < 0.01) using the 3’ RNA-Seq data but also by comparing the average expression level at the high temperatures (25 and 30°C) with that at the low temperatures (11 and 14°C) (S2 Table). Among the total of 7133 clearly expressed genes detected, 215 (3%) were CoolUp and 302 (4%) CoolDown genes (Fig. 2B). For an additional comparison of S2R+ cells with flies, an analogous determination of de genes was done with the 3’ RNA-Seq data from adult male flies. 139 (1%) CoolUp and 262 (2%) CoolDown genes were found among 13168 clearly expressed genes detected in adult male flies (Fig. 2B). In case of the 3’ RNA-Seq data obtained from adult male flies (S4 Table), the overall correlation between replicates from the same temperature was not as pronounced as with S2R+ cells (Fig. 2A). However, the difference between the lower temperatures (11 and 14°C) and the higher temperatures (25 and 30°C) was clearly evident (Fig. 2A).

Applying the same arbitrary thresholds (fold change ≥ 2, FDR < 0.01) resulted in more de genes with microarray data from S2R+ cells in comparison to 3’ RNA-Seq data from S2R+ cells and flies (3.8 - and 4.9 fold, respectively). For further comparison, the overall correlation of the observed fold changes of all the genes/probes with clearly detectable expression in the different experiments was determined (Fig. 2C). For simplicity, we only considered the fold changes based on the comparison of the average expression at the low temperatures (11 and 14°C) with that at the high temperatures (25 and 30°C). The comparison between the two experiments with S2R+ cells (analyzed by microarray and 3’ RNA-Seq, respectively) yielded a considerably higher correlation coefficient than the comparison between S2R+ cells and male adult flies (both analyzed by 3’ RNA-Seq and hence not affected by platform-specific differences) (Fig. 2C). This finding emphasized that the transcriptome response to temperature change is strikingly different in adult flies compared to the highly reproducible response in S2R+ cells. This conclusion was further confirmed by the extent of overlap among the top de genes (Fig. 2D). The overlap among the top 300 CoolDown genes was found to be far greater in case of the comparison of the two S2R+ cell experiments (26%) than the corresponding overlap between S2R+ cells and adult male flies (4%) (Fig. 2D). Comparable findings were also made in case of the top 300 CoolUp genes (Fig. 2D).

Cell-type specificity of transcriptome responses to low temperatures was further indicated by analysis of an additional cell line, HB10. This cell line had been recently established from transgenic embryos with a method exploiting ubiquitous expression *UASt-ras85D*^*V12*^ driven by *Act5C-GAL4* (Simcox et al. 2008). Aliquots of HB10 cells were shifted to 14 and 24°C for 24 hours before transcriptome analysis by DNA microarrays (Fig. 2E) (S5 Table). Although HB10 and S2R+ are both derived from embryos, they are distinct cell lines, as indicated by the comparison of their optimal temperature transcriptomes (Fig. 2E). As previously reported (Cherbas et al. 2011), S2R+ cells are most similar to hemocytes. In contrast, HB10 cells displayed a transcriptome similar to that of adult muscle precursors (AMPs), as typically observed for cell lines generated with this particular method (Dequéant et al. 2015; Gunage et al. 2017). Illustrating this cell-type difference, the genes for the TFs Pannier (Pnr), Serpent (Srp) and Twist (Twi) were within the top 20 de genes when comparing S2R+ and HB10 cells at the optimal temperature. Pnr and Srp were high in S2R+ cells and low in HB10 cells. These TFs are master regulators of hemocyte development (Banerjee et al. 2019). In contrast, Twi was high in HB10 cells and low in S2R+ cells. Twi is crucial for mesoderm formation, remaining strongly expressed especially in AMPs (Gunage et al. 2017). The transcriptome changes in response to temperature downshift to 14°C were surprisingly distinct in S2R+ and HB10 cells. The overall correlation between the observed fold changes (expression at optimal temperature versus 14°C) in S2R+ and HB10 cells (Fig. 2E) was almost as low as that between S2R+ cells and flies (Fig. 2C). The overlap among the top 300 CoolUp and CoolDown genes was 20% in both cases. The overlap was further reduced to 1% by filtering for genes that were CoolUp or CoolDown not only in both cell types but also in adult male flies.

The 3’ RNA-Seq data allowed clarification whether low temperature globally suppresses polyadenylation efficiency. First, PAS were identified and assigned to genes (see Materials and Methods), and genes with multiple PASs were identified (APA genes) (S6 Table). In S2R+ cells, the number of APA genes identified was 2773. These APA genes were associated with two or more of the 14669 PASs detected in total (S3 Fig.). The APA genes represent 37% of the 7495 genes that are significantly expressed in S2R+ cells. In case of adult male flies, the number of significantly expressed genes (10757) and of PASs (22378), as well as the fraction of APA genes (40%, i.e., 4303) was higher than in S2R+ cells, as expected. For an identification of genes subject to temperature-dependent APA regulation, we determined the fold change in read counts when comparing low (mean of 11 and 14°C) and high (mean of 25 and 30°C) temperatures for each PAS using DEXSeq (Anders et al. 2012). For further analysis (Rot et al. 2017), we selected the two PASs with the most significant changes in case of genes with more than two changing PASs. If only one PAS had an adjusted p value < 0.05, then the second PAS was selected based on highest read count. If none of the PAS had a p value < 0.05, then both PASs were selected based on highest read count. In S2R+ cells, 812 (29%) of the APA genes displayed significant temperature-dependent changes in opposite directions at the two most strongly changing PASs. These genes with temperature-dependent APA were rather equally distributed onto the two classes with either the distal preferred over the proximal PAS at low temperature (class I, 47%) or vice versa (class II, 53%) (S3 Fig.). Similar results were obtained with adult males, where 662 (15%) of the APA genes displayed APA regulation by temperature with 49% in class I and 51% in class II (S3 Fig.).

In conclusion, temperature change is clearly accompanied by extensive changes in APA in *D. melanogaster*. Moreover, our results indicate that low temperature does not result in a global inhibition of polyadenylation. Preference for the most distal over the most proximal PAS was observed at a frequency comparable to the opposite.

APA usually results in sequence changes in 3’ untranslated regions, potentially altering target sites for miRNAs and RNA binding protein sites with consequences for mRNA stability, translation and localization (Tian and Manley 2017; Gruber and Zavolan 2019; Sadek et al. 2019). Some cases of APA also change the coding region. In addition, APA can affect formation of RNA secondary structure, which are of crucial importance for thermosensing in plants, bacteria and viruses (Chung et al. 2020; Kortmann and Narberhaus 2012; Somero 2018). The broad spectrum of potential APA consequences thwarts reliable bioinformatic predictions of physiological effects. Extensive experimental analyses will therefore be required to clarify the physiological significance of the observed temperature effects on APA. Moreover, effects on splicing, which are likely to augment the complexity of the transcriptome response to low temperature, will yet have to be analyzed comprehensively.

### Functional implications of transcriptome responses to low temperature

Data on changes in transcript abundance can often provide physiological insight by revealing significant enrichments of functional annotations. In case of S2R+ cells, the CoolDown genes were strongly enriched for functional annotations linked to DNA replication, mitosis and cell cycle progression. Indeed, inspection of the data for curated sets of bona fide S phase and M phase genes clearly confirmed reduced transcript levels at 14°C and even more strongly at 11°C (Fig. 3A, S4 Fig.). Compared to 25°C, the S and M phase genes were 3.8- and 2.5-fold down at 11°C on average. The marked downregulation of cell cycle genes after 24 hours at low temperature agreed well with the cell cycle profiles observed by flow cytometry, which revealed a comparable reduction in the fraction of S phase cells 24 hours after the temperature shift when 14°C was compared with 25 or 30°C (Fig. 1D).

**Fig. 3.**
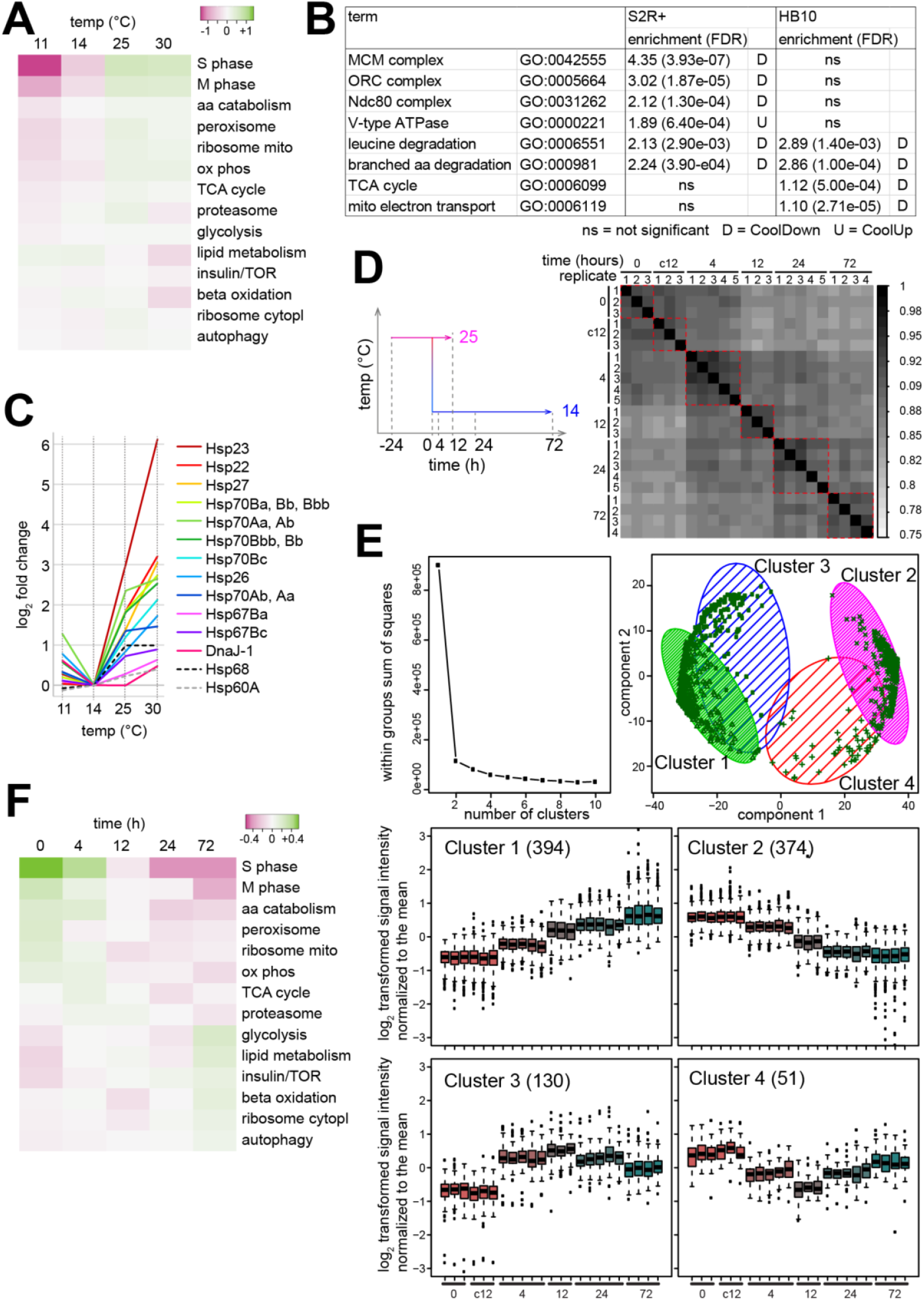
Cool temperature results in cell cycle gene repression but not stress gene induction in S2R+ cells. (**A**) Temperature dependence of transcript levels of genes functionally associated with central cellular processes. Microarray data for the S2R+ cell transcriptomes 24 hours after a shift to the indicated temperatures was used for an analysis of expression of curated sets of genes associated with the indicated central cellular processes. The heat map displays log_2_-transformed signal intensities of average fold change normalized to the means. (**B**) Enrichment of gene ontology terms by temperature-regulated genes reveal cell-type specific differences between S2R+ and HB10 cells. (**C**) Expression of *Hsp* genes, which are known to be strongly induced by heat shocks and additional stressors (like starvation, reactive oxygen species, infection and cold shocks), is minimal at 14°C in S2R+ cells. The log_2_ values of fold changes (average of three biological replicates and multiple probes, if present) relative to expression at 14°C are displayed, as revealed by the microarray data. Some microarray probes detect transcript derived from multiple *Hsp* genes. (**D**) Time course analysis of the transcriptional response to a 25->14°C temperature downshift in S2R+ cells using microarrays. Culture aliquots were analyzed at the indicated times after downshift (0, 4, 12, 24 and 72 hours), as well as samples maintained at 25°C for an additional 12 hours. The Pearson’s correlation coefficients obtained after pairwise comparison of the different samples revealed maximal similarities between the three to five replicates from the same time point (red dashed squares). (**E**) Analysis of gene clusters with similar temporal expression profiles in response to a 25->14°C temperature downshift. Time course microarray data from S2R+ cells was used for selection of probes with differential expression over time, followed by k-means clustering. The plot (top left) obtained by the elbow method (Goutte et al. 1999) after analysis of clusters resulting with increasing *k* revealed two pronounced clusters with limited distinct substructure. Additional plots illustrate the clusters obtained at *k* = 4, where two major clusters (1 and 2) with steadily increasing and decreasing signals, respectively, were detected, as well as two additional partially separated clusters (3 and 4) with a transiently increasing and decreasing signals, respectively. Number of probes assigned to the four clusters and normalized log_2_-transformed signal intensities detected in all the replicates are displayed in the four box plots. For normalization, the mean log_2_ signal intensity over all the samples obtained for a given probe was subtracted from the log_2_ signal intensity detected in the samples. “c12” indicates samples maintained for 12 hours at 25°C after t = 0 (temperature downshift). (**F**) Temporal profile of transcript levels derived from genes with functional association to central cellular processes. Microarray data from the time course analysis of the transcriptional response to a 25->14°C downshift in S2R+ cells was used for an analysis with curated sets of genes of central cellular processes. The heat map displays log_2_ values of average fold change at the indicated time points relative to expression at t = 0.

Beyond cell cycle progression, the de genes regulated by temperature in S2R+ cells did not strongly enrich additional functional annotations. For further corroboration that a temperature downshift to 14°C does not have pronounced transcriptional effects on central cellular pathways, we analyzed the temperature dependence of the average expression of curated sets of genes associated with various processes (glycolysis, TCA cycle, oxidative phosphorylation, mitochondrial ribosome, cytoplasmic ribosome, proteasome, autophagy, insulin/TOR, amino acid catabolism, lipid metabolism, beta-oxidation, and peroxisome) (Fig. 3A). The results clearly confirmed that in S2R+ cells these other processes were far less affected than cell cycle progression (Fig. 3A).

The analysis of enrichment of functional annotations further confirmed the high cell-type specificity of the transcriptome response to temperature, as suggested also by the observed limited overlap in de genes identified in S2R+ and HB10 cells (Fig. 2E). In stark contrast to the strong enrichment in S2R+ cells, annotations associated with cell cycle progression were not enriched by the de genes affected by a temperature downshift to 14°C in HB10 cells (Fig. 3B). Similarly, genes encoding subunits of the V-type ATPase were considerably CoolUp in S2R+ cells, but slightly CoolDown in HB10 cells. A further example for a discordant response concerns genes encoding subunits of the cytoplasmic ribosome. Although only marginally affected, they were overall CoolUp in S2R+ cells and CoolDown in HB10 cells. GO terms associated with the tricarboxylic acid cycle and mitochondrial respiration were clearly enriched among the CoolDown genes of HB10 cells but not in S2R+ cells. Among GO terms with clear enrichment (enrichment score ≥ 1), only two were found to be shared between S2R+ and HB10 cells. Both these shared terms were linked to degradation of branched amino acids (Leu, Val, Ile). Among the 61 genes subject to CoolUp regulation in both S2R+ and HB10 cells, while not causing strong enrichment of functional associations, we noted the presence of some with crucial roles in the control of glucose and lipid metabolism (like *Adipokinetic hormone* (*Akh*), *Insulin-like receptor* (*Inr*), *Pdk1*, *Hnf4*, *Mef2*, *ewg*/*NRF1*, *Pfrx* (Musselman and Kühnlein 2018; Mattila and Hietakangas 2017; Palanker et al. 2009; Clark et al. 2013). Overall, our transcriptome analyses indicate that the effect of temperature on transcript levels is surprisingly cell-type specific. We were unable to identify a set of genes that clearly and invariably responds at the transcriptional level to temperature change within the tolerated range in all three samples (adult male flies, S2R+ and HB10 cells).

To assess the kind and extent of stress that might be triggered in S2R+ cells by a temperature downshift to 14°C, we focused on known stress response genes. Cellular stress is predicted to increase sharply as temperature goes beyond the readily tolerated range. The heat shock protein genes (*Hsps*) are among the most extensively characterized stress response genes. While originally described in response to high temperatures, *Hsps* are also upregulated in response to severe cold in adult *D. melanogaster* (Liu et al. 1994; Colinet and Hoffmann 2010; Štětina et al. 2015; Heckel et al. 2016; Königer and Grath 2018). Therefore, the temperature-dependence of *Hsp* transcript levels should also be informative concerning the lower bound of S2R+ cells’ tolerated temperature range. If *Hsp* transcript levels were higher at 14 compared to 25°C, the former temperature would clearly appear as a stressful condition. However, transcript levels of all 12 *Hsp* genes inducible by heat shocks in cultured *Drosophila* cells (Gonsalves et al. 2011; Duarte et al. 2016) were found to decrease not only from 30 to 25, but also from 25 to 14°C (Fig. 3C; (Radermacher et al. 2014)). Strikingly, from 14 to 11°C, we observed an increase in transcript levels for 10 of the 12 *Hsp* genes, rather than a further drop (Fig. 3C). Thus, according to the transcript profiles of *Hsp* genes, 14°C appears to be within the well-tolerated range, in contrast to 11°C.

For further exploration of the extent of cellular stress imposed by 14°C, we performed a time-resolved analysis of the S2R+ transcriptome after a shift from 25°C to this suboptimal temperature. In the wild, *D. melanogaster* is presumably confronted mostly with relatively slow changes in ambient temperature, and hence adaptation of *Drosophila* cells for coping with rapid extensive step changes in temperature might be limited. Accordingly, a shift of S2R+ cell cultures from 25 to 14°C is predicted to be stressful initially. However, if 14°C were indeed within the well-tolerated range, cellular acclimation should succeed eventually, and reduce or even obliterate cellular stress. Genes characterized by a transcript increase that is only transient might therefore represent “stress genes”. In contrast, genes displaying persistent upregulation would qualify as "acclimation genes”; their regulation by temperature might be required for continuous homeostatic cell function at the suboptimal temperature. To study S2R+ transcriptome dynamics after a 25 to 14°C shift, we isolated RNA at different time points after the shift (0, 12, 24 and 72 hours) for probing DNA microarrays. Moreover, cells kept for 12 hours at 25°C rather than shifted to 14°C were included as well (t12_25 samples). Previous cell counting had indicated that equal cell densities were reached after 12 hours at 25°C and after 72 hours at 14°C, respectively. As some DNA microarrays failed to yield data of acceptable quality, the experiment was replicated so that valid results from at least three and up to five distinct biological replicates were available for each time point (S7 Table). Not surprisingly, cells maintained at 25°C and sampled at t = 0 or 12 hours, displayed an extremely similar transcriptome (Pearson’s correlation coefficient *r* = 0.998 in comparison of t0 with t12_25, log_2_ average signal intensities of the three replicates). To increase statistical robustness in the identification of genes responding to the 14°C shift, we treated the t12_25 data as additional t0 replicates. Compared to this initial time point, a total of 949 probes displayed signals affected by the temperature downshift when retaining only those with a fold change higher than two at one or more time points after the shift (t4, t12, t24, t72). We applied k-means clustering of these differentially expressed (de) probes for an identification of clusters of coregulated genes. As clearly indicated by the elbow method (Fig. 3E) (Goutte et al. 1999), de probes were segregated into two main clusters, which could not be partitioned readily into additional distinct subclusters. However, enrichment of GO terms associated with the resulting clusters was observed to be maximal at k = 4, providing some physiological support for a division into four clusters.

Among the four clusters resulting at k = 4, the two major clusters 1 and 2 contained de probes characterized by signals with an overall gradual increase (394 probes) or decrease (374 probes), respectively, after the temperature downshift (Fig. 3E). In contrast, the minor clusters 2 and 3 contained de probes with a transient increase (130) or decrease (51). The analysis of functional associations revealed by far the strongest and highly significant enrichments in case of cluster 2 (lowest FDR = 4.15e-26), because around a third of the 268 genes identified by the de probes in cluster 2 are well-known to provide functions important for progression through the cell cycle, most during S phase and several during M phase. Analysis of the selected bona fide S phase and M phase genes clearly confirmed this finding (Fig. 3F; S4 Fig.), which is also in full agreement with the initial transcriptome analyses at a single time point (24 hours) after a shift to various target temperatures (Fig. 3A). In contrast to cluster 2, enrichment of functional association terms was modest in case of the other three clusters (FDR at least 3.20e-21 fold higher compared to cluster 2). The de probes in cluster 1 with gradual upregulation after temperature downshift included genes with functions in extracellular matrix, cell adhesion and migration with modest but significant enrichment (FDR < 0.05), consistent with the observed increased spindle shape and clumping of S2R+ cells at 14°C (Fig. 1B). The CoolUp genes of cluster 1 also contained *betaTub97EF*, one of the limited number of genes with CoolUp regulation in both S2R+ and HB10 cells, encoding a beta tubulin isoform that stabilizes microtubules (Myachina et al. 2017). We emphasize that the GO term “response to stress” was not significantly enriched among the 272 gene identified by the de probes in cluster 1.

In contrast, an enrichment of stress genes was detected in case of cluster 3, although only marginally (p value 0.0065). The de probes in cluster 3 identified a total of 95 genes with an overall transient upregulation. Among these, the genes with functional annotations to stress response included *smp-30*, a gene identified early on as cold stress-induced (Goto 2000), *Hsp* genes, but only two (*Hsp26*, *DnaJ-1*) of the 12 heat-inducible *Hsp* genes, as well as *Keap1* (a conserved negative regulator of the response to oxidative stress) (Baird and Yamamoto 2020; Sykiotis and Bohmann 2008) and *Ets21C* (induced by the stress-responsive JNK pathway) (Külshammer et al. 2015). Overall, therefore, the clustering results were consistent with an occurrence of transient cellular stress and stress gene upregulation after the rapid step change from 25 to 14°C. However, the apparent stress response was strikingly limited. For further evaluation, we focused on stress response genes identified by previous transcriptomic analyses, in which adult flies or larvae had been exposed to various stressors (starvation; oxidative stress: paraquat, hydrogen peroxide, and hyperoxia; ER stress: tunicamycin; bacterial and fungal infection) (Girardot et al. 2004; Gregorio et al. 2001; Landis et al. 2004; Zinke et al. 2002). Our time course data did not reveal transient or persistent upregulation of such stress response genes (S5 Fig.). Similarly, overall the *Hsp* genes were observed to decrease over time with a minimum at 72 hours, the last time point analyzed after the temperature downshift (S5 Fig.). We also used antibodies against phosphorylated forms of JNK and p38a/b MAP kinases, which are well known to be activated by phosphorylation in response to many different types of stress including heat shock (Gonda et al. 2012). A transient activation was observed for both kinases with a maximum at 24 hours after the temperature downshift (S6 Fig.). In conclusion, S2R+ cells respond to a rapid step change from 25 to 14°C with a transient induction of stress response pathways. However, at least at the transcriptional level, the accompanying induction of known stress genes remains hardly detectable, indicating that 14°C is likely still within the well-tolerated range.

Acclimation to 14°C appears incomplete at 24 hours in S2R+ cells according to the temporal dynamics of the apparent stress response. Therefore, we inspected the time course data also for the selected genes acting in central metabolic pathways. This revealed changes in overall expression of genes in several functional networks that continued beyond 24 hours (Fig. 3F). However, compared to cell cycle genes, the overall expression changes of genes in other central pathways were clearly more limited.

### Temperature effects on DNA accessibility in chromatin of S2R+ cells

Elegant research with plants has revealed a potentially conserved mechanism for transcriptional control of temperature-regulated genes (Kumar and Wigge 2010; Cortijo et al. 2017). Around half of the transcriptome response to temperature was shown to be regulated by nucleosomes containing H2A.Z at the +1 position in *Arabidopsis thaliana*. Reduction of H2A.Z nucleosomes in mutants resulted in a transcriptome at low temperature that corresponded largely to that normally observed at warm temperature. Similar findings in budding yeast with mutations in the homologous *HTZ1* gene suggested that the crucial role of H2A.Z in the control of temperature-regulated genes is conserved. To evaluate whether the *Drosophila* homolog His2Av is equally central for transcriptional control in response to temperature change, we depleted this histone variant by RNA interference in S2R+ cells and analyzed the effect on the transcriptome after a temperature shift to different temperatures (14, 25 and 30°C) (S7 Fig.). Depletion reduced His2Av to a level estimated to be around 30% of normal levels (S7 Fig.). His2Av depletion had a pronounced effect on the transcriptome (S7 Fig., S8 Table). However, His2Av depletion did not transform the cool temperature transcriptome towards the warm transcriptome (S7 Fig.), suggesting that unlike in plants His2Av might not have a prominent role in the temperature-dependent control of transcription in S2R+ cells.

To identify genomic regions with temperature-dependent DNA accessibility, we used the assay for transposase-accessible chromatin using sequencing (ATAC-Seq) (Buenrostro et al. 2013). S2R+ cells were plated in aliquots at 25°C and shifted 36 hours later for an additional 24 hour of growth at either 14, 25 or 29°C (Fig. 4A). Thereafter, cells were harvested, crude nuclei were isolated and DNA accessibility was probed by tagmentation. All samples were tagmented at 25°C, avoiding data convolution by temperature effects on the enzymatic activity of the Tn5 transposase, but precluding detection of potential, rapidly reversible accessibility change. Three replicate experiments were performed. Accessible genome regions were identified as peaks of mapped ATAC-Seq reads. In total, 31745 total peaks and 17175 consensus peaks (i.e., those overlapping in at least two of the nine samples) were identified (S9 Table). As expected (Buenrostro et al. 2013), transcription start sites (TSS) were found to be highly enriched within the accessible ATAC-Seq peaks, indicating that our data reports DNA accessibility in chromatin (S8 Fig.). For identification of regions with temperature-dependent accessibility, we compared read counts in consensus peaks at 14 and 29°C. In total, 2317 (12.5%) with a significant difference in accessibility were found (FDR < 0.05) (S11 Table). The number of “CoolOpen” and “WarmOpen” regions were comparable (1189 and 1128, respectively) (Fig. 4B). At 25°C, these regions had an intermediate accessibility overall (Fig. 4C). Moreover, the fraction of consensus peaks with statistically significant temperature-dependent accessibility was dramatically lower when 25°C was compared to 29°C (0.03%) or to 14°C (0.03%), emphasizing that temperature change within the tolerated range results in limited and gradual accessibility differences.

**Fig. 4.**
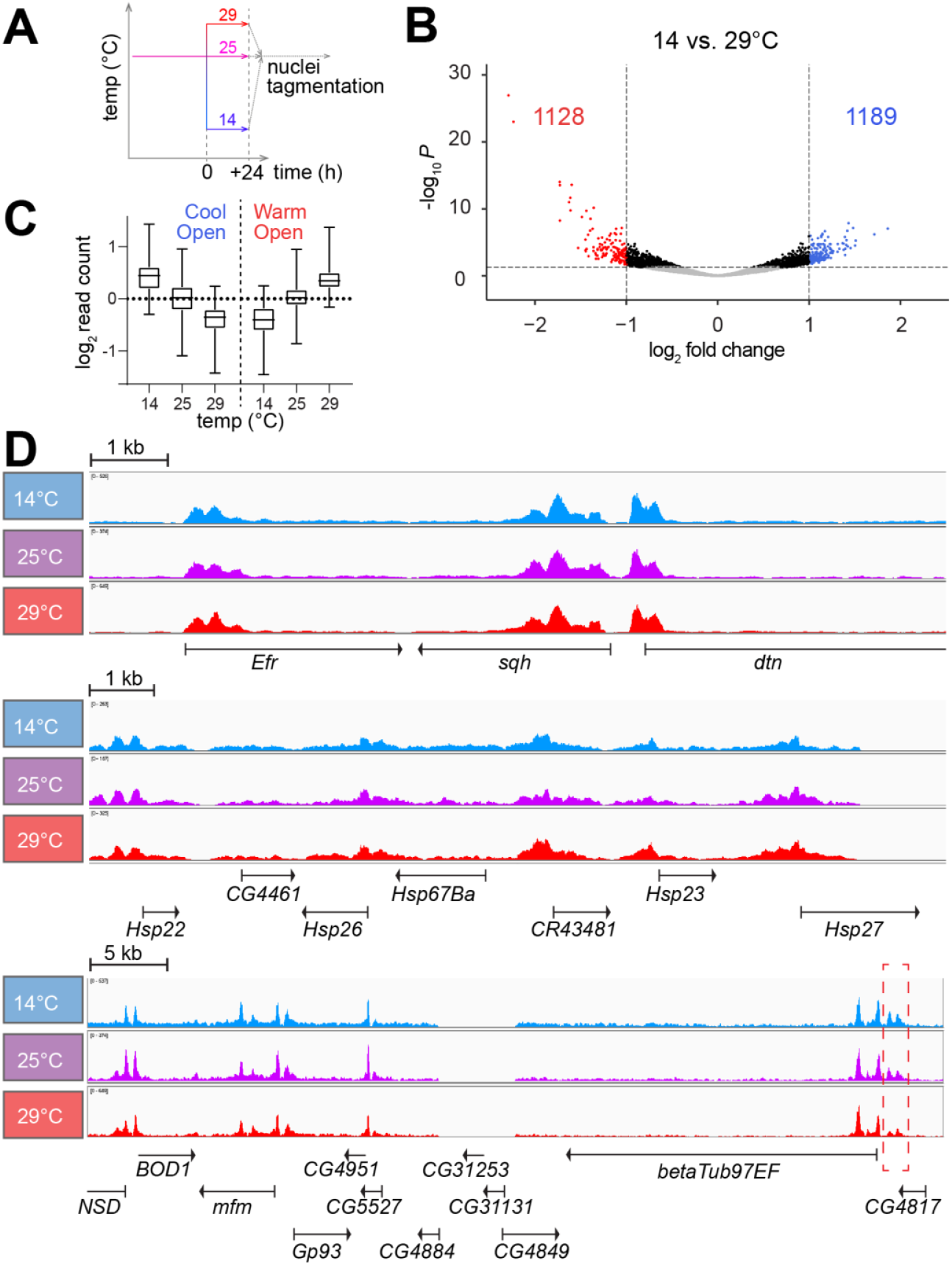
Temperature effects on DNA accessibility in nuclear chromatin of S2R+ cells. (**A**) Culture aliquots were shifted to the indicated temperatures and 24 hours later analyzed by ATAC-Seq involving tagmentation in crude nuclei always at 25°C. Three biological replicates were analyzed. (**B**) Volcano plots illustrate fold changes in read counts in ATAC-Seq peaks when comparing 14 with 29°C. Peaks with FDR ≥ 0.05 are shown in grey, those with a significant fold change ≤ 2 in black and those with a significant fold change > 2 in either blue (CoolOpen) or red (WarmOpen). (**C**) CoolOpen and WarmOpen regions identified as described in panel (B) have intermediate accessibility at 25°C on average. (**D**) Browser tracks display read counts obtained by ATAC-Seq at the indicated temperatures within selected genome regions. While the region shown in the top panel does not include temperature-regulated genes (including *sqh*), the region shown in middle panel contains small *Hsp* genes with transcript levels that were most strongly CoolDown. The bottom panel includes *betaTub97EF* with strongly CoolUp transcript levels (Myachina et al. 2017). Just upstream of this *betaTub97EF* gene, a CoolOpen region was apparent (dashed red rectangle). In contrast, at most modest accessibility alterations appear to be induced by temperature change in the other regions.

Visual inspection with a browser confirmed that the overwhelming majority of the ATAC-Seq peaks were not affected by temperature, as illustrated by a region (Fig. 4D) including the *cis*-regulatory elements (CRE) of the *sqh* gene used for control in some of our subsequent analyses. According to our expression profiling, transcript levels of *sqh* and the flanking genes are at most marginally affected by temperature. However, even in regions with genes strongly responding to temperature at the transcriptional level, parallel changes in DNA accessibility were rarely evident. The region including the *Hsp23* gene is presented for example (Fig. 4D). This region includes additional *Hsp* genes (like *Hsp22*, *Hsp26*, *Hsp27* and *Hsp67Ba*). As shown (Fig. 3C), all these *Hsp* genes have transcript levels correlated with temperature within the range of 14 to 30°C; *Hsp23* was in fact the CoolDown gene with the most extensive change in transcript levels genome-wide (fold change 14°C vs 30°C = 69.7). Nevertheless, in contrast to transcript levels, DNA accessibility in chromatin did not appear to be affected by temperature in the *Hsp23* region (Fig. 4D). In case of the *Hsp70* genes, which are also among the CoolDown genes (Fig. 3C), we also failed to detect significant accessibility changes (data not shown). In case of the strong CoolUp genes that were analyzed by inspection with the browser, we also failed in a clear majority to detect changes in DNA accessibility concurrent with transcript levels. We conclude that with the given sensitivity of our ATAC-Seq analysis the transcriptome response to temperature cannot be linked reliably with parallel changes in DNA accessibility at affected genes. However, we definitely also observed examples with an apparent correlation between transcript levels and DNA accessibility. In case of *betaTub97EF*, which was previously shown to be strongly CoolUp in S2R+ cells (Myachina et al. 2017), an increase in DNA accessibility was detected at low temperature within a region just upstream of the transcriptional start site (Fig. 4D).

### Assay for accurate analysis of temperature effects on CREs

For a further characterization of the mechanisms controlling the transcriptional response of CoolUp genes, an assay for accurate analysis of temperature effects on CREs appeared to be indispensable. The widely used dual luciferase assay was found to be problematic. The activity of Renilla luciferase expressed from a constitutive promoter (*Act5C* in our case), which is typically used for correction of assay variabilities including transfection efficiencies, was found to depend strongly on the temperature, at which cells were cultured before assaying. As an alternative with hopefully sufficient sensitivity for a detection of the gradual and limited transcriptional changes generally observed within the tolerated temperature range, we generated a cell line, in which GFP reporter transgenes could be assembled efficiently by site-directed chromosomal integration of a test DNA fragment, allowing measurement of its CRE activity by flow cytometry (Fig. 5A). Integration of the test fragment was achieved by directional recombinase-mediated cassette exchange (RMCE) with the two integrases PhiC31 and Bxb1. For RMCE, the test fragment was inserted in front of the *Drosophila* synthetic core promoter (DSCP) (Pfeiffer et al. 2008) between the two distinct integrase target sites. The resulting exchange plasmid was subsequently co-transfected into a special recipient cell line along with a dual integrase expression plasmid (pCo-PhiC31_Bxb1). The recipient cell line (SR9rg), a cloned S2R+ derivative, carried a chromosomal target locus for RMCE, which contained a constitutive mRuby transgene between the two integrase target sites, as well as a promoter-less mEGFP gene on one side just outside of the exchange region (Fig. 5A). Therefore, RMCE with test fragments that have enhancer activity will convert SR9rg cells from red into green fluorescent cells (Fig. 5B). Beyond the chromosomal RMCE target locus, the SR9rg cells were also transgenic for *MtnA_P-cas9*, allowing inducible *cas9* expression by addition of CuSO_4_ to the cell culture medium. Actually, the *cas9* construct had been stably integrated already before chromosomal insertion of the RMCE target locus, because this latter step was intended to be achieved by CRISPR/cas9-directed homologous recombination repair. However, SR9rg cells were found to have off-target integrations of the RMCE target sequences (see Materials and Methods).

**Fig. 5.**
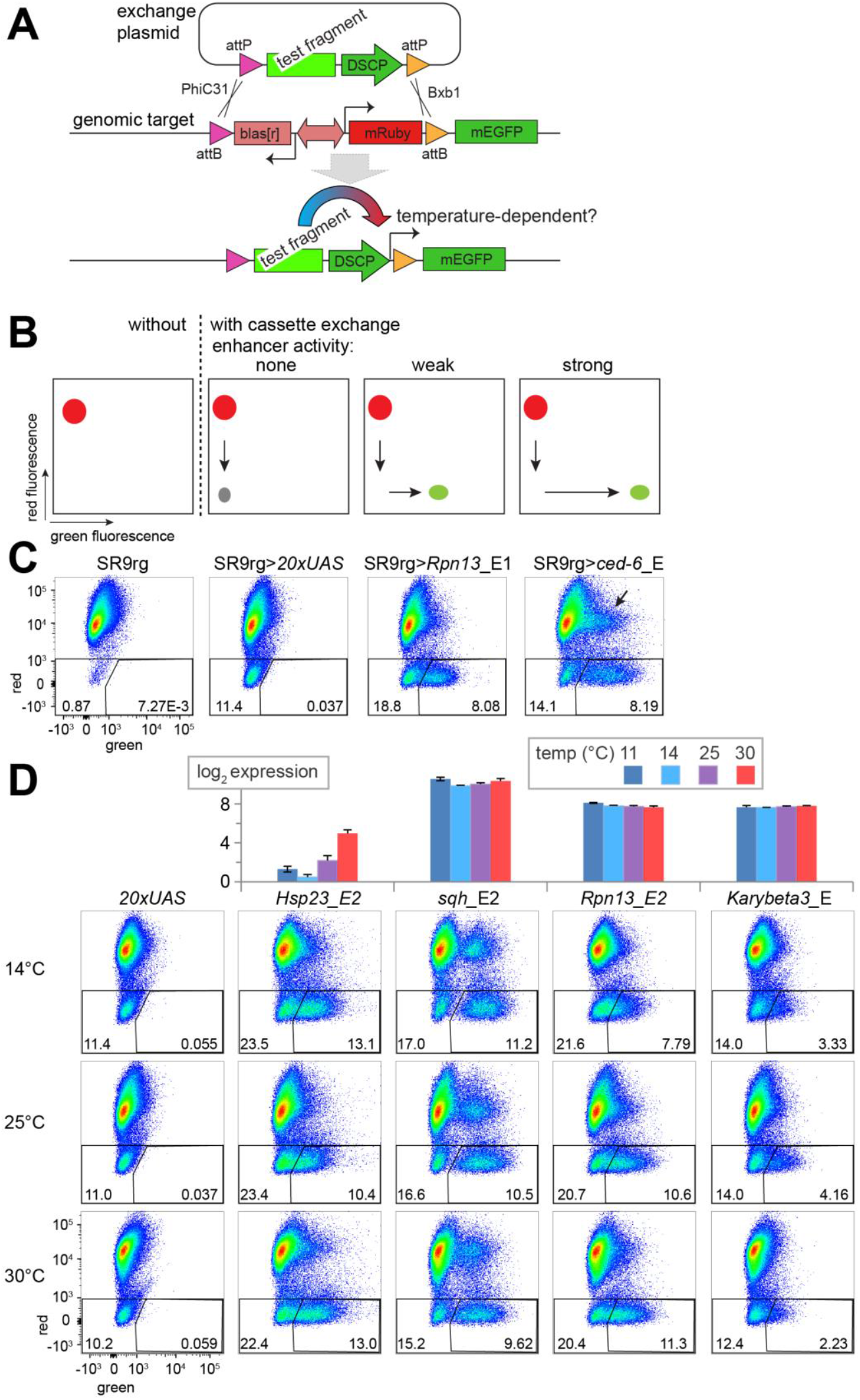
Assay for analysis of temperature dependence of CREs. (**A**) Scheme illustrating the complementation of a chromosomal mEGFP reporter gene by integration of a CRE test fragment into an engineered target locus by directional RMCE with SR9rg cells. After insertion of a candidate CRE into an exchange plasmid and cotransfection with a second dual integrase expression plasmid (not shown) for production of the two distinct PhiC31 and Bxb1 integrases, the chromosomal cassette with the bidirectional *blas*^*r*^ and *mRuby* marker genes can be replaced with the exchange plasmid cassette with the CRE in front of the *Drosophila* synthetic core promoter (DSCP). Thereafter, CRE activity can drive expression of green fluorescence in the resulting cell population, which can be cultured in aliquots at different temperatures before analysis by flow cytometry. (**B**) Analysis of enhancer activity of test fragments after RMCE with SR9rg cells by flow cytometry. Cartoons of scatter plots with red and green fluorescence intensity along x and y axis depict expected results (from left to right): SR9rg cells before RMCE express mRuby (red fluorescence) but not mEGFP (green fluorescence). RMCE with a test fragment lacking enhancer activity will generate a cell population with neither red nor green fluorescence (grey spot). However, RMCE with an enhancer fragment will result in a population that expresses only green fluorescence with an intensity reflecting enhancer activity. (**C**) Validation of the SR9rg assay system. Scatter plots display the results of flow cytometric analysis (from left to right): SR9rg cells before and after RMCE with the test fragments *20×UAS*, *Rpn13*_E1 and *ced-6*_E. The percentage of cells detected in the indicated windows is indicated (left window: cells without red and green fluorescence; right window: cells with only green fluorescence). In the rightmost scatter plot, a population of cells expressing both red and green fluorescence is indicated (arrow); see text for further explanations. (**D**) Analysis of temperature-dependence of CRE activity after RMCE with SR9rg cells. For assay validation, test fragments were also selected from genes with temperature-regulated transcript levels (*Hsp23*) or without (*sqh*, *Rpn13*, *Karybeta3*), as indicated by bar diagrams of 3’ RNA-Seq data on top. Aliquots of the cell populations obtained after RMCE in SR9rg cells with the selected test fragments were shifted either to 14, 25 or 30°C before flow cytometric analysis.

For validation that SR9rg cells permit quantitative analyses of temperature effects on CREs, we first generated four exchange plasmids with distinct test fragments (Fig. 5C). A first test fragment (*reg 069*) does not have enhancer activity in S2 cells according to a genome-wide analysis by STARR-Seq and subsequent confirmation with luciferase assays (Arnold et al. 2014). Similarly, as second fragment (*20×UAS*) with binding sites for the yeast transcription factor Gal4 was predicted to lack enhancer activity in *Drosophila* cells based on the extensive experience with the GAL4/UAS system in flies (Brand and Perrimon 1993). In contrast, the *ced-6*_E fragment has strong enhancer activity according to STARR-Seq and luciferase assays (Yáñez-Cuna et al. 2014). Finally, a fragment upstream of the *Rpn13* transcription start site was chosen; this region appeared likely to have enhancer activity based on position and accessibility according to ATAC-Seq. Each exchange plasmid was co-transfected along with pCo-PhiC31_Bxb1 into SR9rg cells for RMCE. Four weeks later, the resulting cell populations were analyzed by flow cytometry, along with non-transfected SR9rg cells. As expected, the large majority of the non-transfected SR9rg cells displayed red but not green fluorescence (Fig. 5C). The minor fraction of SR9rg cells without red fluorescence (0.8-2%) might reflect genetic instability, for example loss of the chromosome with the mRuby gene. After SR9rg transfection with only pCo-PhiC31_Bxb1 (but no exchange plasmid), the fraction of cells lacking red fluorescence increased slightly (to about 3%; data not shown), perhaps because of a low level of illegitimate integrase activity (Cherbas et al. 2015). After RMCE with the *reg 069* and *20×UAS* exchange plasmids, the proportion of cells without red fluorescence was further increased (Fig. 5C, data not shown), indicating that 10% of the analyzed cells were products of successful cassette exchange. A comparable increase in the fraction of cells without red fluorescence was also obtained after RMCE with the *Rpn13*_E1 and *ced-6*_E plasmids (Fig. 5C). Importantly, most cells lacking red fluorescence in these latter samples displayed increased green fluorescence (Fig. 5C). Average green fluorescence in SR9rg>*Rpn13*_E1 was lower compared to SR9rg> *ced-6*_E (Fig. 5C), indicating that enhancer activities can be assessed quantitatively with the cassette exchange system.

In a second validation step, we addressed whether temperature effects on CREs can be analyzed after RMCE with SR9rg cells (Fig. 5D). Temperature dependence of enhancer activity has hardly been studied in *Drosophila*. So far, a bona fide CoolUp enhancer has not yet been described to our knowledge. However, CREs from the *Hsp70* genes were shown to be heat shock responsive. Similarly, temperature dependence of CREs from *Hsp23*, the strongest CoolDown gene in S2R+ cells, has also been characterized to some extent (Pauli et al. 1986). The *Hsp23* CREs promote increased transcription at elevated temperature. Therefore, an exchange plasmid with a *Hsp23* CRE was generated. For comparison, we used exchange plasmids containing CREs from the loci *Karybeta3, Rpn13* and *sqh*. These genes do not respond to temperature change according to our expression profiling (Fig. 5D). After transfection for RMCE and three weeks of culture, the resulting cell populations were split into three aliquots and shifted 24 hours later to different temperatures (14, 25 and 30°C). Flow cytometry was performed after an additional 24 hour incubation period at these different temperatures. In case of the SR9rg>*Hsp23*_E2 cell population, the cells lacking red fluorescence were found to have more intense green fluorescence after incubation at 30°C compared to 25°C, and to a minor extent also when compared to 14°C (Fig. 5D). In comparison, the difference in green fluorescence intensity at 30 compared to 25°C was less pronounced in the cell populations lacking red fluorescence in case of the CREs from *sqh*, *Rpn13* and *Karybeta3* (Fig. 5D). In conclusion, our validation indicated that the RMCE system might be suitable for an evaluation of temperature effects on CREs. We note that enhancers with an assumed temperature-independent activity might behave as weakly CoolDown in our assay, as suggested by the comparison of green fluorescence intensities at 25 and 14°C in case of the CREs from *sqh*, *Rpn13* and *Karybeta3*. Alternatively, these enhancers might in fact be weakly CoolDown rather than temperature-invariant.

### A fragment from the pst locus confers robust transcriptional upregulation at suboptimal temperature

RMCE with SR9rg cells was used for an analysis of potential CoolUp enhancers, i.e., enhancers with higher activity at low temperature. Candidate CoolUp enhancer fragments were selected based on our data from expression profiling and ATAC-Seq. For example, ATAC-Seq had revealed a region around the transcriptional start site of *pastrel* (*pst*) with an increased accessibility at 14 compared to 25 and 30°C (Fig. 6A). The levels of *pst* transcripts were inversely correlated with temperature, in contrast to neighboring genes within the surrounding 40 kb (Fig. 6A,B). This inverse correlation was not only observed in S2R+ cells, but also in adult male flies, where *pst* was also CoolUp but not as pronounced (Fig. 6B). In HB10 cells, however, *pst* did not appear to be affected by temperature. Flow cytometric analyses with stably transformed S2R+ cells expressing an N-terminally tagged EGFP-Pst fusion protein under control of the *pst cis*-regulatory region demonstrated that *pst* expression is also CoolUp at the protein level (Fig. 6C).

**Fig. 6.**
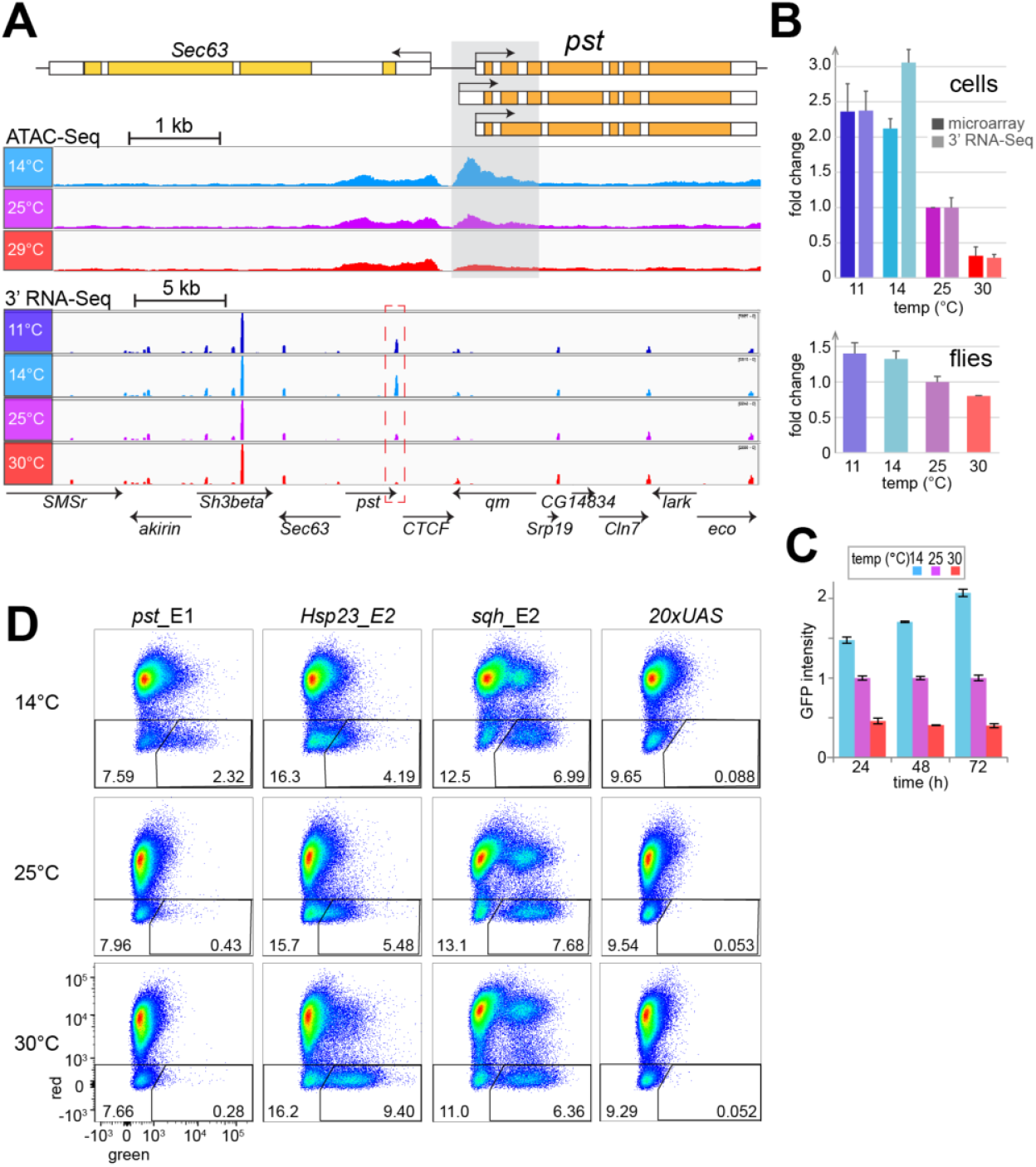
A fragment from the CoolUp gene pastrel (pst) has increased enhancer activity at low temperature. (**A**) Increased DNA accessibility at low temperature in region (grey shading) in the 5’ region of *pst*, a gene with higher transcript levels at low temperature. The genomic *pst* region is shown schematically (top), as well as browser tracks obtained from S2R+ cells at the indicated temperatures (average of three biological replicates) by either ATAC-Seq data (middle) or 3’ RNA-Seq (bottom). (**B**) Quantification of *pst* transcript levels at the indicated temperatures in S2R+ cells (top) and adult male flies (bottom). Results from two independent analyses, by microarray and 3’ RNA-Seq, respectively, are displayed in case of S2R+ cells. The data for flies was obtained by 3’ RNA-Seq. Average of three biological replicates and s.d. are shown, relative to expression at 25°C, which was set to 1. (**C**) Quantification of EGFP-Pst protein expression levels at the indicated temperatures by flow cytometry. Culture aliquots of S2R+_g-EGFP-*pst* cells were shifted to the indicated temperatures and analyzed at the indicated times after the shift. Average of three biological replicates and s.d. are shown, relative to expression at 25°C, which was set to 1. (**D**) Temperature dependence of the enhancer activity of the *pst*_E1 fragment (shaded region in panel A) analyzed after RMCE with SR9rg cells. For comparison the fragments *Hsp23*_E2, *sqh*_E2 and *20×UAS* were analyzed in parallel. After RMCE, cells were shifted eventually to the indicated temperatures and 48 hours later analyzed by flow cytometry. (see Figs.pdf)

To analyze whether the subregion from the *pst* locus that was characterized by increased accessibility at low temperature might confer CoolUp transcription, a corresponding fragment was inserted into the exchange plasmid and used for RMCE with SR9rg cells. The resulting SR9rg>*pst*_E1 cell population was incubated at different temperatures (14, 25, and 30°C) and analyzed by flow cytometry. In parallel, RMCE with the fragments *Hsp23*_E2, *sqh*_E2 and *20×UAS* was repeated and the resulting cell populations were subject to identical temperature treatment. The results obtained for the SR9rg>*pst*_E1 cell population revealed an inverse correlation of incubation temperature and green fluorescence in cells lacking red fluorescence. Green fluorescence was more prominent after incubation at 14°C, compared to 25 and 30°C (Fig. 6D). Conversely, as previously observed (Fig. 5), *Hsp23*_E2 resulted in increased green fluorescence after incubation at high temperature (Fig. 6D), while the *sqh*_E2 behaved as a strong enhancer largely unaffected by temperature, and *20×UAS* failed to display enhancer activity at any temperature. The enhancer activity of *pst*_E1 at different temperatures was robust; it was observed in all of a total of 18 independent transfections and flow cytometric analyses. It is concluded therefore that the *pst_*E1 fragment includes CREs that promote increased transcription at low temperature.

Beyond the *pst_*E1 region, we selected 31 additional regions with putative temperature-responsive CREs for analysis after RMCE with SR9rg cells (S9 Fig.). Three of these were selected because ATAC-Seq and expression profiling had suggested a potential presence of WarmUp enhancers. Reflecting our main aim, a greater number of regions with potential CoolUp enhancers was selected. Twenty eight such candidate regions were chosen applying somewhat variable criteria: 9 were CoolUp and CoolOpen, 5 were primarily CoolUp and 14 primarily CoolOpen. Several distinct subfragments were tested for some regions. Some fragments were tested in both orientations. Moreover, the DSCP promoter was omitted in case of some of the fragments, which already included an endogenous promoter region. As a result, the total number of assays for identification of enhancers with temperature-dependent activity was 48. The success rate was very low. The majority of the analyzed fragments (32, i.e., 66%), representing 26 distinct regions, displayed no or only marginal enhancer activity. Nine fragments from six distinct regions appeared to have enhancer activity, but only four were clearly temperature-dependent. One of the three regions selected as potential WarmUp CREs functioned as expected (*Prx2540-1*_E1) (S9 Fig.). In case of the 28 regions with putative CoolUp CREs, only one (*RNaseX25*) stimulated increased transcription at low temperature. This CoolUp activity was observed with three overlapping fragments (*RNaseX25*_E1, *RNaseX25*_E2 and *RNaseX25*_E3) (S9 Fig.).

In conclusion, only two CoolUp enhancers, one from *RNaseX25* and one from *pst*, were identified with our approach.

### The CoolUp CRE from pst is regulated by the JAK/STAT pathway and ETS family transcription factors

To delineate subregions within *pst*_E1 (968 bp) that are important for increased GFP reporter expression at low temperature, we analyzed a series of *pst*_E1 derivatives with truncations and central deletions after RMCE with SR9rg cells at distinct temperatures (S10 Fig.). A central region (372 bp) was found to be dispensable for CoolUp activity, but not the terminal regions (Fig. 7, S10 Fig.). Interestingly, CoolUp enhancer activity was also almost completely abolished by a larger central deletion (Fig. 7), indicating that essential enhancer sequences are located within a 308 bp region (Fig. 7, dark grey shading). While this central region is clearly essential, some terminal sequences might still contribute to the overall activity.

**Fig. 7.**
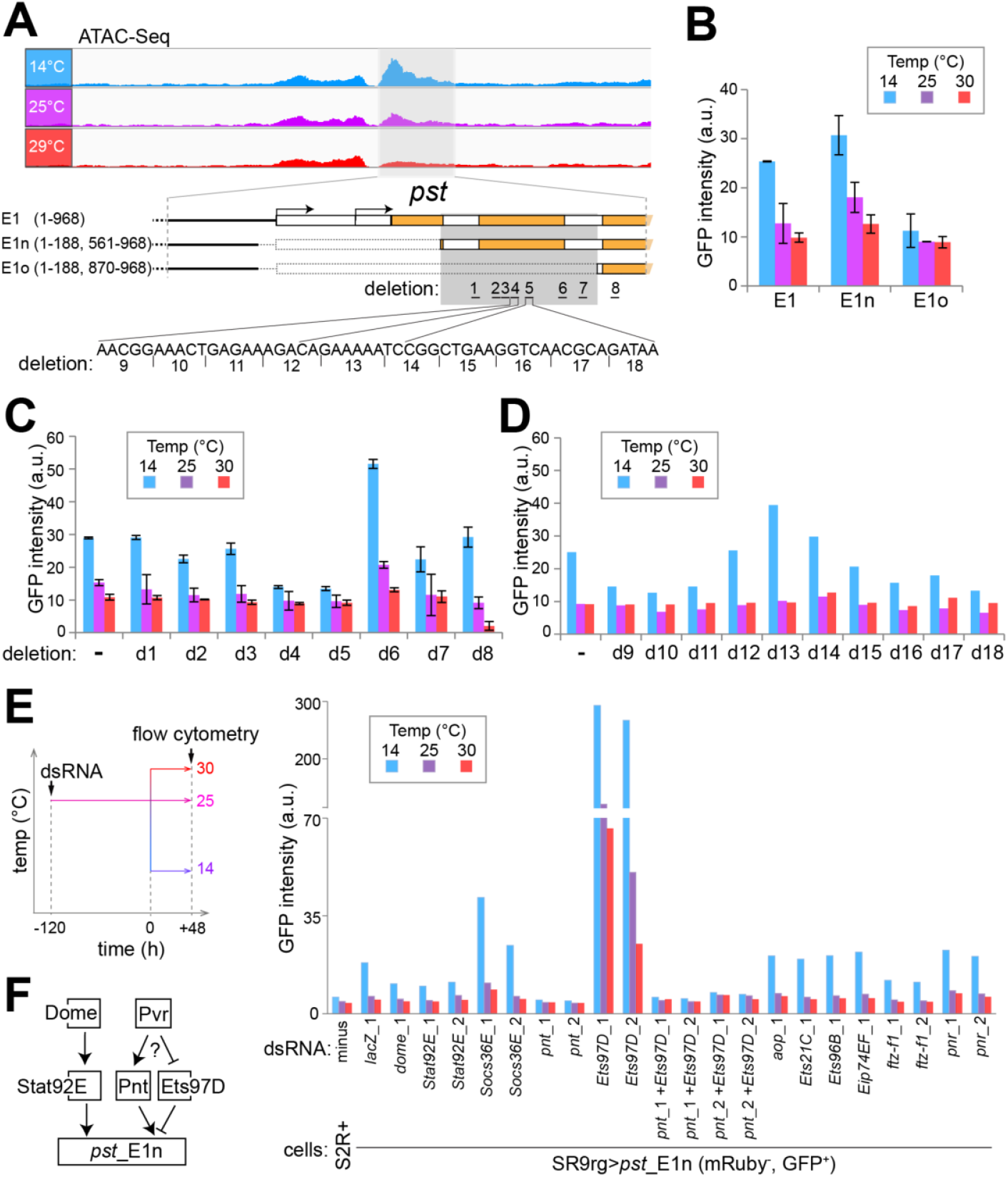
Characterization of the CoolUp enhancer from *pst*. (**A**) The CoolOpen region in the *pst* 5’ region identified by ATAC-Seq (*pst*_E1, light grey shading) with CoolUp enhancer activity was further dissected by analysis of terminal and internal deletion series. The comparison of E1n and E1o revealed an internal region essential for enhancer activity (dark grey shading). The internal deletions d1-8 were designed to eliminate predicted transcription factor binding sites, and the consecutive 5 bp deletions d9-18 for scanning the d4-d5 region. (**B-D**) Comparison of enhancer activity of E1 and derived fragments at the indicated temperatures as detected after RMCE with SR9rg cells. Bar diagrams display the median GFP signal intensity in the scatter plot window with cells expressing green but not red fluorescence. Analysis of the fragments E1, E1n and E1o (**B**), E1n derivatives carrying one of the deletions d1-8 (**C**), or one of the deletions d9-18 (**D**). Average of duplicates +/− s.d. shown in (B), and values from a single experiment in (D). (**E**) Characterization of the role of transcription factors (TFs) with predicted binding sites in E1n. The indicated candidate TFs were depleted in SR9rg> *pst*_E1n (mRuby^−^, GFP^+^) cells, followed by a shift of culture aliquots to the indicated temperature and subsequent flow cytometric analysis. Bar diagram represents median GFP intensity of the cell population lacking red fluorescence. In most cases, two independent dsRNA preparations generated from distinct amplicons (*xy*_1 and *xy*_2) were used for depletion. Untreated S2R+ cells and SR9rg> *pst*_E1n (mRuby^−^, GFP^+^) cells treated with *lacZ* dsRNA were used as negative and positive control, respectively. (**F**) Schematic summary model for the control of *pst*_E1 CoolUp enhancer activity. The JAK/STAT signaling pathway (with the transmembrane receptor Dome and the downstream TF STAT92E) acts positively. The TFs Pnt and Ets97D act as competing activator and repressor, respectively, presumably downstream of the Pvr receptor tyrosine kinase. (see Figs.pdf)

Further dissection of the essential 308 bp region was guided by a bioinformatic prediction of binding sites for known transcription factors (TFs) (Grant et al. 2011), which turned out to be clustered in seven subregions, each approximately 20 bp in length. To evaluate their importance, we analyzed a series of exchange plasmids with corresponding deletions (E1n_d1 to d7). An additional deletion (d8) outside the essential 308 bp region was made for analysis of a predicted STAT92E binding site. Deletion d8 eliminated only 3 bp, comprising the distal half of the STAT92E binding palindrom, in order to keep a predicted overlapping FOXO binding site intact. Flow cytometry indicated that deletion d5 eliminated enhancer activity (Fig. 7). Deletions d7, d4 and d8 also reduced enhancer activity although primarily at low temperature (Fig. 7). Conversely, the d6 deletion increased enhancer activity at all temperatures (Fig. 7).

As the d5 region had proven to be of particular importance, we generated an additional deletion series (E1n_d9 to d18) covering also the adjacent d4 region. Deletions were only 5 bp in length. Deletions d9 - 11 and d16 - 18 were found to decrease enhancer activity partially.

Overall, our dissection of *pst*_E1 with terminal truncations and internal deletions of varying size suggested that its CoolUp enhancer activity involves a complex interplay of multiple TFs.

For identification of TFs acting at the CoolUp enhancer from the *pst* locus, we depleted candidate TFs in SR9rg>*pst*_E1n cells by RNAi and analyzed the effects on GFP levels by flow cytometry. We selected TFs with binding sites predicted to be located in functionally important enhancer regions, as well as TFs encoded by genes with temperature-regulated transcript levels (S11 Fig.). Since several ETS family members (Hsu and Schulz 2000) were among the selected TFs (Aop, Ets21C and Ets97D), we included additional ETS proteins with predicted binding sites outside the essential region of *pst*_E1n, if expressed in S2R+ cells (Ets96B, Pnt and Eip74EF). As deletion d8 had supported an involvement of Stat92E, we selected beyond this TF also other proteins acting in the JAK/STAT signaling pathway (Herrera and Bach 2019), the transmembrane receptor Dome and the negative feedback regulator Socs36E for depletion experiments.

In a first experiment (S12 Fig.), where the SR9rg>*pst*_E1n cells were shifted after four days in the presence of dsRNA to either 14 or 25°C before flow cytometry, depletion of components of the JAK/STAT signaling pathway (Dome, Stat92E and Socs36E) and of some TFs (Pnt, Ets97D and Ftz-f1) had clear effects on *pst*_E1n enhancer activity. For confirmation, we performed a repetition experiment, in which we also examined additional temperature conditions (14, 25 and 30°C). Moreover, in case of factors implicated as functionally relevant by the first experiment, we included dsRNA treatment with a second distinct amplicon. Finally, we also performed double depletion of Pnt and Ets97D, because single depletions of these two ETS family members with overlapping predicted binding sites had given antagonistic effects in the first experiment.

The results of the second depletion experiments clearly confirmed those of the first. Accordingly, we conclude that the TFs Ftz-f1, Stat92E and Pnt function as positive, and Ets97D as negative regulator of *pst*_E1n enhancer activity (Fig. 7F). We point out that depletion of these TFs in SR9rg>*ced-6*_E cells, where GFP expression is driven by *ced-6*_E instead of *pst*_E1n, resulted (S12 Fig.) in either no effect on GFP fluorescence (in case of Ftz-f1 and Stat92E) or in far milder effects than those observed for *pst*_E1n (in case of Pnt and Ets97B). Thus, the identified TFs (Ftz-f1, Stat92E, and Pnt) presumably act at least to a considerable extent in a direct manner on *pst*_E1n enhancer activity.

The involvement of Stat92E in the control of *pst*_E1n enhancer activity appears to reflect its established function in JAK/STAT signaling. Depletion of additional JAK/STAT pathway proteins also had effects on *pst*_E1n enhancer activity, in directions consistent with their established functions. Dome, the upstream transmembrane receptor, qualified as a positive regulator and the negative feedback component Socs36E as an inhibitor. Expression of *dome* and *Stat92E* is not altered by temperature change in S2R+ cells or at most mildly according to our expression profiling (S11 Fig.). In contrast, we note that *Socs36E*, as well as *upd2* and *upd3* are clearly CoolUp (S11 Fig.) (and *upd1* expression was absent in S2R+ cells). The three *upd* genes code for the secreted ligand proteins that bind to the Dome receptor and activate JAK/STAT signaling in *Drosophila* (Herrera and Bach 2019).

Ets97D appears to counteract the positive regulator Pnt. Ets97D depletion resulted in strong stimulation of *pst*_E1n enhancer activity (about 15 fold at 14°C) (Fig. 7E). However, if in addition to Ets97D, Pnt was also depleted, enhancer activity was eliminated (Fig. 7E). Therefore, the strong stimulation of enhancer activity resulting from Ets97D depletion is entirely dependent on Pnt. As these two ETS family TFs have overlapping DNA binding specificity (Zhu et al. 2011), we propose that repressive Ets97D prevents binding of activating Pnt by competition for overlapping binding sites. Interestingly, the Pnt-P2 isoform is known to act as a transactivator downstream of various receptor tyrosine kinases (EGFR, Sevenless, Pvr and the FGFRs Heartless and Breathless) also by antagonizing an ETS family repressor, although not Ets97D, but rather Aop/Yan (Sopko and Perrimon 2013). Depletion of Aop, however, even though effective (S13 Fig.) did not affect *pst*_E1n enhancer activity (Fig. 7E, S12 Fig.). As the transcripts coding for the Pvr ligands Pvf2 and Pvf3 were CoolUp in S2R+ cells, we analyzed the effect of depletion of Pvf1, 2 and 3 as well as Pvr on *pst*_E1n enhancer activity (S11 Fig.). Pvr was found to be required for enhancer activity (S11 Fig.), suggesting that Pvr signaling might regulate Pnt activity and thereby also *pst*_E1n enhancer activity.

Overall, the results observed after depletion of TFs with predicted binding sites within *pst*_E1n confirmed that the function of this CoolUp enhancer depends on multiple TFs. Moreover, some of these TFs are likely regulated by signaling pathways that respond to secreted ligands. Depletion of Stat92E, Pnt and Ets97D in SR9rg>*pst*_E1n cells affected the level of transcripts derived from the endogenous *pst* gene in a manner analogous to the effects on the *pst_*E1n EGFP reporter (S13 Fig.). However, the effects were less pronounced, indicating the importance of additional regulatory inputs for transcriptional control of the endogenous *pst* gene.

## Discussion

The molecular mechanisms that allow cells in ectothermic animals a successful acclimation to variable environmental temperatures are likely complex. To avoid the complexities potentially caused by cell type- and tissue-specific responses in whole animals, we have focused here on S2R+ cells, a cell line derived from *Drosophila melanogaster* embryos. Moreover, we have studied transcriptional responses to temperatures below the optimum, as these have received less attention than the extensively studied heat shock response. However, rather than effects of extreme cold, our work concerns transcriptional responses to cool temperature within the readily tolerated range. The lower limit of this range in case of S2R+ cells appears to be similar as for flies. For both S2R+ cell proliferation and *D. melanogaster* propagation over generations, the lower limits are at around 14°C. Based on our analysis of the transcriptome and stress-activated kinases (JNK and p38), a rapid drop in ambient temperature from the optimal temperature (25°C) down to 14°C results in only a mild transient stress response in S2R+ cells. Eventual acclimation to this low temperature appears to evolve over several days with mostly gradual increases or decreases, respectively, in the transcript levels of hundreds of “CoolUp” and “CoolDown” genes. While CoolDown genes were mostly cell cycle genes, consistent with an almost complete halt of S2R+ cell proliferation at 14°C, CoolUp genes serve highly diverse functions. Interestingly, transcriptional changes were not accompanied by evident changes in chromatin organization detectable by ATAC-Seq, except for a minority of the temperature-regulated genes. To identify and characterize *cis*-regulatory elements (CREs) responsible for upregulation of CoolUp genes at 14°C, a reporter assay was developed where test DNA fragments were chromosomally integrated upstream of the *Drosophila* synthetic core promoter (DSCP) and the EGFP coding region by RMCE in SR9rg cells, an engineered clonal S2R+ derivative. Temperature-dependence of candidate CREs was assessed by flow cytometry after culturing aliquots of the resulting reporter cell population at distinct temperatures. Thereby, only two out of 29 candidate CoolUp CREs were found to result in a robust increase in reporter EGFP expression at 14°C in comparison to the optimal temperature. By additional characterization of one of these two, a fragment from the *pastrel (pst)* locus, the TFs Stat92E, Pnt and Ets97D were found to be crucial for the function of this CoolUp enhancer. We suggest that the activity of these TFs is controlled by JAK/STAT and RTK signaling pathways.

While intriguing, the similarity of what we have designated the “readily tolerated temperature range” in S2R+ cells and flies should not be overrated. The genetic stability in *Drosophila* cell culture lines is clearly lower than in flies. Genome resequencing of close to twenty distinct cell lines has exposed extensive copy number changes (Lee et al. 2014). While mostly cell line-specific, some changes were common to many lines, including a high copy region encompassing *Pvr*, which has been suggested to reflect selection for provision of protection against apoptosis by this gene. Clearly, drift and altered selection pressure during S2R+ cell propagation might have modified some of the responses to temperature. However, *Drosophila* cell lines retain characteristics of the tissue of origin according to transcriptome comparisons between cell lines and tissues (Cherbas et al. 2011). Accordingly, the low similarity in the transcriptional response to temperature between adult male flies, S2R+ and HB10 cells that we have observed, might reflect genetic or epigenetic differences or both. In any case, it indicates a high plasticity in this response, precluding instant wide-ranging generalizations. Our 3’ RNA-Seq analyses with S2R+ cells and adult male flies, as well as previous RNA-Seq studies with adult female flies (Chen et al. 2015; Jakšić and Schlötterer 2016), emphasize that acclimation within the readily tolerated temperature range is accompanied by transcriptome changes beyond transcript abundance. Extensive changes in alternative mRNA splicing and polyadenylation further increase the response complexity.

Based on the transcriptional response of genes previously shown to be strongly induced by various types of stress (heat, ROS, ER failure, infection and starvation), incubation of S2R+ cells at 14°C appears to be at most mildly stressful. However, progression through the cell cycle was severely inhibited after a shift to 14°C. Transcript levels of S- and M phase genes were strongly decreased and flow cytometry revealed a transient G2 arrest. S2R+ cells also stall in G2 when reaching high density during growth at the optimal temperature. Transient cell cycle arrest in G2 is frequently observed in *D. melanogaster* during normal development and in response to stress (Wartlick et al. 2011; Guo et al. 2016; Otsuki and Brand 2018; Cosolo et al. 2019; Edgar and O’Farrell). In contrast to S2R+ cells, cell cycle genes were not downregulated when HB10 cells were grown at 14°C. This cell line, which we have generated from embryos using transgenic transformation with activated Ras (Simcox 2013), has a considerably higher cell doubling time at the optimal temperature compared to S2R+ cells. As HB10 cells were generated recently, they have not been exposed to selection for rapid proliferation in culture as intense as the S2R+ cells, which have been passaged during the past 20 years. The distinct history might well be responsible for the striking difference in temperature-dependence of cell cycle gene expression in these two cell lines. It is not excluded, however, that cell-type differences contribute as well. S2R+ cells are most similar to hemocytes and HB10 cells to adult muscle precursors.

To evaluate whether temperature-dependent transcriptome change is linked to alteration of DNA accessibility in chromatin, we have applied ATAC-Seq after incubation of S2R+ cells at different temperatures. Around 2300 ATAC-Seq peaks were significantly affected by temperature when comparing 14 with 29°C. Although numerically greater than the number of temperature-regulated genes, such differentially accessible regions were often conspicuously absent from chromosomal locations with differentially expressed genes. The extensive differences in *Hsp* transcript levels for example were associated with at most marginal changes in chromatin accessibility. This observation is in full agreement with the findings concerning the molecular mechanisms, which mediate heat shock-induced upregulation of *Hsp70* expression (Vihervaara et al. 2018). Extensive analyses have clearly established that the regulated step consists in the release of RNA polymerase II from a promoter-proximal pause site into productive elongation. While the recruitment of RNA polymerase is linked to prominent chromatin opening within the cis-regulatory region around the TSS, all subsequent steps (initiation, pausing, release and elongation) occur with minimal changes in DNA accessibility (Vihervaara et al. 2018). The *Hsp70* promoter region is therefore already highly accessible in the non-induced state. This paradigm is known to govern the regulation of very a substantial fraction of all genes (Vihervaara et al. 2018; Adelman and Lis 2012). Conceivably, it applies to many of the genes that are subject to temperature-dependent regulation within the range of 14-30°C. Although we failed to detect significant changes in DNA accessibility at most of the temperature-regulated loci, there were also clear exceptions which displayed correlated changes in DNA accessibility and transcript levels in response to temperature change. Moreover, in case of the *pst* locus, a region with CoolUp DNA accessibility was also found to have CoolUp enhancer activity.

For precise quantitative analyses of temperature effects on the activity of candidate CREs, we have developed a novel reporter assay. In this assay, DNA fragments of interest are first inserted into an exchange plasmid, which is then co-transfected with an integrase expression plasmid for RMCE-mediated transfer of the fragment into engineered target sites in the genome of SR9rg cells. This targeted integration strategy increases assay reproducibility compared to analyses after transient transfection with or without selection of random integrations. With our assay, comparison of an original with a mutated derivative of an enhancer fragment for example is far less convoluted by unavoidable variabilities in transfection efficiency, integration site and copy number. Recently, several similar *Drosophila* cell lines allowing targeted RMCE-mediated integration of DNA fragments were described (Cherbas et al. 2015; Manivannan et al. 2015; Neumüller et al. 2012; Viswanatha et al. 2018). These cell lines carry single target site for non-directional RMCE. In contrast, our SR9rg cell line has multiple target sites, which can be disadvantageous for certain applications, but our target sequence includes special features designed for enhancer analysis. Directional RMCE provides control over fragment orientation. Moreover, since the EGFP reporter coding region is part of the target site and hence absent from the exchange plasmid for test fragment delivery, random off-target integrations of this exchange plasmid cannot contribute to reporter expression. Therefore, no selection against such variable off-target integrations is required. This is highly advantageous because in cultured *Drosophila* cells random integrations occur frequently and methods for selection against them are inefficient (Cherbas et al. 2015; Manivannan et al. 2015) (Y.B and C.F.L., unpublished observations). For analysis of CREs, the design of our target site results in the following constraints. For analyses of enhancer activity, the positioning of test fragments downstream of the reporter gene is usually preferred. In our assay system, test fragments are necessarily upstream, as we aimed for an approach that permits an analysis also of promoter activities. However, if the test fragment includes a TSS and a downstream ATG motif, the latter needs to be in frame with the EGFP coding sequence and free of downstream stop codons, or else potential transcriptional stimulation by the test fragment cannot be detected. Problems with TSSs followed by out-of-frame ATGs and premature stop codons might be bypassed by inversion of a given test fragment in front of the DSCP, although interference by promoter competition might then arise. The fraction of putative CoolUp CREs that might have compromised our assay because of promoter competition is likely substantial, as our selection of candidate CRE fragments was based to a large part on ATAC-Seq peaks covering the region with annotated TSSs. We suggest that the observed low rate of successful validation of putative CoolUp CREs is explained in part by such technical problems.

Apart from some limitations as discussed, our reporter assay involving flow cytometric EGFP signal quantification with SR9rg cells after integration of test DNA fragments by RMCE is highly reproducible. Moreover, by incubation of culture aliquots at different temperatures before flow cytometry, temperature-dependence of CREs can be assessed accurately. Thereby, we have been able to demonstrate that the *pst*_E1 fragment stimulates EGFP reporter expression more effectively at 14°C than at 25 and 30°C. We are not aware of any other CoolUp enhancers previously identified in *D. melanogaster*, and the number described in other animal species appears to be low (Sumitomo et al. 2012; Thaisuchat et al. 2011). By further analysis with the *pst*_E1n subfragment, we have uncovered an involvement of the JAK/STAT pathway and the ETS family proteins Ets97D and Pnt in the function of this CoolUp enhancer. In particular, Ets97D and Pnt depletion resulted in striking but opposite effects. Ets97D depletion boosted enhancer activity strongly, while Pnt depletion abolished it, as also double depletion of Ets97D and Pnt. Therefore, Ets97D appears to act as a repressor at *pst*_E1n that prevents the activator Pnt from inducing transcription by competition for the overlapping binding sites.

Pnt is known to function as a key TF activated by RTK signaling (Brunner et al. 1994; O’Neill et al. 1994; Shilo 2014). *Drosophila* RTKs (including EGFR, Heartless/FGFR, Breathless/FGFR, Torso, Sevenless) signal via the ras/MAP kinase cascade, resulting in activated MAPK, which phosphorylates the Pnt-P2 isoform. Thereby Pnt-P2’s function as a transcriptional activator is stimulated. Moreover, activated MAPK also phosphorylates the ETS family protein Aop/Yan and thereby inhibits Aop/Yan’s repressive Pnt-P2-antagonizing function. The role of Aop/Yan appears to have been taken over by Ets97D in case of *pst*_E1n. Ets97D (also named D-elg or Delg) has been proposed to function as a subunit of a TF complex homologous to mammalian NRF-2/GABP which has been implicated primarily in the regulation of housekeeping genes involved in ribosomal and mitochondrial biogenesis (Rosmarin et al. 2004). The proposal that *Drosophila* Ets97D corresponds to GABPα is supported by functional characterizations, which have also revealed an Ets97D requirement for the growth promotion of Cyclin D-Cdk4 (Baltzer et al. 2009; Frei et al. 2005), as well as for normal oogenesis (Schulz et al. 1993). The apparent growth-promoting role of Ets97D is intriguing in the context of temperature acclimation. Interestingly, Pnt has also been implicated very recently in the control of metabolic gene expression (Dobson et al. 2019).

Future work will be required for an identification of the cellular “thermometer” acting at the top of the pathways responsible for the increased activity of the *pst*_E1n CoolUp enhancer at low temperature. The endogenous *pst* gene is also expressed at higher levels at low temperature, as shown by our analysis of transcript and protein levels. Depletion of proteins crucial for *pst*_E1n reporter expression was found to have the same effects on *pst* transcript levels, although less pronounced and with some exceptions (*Socs36E* and *Pvr*). Thus, its transcriptional control might involve some additional regulatory inputs beyond those acting on *pst*_E1n.

The function of *pastrel* (*pst*) is not understood at the molecular level. The gene was named after one of Pavlov’s dogs because of an olfactory conditioning defect associated with a P element insertion in *pst* (Dubnau et al. 2003). Knockdown of *pst* in larval muscles impairs the formation of neuromuscular junctions (Fukui et al. 2012). More recently, *pst* function has also been linked with susceptibility to infection with *Drosophila* C Virus (DCV) (Magwire et al. 2012). The *pst* gene appears to be highly polymorphic in natural populations, and some of these polymorphisms are associated with either increased or decreased DCV susceptibility (Cao et al. 2017). Genes with sequence similarity to *pst* can be detected in insects but not in vertebrates. By analysing the Pst amino acid sequence using I-TASSER (Yang et al. 2015), we obtained evidence for structural similarity with Vps35, a subunit of the retromer complex, which is involved primarily in selection of transmembrane proteins for transport between endosome and trans Golgi network or plasma membrane. We have made an initial attempt to assess *pst* function in acclimation of *D. melanogaster* to cool temperatures by generating null alleles using CRISPR/cas. Somewhat surprisingly, these mutations did not prevent development into fertile adults and did not cause obvious developmental cold sensitivity. Clearly, additional work will be required to resolve whether and how *pst* contributes to acclimation to low temperature.

In conclusion, we are providing a rich data resource for future analyses of the transcriptional regulation of genes within the readily tolerated range in the ectotherm animal *D. melanogaster*. Moreover, we have developed a novel reporter assay that permits the characterization of the temperature dependence of the activity of CREs. Our identification and functional dissection of the *pst*_E1 enhancer demonstrates the utility of data resources and assay. Our results concerning the function of this CoolUp enhancer provides initial mechanistic insights into transcriptional upregulation induced by a shift to temperatures at the lower end of the readily tolerated range.

## Materials and Methods

### Cell culture

S2R+ cells (Yanagawa et al. 1998) and S2R+_*MtnA*p-*His2Av*-mRFP_hygro cells (Lidsky et al. 2013) were cultured in Schneider’s medium (Gibco, cat# 21720, Thermo Fisher Scientific, Waltham, MA), supplemented with 10% fetal bovine serum (Gibco, cat# 10500-064) and 1% Penicillin-Streptomycin (Gibco, cat# 15140). Cells were cultured at 25°C, unless otherwise noted. The number of live and dead cells was determined with an automated cell counter (Countess, Invitrogen, Thermo Fisher Scientific, Waltham, MA) after harvesting cells with 1 ml trypsin-EDTA (Gibco, cat# 25050-014) and staining with trypan blue (0.2% final concentration).

For the analysis of temperature effects on the proliferation of S2R+ cells, we plated aliquots of S2R+ cells (2 ml of suspension with 1×10^5^ cells/ml) in round cell culture dishes (35 mm diameter). Four aliquots were plated for each time point and temperature to be analyzed. Two of these aliquots were eventually used for cell counting and for the analysis of the cell cycle profile by flow cytometry. The other two aliquots were used for live imaging by phase contrast microscopy. All aliquots were first incubated for 24 hours at 25°C. Thereafter (t = 0), aliquots were shifted into incubators running at different temperatures (14, 18, 25 and 29°C) and incubated until analysis at the chosen time points. For experiment where temperature effects were analyzed only by phase contrast microscopy, cells were plated in 24 well plates, one plate for each temperature (9, 11, 13, 15 and 17°C) in three wells per plate.

The cell line HB10 was generated as described (Simcox 2013) by dissociation of embryos that were collected for two hours and aged for six hours at 25°C. The embryos were collected from a cross of *yw*; *P{UAS-Ras85D.V12}2*; *P{CaSpeR-CoStart_attP_STOP}32.1* males with *yw*; *P{w[+mC]=Act5C-GAL4}17bFO1*/*TM6B, Tb* virgins. Fly lines with *P{UAS-Ras85D.V12}2* (# 64196) and *P{w[+mC]=Act5C-GAL4}17bFO1* (# 3954) were obtained from the Bloomington *Drosophila* Stock Center (Indiana University, Bloomington, IN, USA). HB10 cells were grown in the same culture medium as the S2R+ cells and passaged once or twice each week. The HB10 cells were used for expression profiling after 31 passages.

S2R+ cell transfections were performed using FuGENE HD (Promega, cat# E2311) in 6 well plates or 25-cm^2^ flasks. In a 25-cm^2^ flask, 2.6×10^6^ cells were plated in 4 ml complete medium and incubated at 25°C. One hour after plating, 200 μl transfection mix (2 μg plasmid DNA and 8 μl FuGENE HD in Schneider’s medium) was added. To establish stably transfected cell lines, either 25 μg/ml blasticidin or 300 μg/ml hygromycin was added two days after transfection, unless otherwise noted. S2R+_EGFP-pst cells were generated by selection of stable integrations after transfection with pCaSpeR4-gEGFP-pst-blas^r^ (see below), and similarly, SR9 cells, which express *cas9* under control of the *MtnA* promoter, after transfection with pMT-cas9-hygro (see below). For induction of *cas9* expression, CuSO_4_ was added to the culture medium (final concentration 500 μM) 24 hours before transfection with gRNA and repair template plasmids (see below).

Single cell cloning was done with S2R+ feeder cells in 96 transwell co-culture plates (Corning, cat# CLS3380). First, 150 μl of a S2R+ cell suspension (3×10^5^ cells/ml) were plated in the bottom compartments below insert. Eighty μl of complete medium were then added into each insert. After 24 hours at 25°C, single cells were sorted by fluorescence-activated cell sorting (FACS) into the inserts. To minimize evaporation, plates were closed with parafilm and cultured in a plastic box with moistened tissue paper. When the feeder cells reached confluency, the upper insert plate was transferred onto a plate with fresh feeder cells prepared in an accessory plate (Corning, cat# CLS3382). Complete medium was added to the upper inserts every few days. When the clones had reached 100% confluence, cells were harvested by pipetting up and down. For expansion of the clonal population, cells were first transferred into a well of a 24-well plate and cultured in complete medium. Cloning efficiency after 8 weeks incubation was 40-50%. The characterization of the SR9rg clonal line is described in detail below.

Transfection of SR9rg for RMCE was performed in 6-well plates. 1×10^6^ cells were plated in 2 ml complete medium and incubated for 1 hour at 25°C before addition of 100 μl of transfection mix (500 ng exchange plasmid and 500 ng pCo-Bxb1_PhiC31, 4 μl FuGENE HD in Schneider’s medium). Whenever the transfected cells reached 100% confluence, they were passaged (1:4-5). Three weeks post-transfection, cells were collected and resuspended in complete medium at a concentration of 1×10^6^ cells/ml. Thereafter, aliquots of cells were plated in 6-well plates. The number of cells seeded was adapted to the temperature, to which they were shifted eventually (30 and 25°C: 1×10^6^ cells; 14°C: 2×10^6^ cells) in order to generate comparable cell densities at the time of cell harvesting. After seeding, the cells were cultured for an additional 24 hours at 25°C before shifting to either 14, 25 or 30°C. After 48 hours of incubation at these variable temperatures, cells were harvested and re-suspended in PBS (Gibco, cat# 10010-015). Cells were filtered with a Cell-Strainer (Falcon, cat# 352235) and kept on ice until analysis by flow cytometry.

Depletion of candidate transcription factors with predicted binding sites within *pst*_E1n was performed with SR9rg>*pst*_E1n (mRuby^−^, GFP^+^) cells. This cell population was obtained by FACS (see below), selecting cells with significant levels of green fluorescence and only background levels of red fluorescence from SR9rg>*pst*_E1n cells. After sorting by FACS, the selected cells were expanded at 25°C and frozen in aliquots. Depletion of the JNK protein kinase encoded by the *basket* (*bsk*) gene in S2R+ cells and treatment with bacterial lipopolysaccharides (LPS) were done as described previously (Radermacher et al. 2014).

### Flow cytometry and fluorescence activated cell sorting

Fluorescence activated cell sorting (FACS) and flow cytometry were carried out at the Cytometry Facility at the Irchel campus of the University of Zurich.

For determination of the cell cycle profiles of S2R+ cell populations after incubation at different temperatures, cells were harvested and fixed with 95% ethanol. Fixed cells were stored at 4°C (for maximally three weeks) before analysis by flow cytometry. Cells were resuspended in 1 ml PBS. To degrade RNA and stain DNA, 25 μl of RNase A stock solution (1 mg/ml) and 25 μl of propidium iodide stock solution (1 mg/ml, Fluka, cat# 81845) were added. After incubation at 4°C overnight, flow cytometric analysis of DNA content was completed using a BD FACSCanto instrument. PI was excited with a 561 nm laser and emission was detected using a 575/26 nm band pass filter. The resulting cell cycle histograms were analyzed with the software FlowJo (Treestrar Inc.).

For single cell cloning, cells were sorted with a FACSAria III cell sorter (BD Biosciences) using a 100 μm nozzle into 96 transwell co-culture plates. The same instrument and nozzle were also used for the isolation of SR9rg>*pst*_E1n (mRuby^−^, GFP^+^) cells into tube (Falcon, 352058) and plated in a 24-well plate for expansion of the population. Red fluorescence was excited with a 561 nm laser and emission was detected using a 610/20 nm band pass filter. Green fluorescence was excited with a 488 nm laser and emission was detected using a 530/30 nm band pass filter.

To determine enhancer activity after RMCE with SR9rg cells, we analyzed 1×10^5^ single cells for each sample using an LSR II Fortessa instrument (BD Biosciences). For quantification of the data obtained with *pst*_E1n and its derivatives, we used the software FlowJo. First, we defined a gate for the mRuby-negative population based on the data obtained with S2R+ cells (mRuby-negative) and SR9rg cells (mainly mRuby-positive cells). Second, we defined a gate for GFP-positive cells (gate 2) and a gate for GFP-negative cells (gate 3) based on the data observed after RMCE with *20×UAS* which do not contain GFP-positive cells. The median GFP signal intensity of the cells within gate 3 was used for background correction. It was subtracted from the median GFP signal intensity of the cells within gate 2, yielding the final value representing enhancer activity used for the bar diagrams.

### RNA interference

DNA templates for production of dsRNA by *in vitro* transcription were amplified enzymatically from genomic DNA of S2R+ cells with primers that introduce terminal T7 RNA polymerase promoter sequences (S10 Table). The DNA template fragments were purified from agarose gel using gel extraction kit (QIAGEN, cat# 28706) and used for *in vitro* transcription with the Ambion T7 Megascript Kit (Invitrogen, cat# AM1334). dsRNAs were precipitated by adding 3.3× volumes of 100% ethanol, followed by chilling overnight at −20°C. After a centrifugation (13100×g, for 15 minutes at 4°C), the supernatant was discarded. The retained pellet was washed with 75% ethanol, air-dried and dissolved in RNase-free water. The concentration of dsRNA in the final samples was determined using Nanodrop (Thermo Scientific, cat# ND-ONEC-W).

Depletion of candidate transcription factors involved in the function of the *pst*_E1n enhancer fragment was performed with SR9rg>*pst*_E1n (mRuby^−^, GFP^+^) and SR9rg>*ced6*_E (mRuby^−^, GFP^+^) cells. First, cells were harvested and resuspended in Schneider’s medium (without serum and other additives) at a density of 1.5×10^6^ cells/ml. 1.5 ml cell suspension were mixed with 15 μg dsRNA and seeded into a well of a 6-well plate. After an incubation of 45 minutes, 3 ml complete medium were added, followed by gentle mixing. After 4 days of incubation with dsRNA, cells were harvested and resuspended in complete medium at a concentration of 1×10^6^ cells/ml. Thereafter, aliquots of cells were plated in 6-well plates. The number of cells seeded into a well was adapted to the temperature, to which they were shifted eventually (30°C: 1×10^6^ cells; 25°C: 1×10^6^ cells; 14°C: 2×10^6^ cells). After seeding, the cells were cultured for an additional 24 hours at 25°C before shifting to either 14, 25 or 30°C. After 48 hours of incubation at these variable temperatures, cells were harvested and resuspended in PBS (Gibco, 10010-015). Cells were filtered with a Cell-Strainer (Falcon, 352235) and kept on ice until analysis by flow cytometry.

### Immunoblotting

Total cell extracts were prepared in 3x Laemmli buffer, heated for 5 min at 95°C and cleared by centrifugation at 4°C (3 min, 17000xg). Aliquots were frozen in liquid nitrogen and stored at −70°C. Protein concentration was determined with the Pierce 660nm Protein Assay (Thermo Fisher Scientific, cat# 2262). Proteins were resolved by standard SDS polyacrylamide gel electrophoresis using pre-stained PAGE RULER plus (Thermo Fisher Scientific, cat# 26619) molecular weight markers. Proteins were transferred onto nitrocellulose membranes by electrotransfer with a tank system. Membranes were transiently stained with Ponceau S to confirm successful transfer. After blocking, membranes were probed with the following primary antibodies: rabbit anti-JNK (Santa Cruz Biotechnology, Inc., Heidelberg, Germany, cat# sc-571, 1:1000), rabbit anti-phospho-JNK 81E11 (Cell Signaling Technology, Leiden, Netherlands, cat# 4688, 1:1000), rabbit anti-phospho-p38MAPK (Cell Signaling Technology, cat# 9211, 1:400), mouse anti-alpha-tubulin DM1A (Sigma Aldrich Chemie GmbH, Buchs, Switzerland, cat# T9026, 1:50’000), mouse anti-PSTAIR (Sigma Aldrich Chemie GmbH, cat# P7962, 1:50’000), mouse anti-FLAG M2 (Sigma, cat# F1804, 1:1000) and rabbit anti-mCherry (1:1000) (Herzog et al. 2013). As secondary antibodies we used horseradish peroxidase conjugated goat IgG anti-rabbit and anti-mouse IgG (H+L) (Jackson ImmunoResearch Europe Ltd, Cambridge, UK, cat# 111-035-003 and 115-035-003, 1:1000). Signals were detected by chemiluminescence. Signals corrected by subtraction of local background were quantified using ImageJ and compared based on linear interpolation of intensity values obtained with the dilution series that was resolved in parallel.

### qRT-PCR

Total RNA was extracted from cultured cells with TRIzol (Invitrogen, cat# 15596026), followed by DNase digestion (DNA-free DNA Removal Kit, Ambion, cat# AM1906). Alternatively, we also used Direct-zol™ RNA MiniPrep Plus kit (ZYMO, cat# R2070, Lucerna-Chem AG, Lucerne, Switzerland) for RNA preparation. cDNA synthesis was performed using Transcriptor High-Fidelity cDNA Synthesis Kit (Roche) with 500 ng RNA per reaction. Quantitative real-time PCR was performed using SYBR Green with an Applied Biosystems 7900HT using the recommended two-step cycling protocol. Alternatively, the QuantStudio™ 3 Real-Time PCR System (ThermoFisher, cat# A28137) was used. Primer sequences are given in S10 Table. For normalization, we used primer pairs for three genes, *Act5C*, *alphaTub84B* and *Tbp*.

### Plasmid constructions

All PCRs for cloning were performed with Phusion High Fidelity DNA polymerase (NEB, cat# M0530) using the default buffer (NEB, cat# B0518S).

For generation of pMT-cas9-hygro^r^, we first introduced a ClaI restriction site into pMT-hygro^r^ (Invitrogen, Carlsbad, CA, USA) by digesting with BglII and XbaI, followed by ligation with a double-stranded DNA oligonucleotide (ds oligo) that was obtained by annealing the oligos YB007 and YB008 (see S10 Table). After digestion of the resulting intermediate with ClaI and XbaI, a fragment containing the 3×Flag-nls-cas9-nls coding sequence isolated from pBS-Hsp70-cas9 (Addgene, #46294) with same enzymes was inserted, yielding the final plasmid, which was verified by sequencing the insert region.

For chromosomal integration of the RMCE target region in SR9 cells by CRISPR/cas9, we generated pCFD3:U6:3_attP40, a derivative of pCFD3:U6:3gRNA (Addgene, #49410) for expression of a single guide RNA. The sgRNA was designed for targeting the “attP40” locus within a facultative *Msp300* intron on the left arm of chromosome 2. This region is known to host the recombinant transposon P{CaryP}attP40 that one of the most frequently used attP landing sites for PhiC31-mediated generation of transgenic *Drosophila* lines, including thousands of UASt-RNAi lines with limited basal and high GAL4-mediated expression (Markstein et al. 2008). The expression of flanking genes within a 40 kb range was found to be independent of temperature in S2R+ cells. pCFD3:U6:3gRNA was digested with BbsI and a ds oligo obtained by annealing YB109 and YB110 was inserted. The insert regions was verified by sequencing.

The plasmid pUC57-RMCE-target was generated for chromosomal integration of the target region required for recombinase-mediated directional cassette exchange. A synthetic EcoRI-HindIII DNA fragment inserted into the corresponding restriction sites of pUC57 was obtained from GenScript (Leiden, Netherlands). The insert DNA of this intermediate 1 contained the minimal attB target sites for the PhiC31 and Bxb1 integrases as well as the mEGFP coding sequence. An mRuby2 marker gene was generated and inserted between the attB target sites of intermediate 1 as follows. The mRuby2 coding sequence was amplified from pCDNA3_mRuby2 (Addgene, # 40260) using JB007 and JB008. The resulting PCR fragment was digested with EcoRI and XbaI, and inserted into the corresponding sites of pUASt downstream of the minimal promoter and the 5’ UTR of *Hsp70*. The resulting *Hsp70*P-mRuby2 cassette was then amplified from the pUASt-mRuby2 construct using JB005 and JB006. The PCR fragment was digested with NcoI and AvrII, and inserted into the corresponding sites of intermediate 1. The resulting intermediate 2 was complemented with a *copia* promoter-blas^r^ cassette after amplifying the cassette using JB011 and JB012 from a plasmid with a corresponding synthetic DNA insert obtained from GenScript. The resulting PCR fragment was digested with NcoI and SpeI, and inserted into the corresponding sites of intermediate 2, yielding intermediate 3. Finally, flanking homology arms for targeting to the attP40 region were added. The left homology region (HRl, 938 bp) was amplified from S2R+ genomic DNA using JB003 and JB004. The resulting PCR fragment was digested with BglII and AflII, and inserted into the corresponding sites of intermediate 3, yielding intermediate 4. The fragment with the right homology region (HRr, 573 bp), also amplified from S2R+ genomic DNA with JB009 and JB010, was digested with XhoI and HindIII, and inserted into the corresponding sites of intermediate 4 to arrive at the final construct pUC57-RMCE-target.

For the generation of the dual integrase expression plasmid pCo-Bxb1_PhiC31, we first deleted the blas^r^ coding region from a precursor plasmid (pCoBlast_mitoKillerRed) using a one-primer method (Makarova et al. 2000) with primer JB020. The presence of the deletion was confirmed by sequencing. The region coding for PhiC31 was amplified from a template plasmid (pHsp70_phiC31-NLS_SV40; kindly provided by J. Bischof and K. Basler, University of Zurich, Zurich, Switzerland) using JB021 and JB022. The resulting PCR fragment was digested with EcoRI and XbaI, followed by ligation into the corresponding sites of the modified precursor plasmid, yielding pCo_PhiC31. The region coding for Bxb1 was amplified from the plasmid pET11_Bxb1 (Huang et al. 2011) using oEC134 and oEC135. The resulting PCR fragment was digested with AccIII and ligated into AgeI digested pCo_PhiC31, yielding the final construct pCo-Bxb1_PhiC31. In this product, a *copia* promoter fragment is upstream of the Bxb1 coding sequence. Moreover, the PhiC31 coding region is further upstream of the *copia* promoter fragment in reverse orientation with a minimal *Hsp70* promoter at its start. As a result, the two promoters (*copia* and *Hsp70*) drive bidirectional expression of the integrases, stimulated by transcription factors recruited by the *copia* promoter fragment. The correctness of the plasmid was confirmed by sequencing.

As a vector for the production of exchange plasmids to be used for RMCE in SR9rg cells, we designed pUC57-attP_P_-mcs-DSCP-nls-attP_B_. The insert region in this pUC57 derivative has the attP sequences for the PhiC31 and Bxb1 integrases on the left and right ends, respectively. Moreover, it contains a multiple cloning site (mcs) and the *Drosophila* synthetic core promoter (DSCP) (Pfeiffer et al. 2008) followed by a translational start codon and sequences coding for a nuclear localization signal (nls). The plasmid was obtained from GenScript and its insert region (EC358) is given in S10 Table. Using a one-primer method (Makarova et al. 2000) with YB317, a derivative of pUC57-attP_P_-mcs-DSCP-nls-attP_B_ was made lacking the DSCP. This pUC57-attP_P_-mcs-attP_B_ vector was used for assaying candidate CREs together with their linked endogenous promoter instead of relying on the DSCP.

Exchange plasmids with candidate CRE fragments for analysis after RMCE in SR9rg cells were generated using either pUC57-attP_P_-mcs-DSCP-nls-attP_B_ or pUC57-attP_P_-mcs-attP_B_. Candidate CREs were amplified by PCR from S2R+ cell genomic DNA (for primers, see S10 Table). Alternatively, synthetic gene blocks (S10 Table) were used. Some of the resulting exchange plasmids were further modified. To generate the exchange plasmids containing the fragments *pst*_E1b, *pst*_E1m, *pst*_E1n and *pst*_E1o, we introduced deletions into the exchange plasmid containing *pst*_E1 using a one-primer method (Makarova et al. 2000) and either YB440, YB451, YB453, or YB439. This method and YB316 were also used to delete the DSCP from the exchange plasmid containing the *Lip4*_E fragment.

The plasmid pgEGFP-pst-blas^r^ was used for transfection of S2R+ cells and selection of random chromosomal integrations. It was obtained by modifying the precursor plasmid pCaSpeR4-gEGFP-pst, which was generated as follows. In a first step, the multiple cloning site of the pCaSpeR4 vector was modified to include an AvrII site. Thus, pCaSpeR4 was digested with BglII and XbaI, followed by ligation with a ds oligo obtained by annealing CL328 and CL329. Using the EcoRI and NotI in the resulting intermediate 1, we inserted a fragment encompassing the 5’ region of the *pst* locus after enzymatic amplification with CL330 and CL331 from *w*^1^ genomic DNA and digestions with EcoRI and NotI, yielding intermediate 2. A fragment encompassing the middle region of *pst* was amplified with CL332 and CL333 from *w*^1^ genomic DNA, digested with NotI and AvrII, and ligated into corresponding restriction sites of intermediate 2. The resulting intermediate 3 was modified by insertion of a fragment encompassing the 3’ region of *pst*, which was amplified with CL334 and CL335 from *w*^1^ genomic DNA. The PCR fragment as well as intermediate 3 were digested with AvrII and XbaI before ligation into intermediate 4. For insertion of the EGFP coding region, we used enzymatic amplification of this sequence with CL336 and SCH21, and inserted it into intermediate 4 using NotI to arrive at pCaspeR4-g-EGFP-pst. The *pst* gene region with an N-terminal EGFP insertion was then cut out from pCaspeR4-g-EGFP-pst using AfeI and XbaI. The resulting insert fragment was ligated with a vector fragment derived from pCoBlast. The vector fragment was made by digestion of pCoBlast with HindIII. The resulting overhangs were filled with Klenow fragment before digestion with a second restriction enzyme, XbaI. The ligation of insert and vector fragment yielded pg-EGFP-pst-blas^r^.

For the generation of *pst* mutant fly lines by CRIPSR/cas9, we generated derivatives of pCFD5:U6:3-t::gRNA (Addgene, #73914). A first derivative (pCFD5:U6:3-t::gRNA_pst-1) was made with the help of the gene block YB392 (Integrated DNA Technologies, Leuven, Belgium). The resulting plasmid allowed expression of two distinct gRNA targeting the *pst* coding region. An analogous construct (pCFD5:U6:3-t::gRNA_pst-2) for expression of another gRNA pair was made with gene block YB393. The gene blocks were digested with BbsI and ligated into the corresponding sites of pCFD5:U6:3-t::gRNA.

### Characterization of the clonal SR9rg cell line

The SR9rg cells, which we generated for analysis of temperature effects on CREs after RMCE, were derived from SR9 cells. Expression of *cas9* was induced in SR9 cells for 24 hours by addition of CuSO_4_ (final concentration 500 μM) before co-transfection with the gRNA plasmid pCFD3:U6:3_attP40 and the repair template plasmids pUC57-RMCE-target. Blasticidin was added to the culture (final concentration 25 μg/ml) for selection of stable integration events. Microscopic analysis of the blasticidin-resistant cell population revealed mRuby expression as expected. By single cell cloning, we established 62 clonal lines. From seven of these, we were able to amplify a PCR fragment with a size indicating the presence of functional RMCE exchange target regions. Moreover, preliminary evidence from the initial PCR assays was consistent with on-target integrations in the “attP40” chromosomal region. However, these initial PCR analyses also indicated the presence of additional off-target integrations of sequences derived from pUC57-RMCE-target. One of the seven clonal cell lines, SR9rg, was chosen for evaluation of the efficiency of RMCE after co-transfection with the dual integrase expression plasmid pCo-Bxb1_PhiC31 and an exchange plasmid derived from pUC57-attP_P_-mcs-DSCP-nls-attP_B_. As these experiments indicated that SR9rg cells permit successful RMCE, we continued to use this cell line. In parallel, we analyzed the SR9rg cells in further detail, including whole genome sequencing (WGS) (see below). These analyses failed to confirm the presence of an on-target integration of the pUC57-RMCE-target sequences flanked by the left and right homology arms. Moreover, they confirmed the presence of off-target integrations of pUC57-RMCE-target sequences. Using the WGS data for read count analyses, sequences derived from pUC57-RMCE-target appear to be present in the SR9rg genome amounting to about 10 copies of the plasmid. By analysis of paired end reads, where one read maps to the *Drosophila* reference genome and the other to the pUC57-RMCE-target plasmid, sequences derived from this plasmid appear to be integrated in at least 34 distinct genomic locations. The analysis of such chimeric paired reads also suggested that several of these integrations have structures that cannot support an RMCE event, which results in a switch from mRuby to EGFP expression. The apparent structural complexity of the off-target integrations and the aneuploid genome of S2R+ cells precluded a comprehensive identification and structural clarification of all the integration events. We point out that we observed the same high propensity for undesired off-target integrations with S2R+ cells during several additional attempts to generate cell lines with single on target integration events by similar CRISPR/cas strategies. However, in SR9rg cells we also identified one clearly functional integration on chromosome 3L that could be demonstrated to undergo the expected RMCE after cotransfection with the dual integrase expression plasmid pCo-Bxb1_PhiC31 and an exchange plasmid derived from pUC57-attP_P_-mcs-DSCP-nls-attP_B_ (S14 Fig.). The FACS analysis of SR9rg derived cell populations obtained after RMCE with an exchange plasmid containing a strong enhancer suggested that the identified functional target region is not the only one, since we observed a fraction of cells expressing both red and green fluorescence beyond those expressing green but not red fluorescence (see for example Fig. 4B). The cells expressing both red and green fluorescence have presumably undergone RMCE at least at one but not at all functional target regions. The FACS analyses of SR9rg derived cell populations after RMCE with a strong enhancer also indicated that the number of functional target regions permitting the generation of a reporter gene expressing GFP by RMCE is either rather low, or that RMCE occurs usually with very high efficiency at almost all the functional regions. Otherwise, the population of cells expressing green but not red fluorescence after RMCE should have been exceeding low, in contrast to our observations (see for example Fig. 4B).

For confirmation of presence and function of the RMCE target region integrated within the *CG13288* locus in chr3L according to our WGS analysis (S14 Fig.), we performed PCR assays using genomic DNA from SR9rg and SR9rg>*pst*_E1n cells and the primers oEC264, oEC267, oEC263, EC265 and YB654 (S10 Table).

### Expression profiling with DNA microarrays

For the analysis of the S2R+ cell transcriptomes at distinct temperatures (11, 14, 25 and 30°C) (experiment M1), cells were seeded in 60 mm culture dishes in 2.5 ml medium in numbers adjusted to the temperature, to which they were shifted eventually (30°C: 1.5×10^6^; 25°C: 1.7×10^6^; 14°C: 3.5×10^6^; 11°C: 4.5×10^6^). One day after seeding, cells were shifted to different temperatures (11, 14, 25 or 30°C) for 24 hours before isolation of total RNA. Three biological replicates were performed (without prior computation of an appropriate sample size). RNA was isolated with the NucleoSpin RNAII kit (Macherey-Nagel, Oensingen, Switzerland).

For analysis of the effects of *His2Av* depletion on the temperature dependence of the S2R+ cell transcriptome (experiment M2), aliquots of cells were also seeded into 60 mm culture dishes in adjusted numbers (30°C: 0.4×10^6^; 25°C: 0.65×10^6^; 14°C: 0.7×10^6^). Twenty-four hours after seeding, we added 30 μg dsRNA per dish, derived from either *His2Av* or *lacZ* for control. Four days after dsRNA addition, aliquots were shifted to different temperatures (14, 25 and 30°C) for 24 hours, followed by isolation of total RNA using the NucleoSpin RNAII kit. Three replicates were processed (without prior computation of an appropriate sample size).

For the time course analysis of the S2R+ cell transcriptome after a temperature downshift to 14°C (experiment M3), we seeded aliquots of 1.4x×10^6^ cells in 2.5 ml of medium in 60 mm culture dishes. After 36 hours of incubation at 25°C, culture aliquots were shifted to 14°C. Total RNA was isolated at different times after the downshift (0, 4, 12, 24 and 72 hours). In addition, instead of shifting to 14°C, control aliquots were grown for an additional 12 hours at 25°C before isolation of total RNA. At least three biological replicates for each time point were generated (without prior computation of an appropriate sample size).

For analysis of a temperature downshift to 14°C on the transcriptome of HB10 cells (experiment M4), aliquots of 2.6 ×10^6^ cells were seed into 35 mm culture dishes. After 36 hours of incubation at 24°C, half of the aliquots were shifted to 14°C while the other half was maintained at 24°C. Total RNA was isolated 24 hours later. Three biological replicates were processed (without prior computation of an appropriate sample size). In case of the time course and HB10 cell experiments, total RNA was isolated using TRIzol (Invitrogen, cat# 15596026).

Before probe generation for DNA microarray hybridization, the total RNA preparations were further cleaned. Potential contaminations with DNA were eliminated by DNase I treatment, followed by RNA purification with the RNeasy MiniKit (Qiagen). The concentration of the resulting RNA preparations was determined with a NanoDrop 1000 Spectrophotometer V3.7 and RNA integrity was confirmed with an Agilent 2100 Bioanalyzer.

For generation of Cyanine 3 (Cy3)-labelled complementary RNA (cRNA) probes, we used the Low-Input QuickAmp kit (Agilent, Santa Clara, CA, USA). The resulting cRNA was purified using the Absolute RNA Nanoprep kit (Agilent). Successful Cy3 incorporation was confirmed by NanoDrop measurements and cRNA integrity with the Bioanalyzer. The hybridization mix (55 μl) contained 1.65 μg of fragmented cRNA, 2.2 μl of 25× hybridization buffer, 11 μl of 10× blocking agent and RNase-free water. For hybridization, we used single-color gene-expression microarrays purchased from Agilent. Two experiments (M1 and M2) were completed with *Drosophila* Gene Expression Microarray 4×44K (# G2519F, design ID: 018972) and two experiments (M3 and M4) with *Drosophila* Gene Expression Microarray (V2) 4×44k (#G2519F, design ID: 021791). After microarray hybridization and washing, signal intensities were recorded with Agilent Feature Extraction Software. Raw data obtained in experiments M1-4 have been deposited in NCBI’s Gene Expression Omnibus (Edgar et al. 2002) and are accessible through GEO Series accession number GSE159174 (https://www.ncbi.nlm.nih.gov/geo/query/acc.cgi?acc=GSE159174).

In case of experiment M1, raw data was processed with GeneSpringGX11.0 (Agilent). Inter array differences were corrected using the option “median scaling to a control sample” of this software and the three replicates obtained after incubation at 25°C as control samples. For comparison of the temperature dependence of signal intensities observed with a given microarray probe oligo with the signal intensities detected by other probes, a baseline transformation was applied so that all log_2_ values of signal intensities observed in the different replicates and temperature conditions with a given probe that a median = 0. Probes with low or incoherent values were filtered out before further analysis. Values were discarded, if the flag “present” was not specified in at least two of the three replicates obtained for a given temperature. Moreover, in case of the analyses concerning temperature-dependence of central cellular processes (Fig. 3A), probes with an expression level below 100 were not considered.

In case of the experiments M2-4, raw data was processed using R. For interarray comparisons, quantile normalization as implemented in the Bioconductor package preprocessCore (Bolstad B. 2020. DOI: 10.18129/B9.bioc.preprocessCore) was applied in case of experiment M2. In case of the experiments M3 and M4, we used the limma package (Smyth 2005). Signal intensities were background-corrected with normexp convolution method and quantile-normalized. Signal intensities were further adjusted with the R surrogate variable analysis (SVA) package (Leek and Storey 2007) for removal of batch effects. We computed the median of signal intensities among technical replicate probes present more than once on the microarray. Moreover, for further analyses in case of experiment M3, we only retained probes that had log_2_ signal intensities higher than 6.6 on at least three of all the 23 analyzed microarrays. To identify probes with significant temporal change in signal intensities, we used an empirical Bayes extension of analysis of variance (ANOVA) as implemented in the limma package. Importantly, samples taken at t0 and at t12 after incubation at the control temperature 25°C (ct12) were grouped and considered as reference baseline group for this statistical test. The t0 and ct12 transcriptomes were actually found to be very highly correlated. For further analysis, we selected all the probes with an FDR-corrected p-value < 0.05 and with a fold change in expression level ≥ 2 at one or more time points compared to the baseline value. To cluster probes with similar temporal expression profiles (Fig. 3E), we used computed Spearman distance metric among signal intensities and run a *k*-means cluster (R stats package). The elbow method (Goutte et al. 1999) was used for assessing the number of clusters in our data set. Probes associated with systematic gene names were annotated with the R biomaRt library.

For the analyses concerning temperature-dependence of central cellular processes (Fig. 3A, 3F), we first generated curated lists of genes with unequivocal association to one of the selected functional networks or to an organelle (S phase, M phase, autophagy, proteasome, amino acid catabolism, insulin/TOR signaling, cytoplasmic ribosome, mitochondrial ribosomes, oxidative phosphorylation, glycolysis, TCA cycle, beta oxidation, lipid metabolism, and peroxisome). To generate these gene lists (S11 Table), *D. melanogaster* genes were first filtered using corresponding GO terms. The resulting list were manually curated and further complemented using information from the KEGG database, as well as from various publications, including (Baker and Thummel 2007). All probes annotated to the selected genes were identified and probes with significant expression were used for quantitative analyses. All the additional analysis of functional associations of differentially expressed genes (Fig. 3B) were performed using the option “Proteins with Values/Ranks” of String (v11.0) (Szklarczyk et al. 2019).

For the analysis of temperature effects on genes known to be regulated during response to different types of cellular stress, we first generated lists of stress-regulated genes primarily based on the results of (Girardot et al. 2004) and additional publications cited by these authors (Gregorio et al. 2001; Landis et al. 2004; Zinke et al. 2002). These publications describe microarray analyses of the transcriptome response in adult flies or larvae after exposure to various stressors (oxidative stress: paraquat, hydrogen peroxide, hyperoxia; ER stress: tunicamycin; starvation; bacterial or fungal infection). For each type of stress, genes induced more than 1.5 fold (or 2 fold in case of paraquat) were selected for our analysis. All probes detecting transcripts derived from these stress genes were identified (S11 Table) and the signals observed with these probes in our microarray experiments were extracted for the analysis of temperature on stress gene expression in S2R+ cells.

### Expression profiling with 3’ RNA-Seq

S2R+ cells were plated into 60 mm dishes. The number of plated cells was adjusted according to the temperature to be analyzed eventually (30°C: 1.5×10^6^ cells; 25°C: 1.7×10^6^ cells; 14°C: 3.5×10^6^ cells; 11°C: 4.5×10^6^ cells). Three biological replicates were prepared for each temperature (without prior computation of an appropriate sample size). All aliquots were first cultured at 25°C for 24 hours after plating, before shifting to either 11, 14, 25 or 30°C.

For the analyses with adult males, *w*^1^ flies were used to start cultures in bottles at 25°C. Newly eclosed adult male flies (0-12 hours after eclosion) were collected and distributed into aliquots into fresh food vials. Three biological replicates were prepared for each temperature. The aliquots were then shifted to either 11, 14, 25 or 30°C.

After 24 hours of incubation at the different target temperatures, total RNA was extracted from the S2R+ cells and the male flies using TRIzol (Invitrogen, cat# 15596026), followed by DNase digestion (DNA-free DNA Removal Kit, Ambion, cat# AM1906). Libraries were generated with the QuantSeq 3’ mRNA-Seq Kit REV for Illumina (Lexogen) and sequenced by the Functional Genomics Center Zürich (FGCZ, University and ETH Zürich). 2×150 bp paired-end sequencing was performed with HiSeq4000 (llumina, San Diego, CA).

Data analysis was completed as follows. To identify genes differentially expressed at the different incubation temperatures, sequencing data from the 24 samples (12 samples from S2R+ cells and 12 samples from adult males, 419.5 million reads in total) were aligned to the *Drosophila melanogaster* (dm6) reference genome (Ensembl version 98) with STAR aligner (Dobin et al., 2013). The STAR dm6 index was built with adjusted parameters (--genomeSAindexNbases 10) to accommodate the size of the genome. Moreover, the gene annotation was incorporated to allow correct mapping of spliced reads. After alignment, count tables of reads mapping to genes across all the 24 samples were obtained using htseq-count (Anders et al. 2015). Normalization was computed using the cpm function of edgeR (Robinson et al. 2010). Two comparisons (Cells 11°C+14°C vs. Cells 25°C+30°C and Flies 11°C+14°C vs. Flies 25°C+30°C) were made using edgeR. Weakly expressed genes were filtered out using HTSFilter (Rau et al. 2013).

To calculate the correlation between the results of 3’ RNA-Seq and DNA microarray analyses, log_2_ values of fold change of the expression level of a given gene at high temperature (25 and 30°C values pooled) compared to low temperature (11 and 14°C values pooled) was evaluated using Pearson’s method.

For analysis of alternative polyadenylation, a poly(A) site (PAS) database was constructed by processing the pooled 3’ RNA-Seq reads from all samples in a first step. After read-mapping to the reference genome (dm6), the position of the last nucleotide before the start of the poly(A) tail in given read was determined for each read. This position represents the cleavage and polyadenylation site. The detected PAS positions were assigned to clusters using a window size from 25 nucleotides upstream to 25 nucleotides downstream of the actual PAS. This clustering accounts for the fact that cleavage and polyadenylation do not occur with single nucleotide precision. Since the REV version of the Lexogen QuantSeq 3’ mRNA-Seq Kit was used, we filtered out positions in the PAS database that corresponded to internal priming (IP) events rather than true polyadenylation. To this end, the genomic sequence in the region from −30 to 10 around each putative PAS was checked and all sites which contained seven or more consecutive A nucleotides in this region were removed from the PAS database. Moreover, putative PAS were also removed if the region from −30 to 0 did not contain any of the known polyadenylation signals (Gruber et al. 2016). The filtered PAS in the database were assigned to genes. This was accomplished after extending the 3’ UTR region annotated in FlyBase to include an additional 5000 nucleotides or up to the middle of the distance between the PAS and the start of the downstream gene. This extension was motivated by the observation that 3’ UTR annotations are overly stringent. The database with all the filtered and assigned PASs was used for the generation of count tables. For each condition (cells or flies at either 11, 14, 25 or 30°C, respectively) the read counts observed at the PASs in the database were determined. Normalized values were computed using the cpm function of edgeR. To identify genes with APA switches, we used DESeq (Anders et al. 2012) for the analysis of two comparisons (Cells 11°C+14°C vs. Cells 25°C+30°C, Flies 11°C+14°C vs. Flies 25°C+30°C). For each gene, we considered all possible poly(A) site pairs and selected only those pairs that have opposing direction (fold-change) of regulation in a particular comparison with FDR < 0.05. Among the retained pairs, the one with the greatest fold-change distance was ultimately chosen and labeled as enhanced (fold-change (proximal/distal) > 0) or repressed (fold-change (proximal/distal) < 0). Thereby we obtained lists of genes with temperature-dependent differential APA, which contained for each of these genes the two PASs that were most divergently used in a particular comparison.

For generation of correlation plots (Fig. 2A, E, 3D, S7 Fig.) and volcano plots (Fig. 2B, E; S7 Fig), data for expression levels after normalization and filtering to eliminate genes with spurious expression was processed with the R packages corrplot (Taiyun Wei and Viliam Simko (2017). R package "corrplot": Visualization of a Correlation Matrix (Version 0.84), https://github.com/taiyun/corrplot) and EnhancedVolcano (Blighe K, Rana S, Lewis M (2020). EnhancedVolcano: Publication-ready volcano plots with enhanced colouring and labeling. R package version 1.6.0, https://github.com/kevinblighe/EnhancedVolcano).

The 3’ RNA-Seq data have been deposited in NCBI’s Gene Expression Omnibus (Edgar et al. 2002) and are accessible through GEO Series accession number GSE159174 (https://www.ncbi.nlm.nih.gov/geo/query/acc.cgi?acc=GSE159174) and also at http://www.expressrna.org/index.html?action=library&library_id=20170831_yu.

### ATAC-Seq

Aliquots of 3.5 million S2R+ cells were seeded in round 60 mm dishes in case of the samples that were eventually exposed to 25 or 29°C. In case of the samples eventually exposed to 14°C, the starting number of cells was 7 millions. Four replicate cultures were prepared for each analyzed temperature (without prior computation of an appropriate sample size). After a 36 hour incubation at 25°C, the four aliquots were shifted to either 14, 25 or 29°C. Cells were harvested 24 hours after this temperature shift. One aliquot for each temperature was harvested first at room temperature for cell count determination. Cell harvest in case of all the remaining aliquots was performed quickly using 0.025% Trypsin-EDTA (ThermoFisher Scientific, 25200-056) within the incubators and at the temperatures used during the preceding 24 hour incubation, with solutions pre-equilibrated to the different incubation temperatures. The culture medium was removed and cells were rinsed once with PBS before addition of 1 ml Trypsin-EDTA solution for 5 minutes. The released cells were transferred into a 15 ml Falcon tube and combined with four rinses of the plates (each with 1 ml of complete medium). The resulting master cell suspension was diluted into ice-cold complete medium in order to obtain an Eppendorf tube with 75000 cells in 1 ml for each aliquot. Cells were sedimented (500×g for 5 min, 4°C) and washed once with 50 μl PBS. Cells were resuspended and lysed with 50 μl of cold lysis buffer (10 mM Tris-HCl, 10 mM NaCl, 3 mM MgCl_2_, 0.1% NP-40). Lysed cells were sedimented (500×g for 10 min, 4°C). Tagmentation was performed during 30 minutes at 25°C in a 50 μl reaction in TD Buffer (Illumina, FC-121-1030) containing 2.5 μl Tn5 Transposes (Illumina, FC-121-1030) (Buenrostro et al. 2013). Tagmented DNA was purified using a Qiagen MinElute Kit and amplified as described (Buenrostro et al. 2013) using a Nextera forward primer (Ad1_noMX, S10 Table) in combination with a replicate-specific bar-coded reverse primer (Ad2.1 to Ad2.9, S10 Table). Libraries were purified using Qiagen PCR Cleanup Kit. Samples were pooled and sequenced at the FGCZ using HiSeq2500 v4 providing 126 bp paired end reads. The nine ATAC-Seq libraries were sequenced to a median depth of 52 million reads per sample.

The sequence data was further processed as follows. Sequencing adaptors were trimmed with Trimmomatic (version 0.38) (Bolger et al. 2014). All reads were mapped to *Drosophila* genome dm6 using Bowtie2 (Langmead et al. 2019) with the parameters -X 2000 --very-sensitive. Non-mapped, non-unique, and mitochondrial reads were filtered out with SAMtools (Li et al. 2009; Li 2011). PCR duplicates were marked and removed with Picard (https://broadinstitute.github.io/picard/). Subsequently, all reads were adjusted for the binding footprint of Tn5 transposase, i.e., all reads aligning to the top strand were shifted by four bp and all reads aligning to bottom strand were shifted five bp in 5’ direction.

ATAC-Seq peaks were called with MACS2 (Zhang et al. 2008) for each library with parameters -f BEDPE --keep-dup all -q 0.01 -g dm --nomodel. Differential peaks were analyzed with DiffBind (Stark and Brown, 2011). Read counts were obtained with the dba.count function. To keep the peaks at a consistent width, peaks were re-centered around the point of greatest enrichment with “summits = 250”, yielding 17175 consensus peaks among all nine libraries. Next, DESeq2 (Love et al. 2014) was employed for count normalization and differential analysis using default threshold of FDR <= 0.05. Normalized peaks with low reproducibility among replicates were eliminated as described (Causton et al. 2003). The ATAC-Seq data have been deposited in NCBI’s Gene Expression Omnibus (Edgar et al. 2002) and are accessible through GEO Series accession number GSE159174 (https://www.ncbi.nlm.nih.gov/geo/query/acc.cgi?acc=GSE159174).

### Whole genome sequencing

For the characterization of SR9 derived cell lines by whole genome sequencing, we selected seven lines, including SR9rg, obtained by single cell cloning from the cell population resulting after SR9 cell transfection with pCFD3:U6:3_attP40 and pUC57-RMCE-target. In addition, we selected three additional clonal cell lines that were obtained after transfection of SR9 cells with pCFD3:U6:3_attP40 and distinct repair template plasmids. The selected cell lines were cultured in 25-cm^2^ flasks at 25°C. DNA was isolated using QIAamp DNA Mini Kit (Qiagen, 51306). Libraries were prepared with TruSeq Nano DNA library Prep kit and sequenced at the FGCZ. 2×150 bp paired-end sequencing was performed with HiSeq4000.

Preqc (Simpson 2014) was used to estimate sequence coverage, per-base error rates, as well as genome size, heterozygosity and repeat content. De novo assembly of the paired-end sequencing data was performed using SPAdes (Bankevich et al. 2012) with default parameters. Additionally, AByss (Simpson et al. 2009) was also used for de novo assembly with SPAdes error corrected reads. QUAST (Gurevich et al. 2013) analysis showed that these two algorithms produced similar results in terms of the number and length of contigs and scaffolds identified in each sample. Assembly quality of SPAdes and Abyss was determined using QUAST.

We built three in-silico reference genomes, which consisted of the *Drosophila* reference genome sequence (dmel_r6.19) plus the plasmid sequence of pMT-cas9-hygro^r^ and of one of the three distinct repair plasmids (including pUC57-RMCE-target). Paired-end reads were aligned to in-silico reference genome using BWA aligner (Li and Durbin 2009), duplicates were marked by SAMBLASTER (Faust and Hall 2014) and removed. The in-silico genomes were then used as reference genomes for a search of hybrid structure variants (hSvs) linking dmel_r6.19 sequences with sequences in the plasmids. To do so, we used two algorithms LUMPY (Layer et al. 2014) and Delly (Rausch et al. 2012). In the case of LUMPY analyses, reads with mapping quality either above Q20 or Q30 were used. In the case of Delly analyses, either all reads or reads with mapping quality above Q30 were used. Paired read analysis for the identification and mapping of the chimeric pairs, where on read was aligned to the *Drosophila* reference genome and the other read to the plasmid sequences, was performed by homemade scripts written in Python. Copy number variation was analyzed by CNVnator (Abyzov et al. 2011).

### Generation of pst mutant fly lines

CRISPR/cas9 was used for the generation of fly lines with mutations in *pst*. The plasmids allowing expression of a pair of gRNAs (see above) were injected into eggs collected from *yw*; *attP40*{*nos-cas9}/*CyO flies, respectively (BestGene Inc, Chino Hills, CA, USA). The males that eclosed from the injected eggs were crossed individually with virgins *w*\*; *Sb*/TM3, *Ser*. Some of the F1 progeny resulting from these crosses were analyzed with a PCR assay, involving amplification from genomic DNA with the primers YB444 and YB445 (after injection of gRNA plasmid 1) or YB446 and YB447 (after injection of gRNA plasmid 2), which anneal to sites flanking the gRNA target sites. Therefore, shortened PCR products indicated deletion of the *pst* sequences between the two gRNA target sites. If deletion fragments were detected among the tested progeny of a given founder male, additional male progeny flies derived from this founder male were crossed individually to virgins *w*\*; *Sb*/TM3, *Ser*. Some progeny from these crosses were again analyzed with the PCR assays described above for the identification of flies with intragenic *pst* deletions. Two independent lines were established: *w*\*; *pst*^cc1^/TM3, *Ser* and *w*\*; *pst*^cc2^/TM3, *Ser*. DNA sequencing of the PCR fragment spanning the intragenic deletions revealed the presence of the following mutations. The *pst*^cc1^ allele is caused by an intragenic out-of-frame deletion. It is predicted to express a protein that includes only the first 94 amino acids of Pst. The *pst*^cc2^ allele is caused by an intragenic in-frame deletion. The Pst-PE variant protein expressed from this allele is predicted to lack a central region of 309 amino acids (…VVGSSGTSSVSSNCCN-deletion-GKGFLLNGDV…).

## Supporting information

All Figures and Supplementary Figures

---

**S1 Fig.**
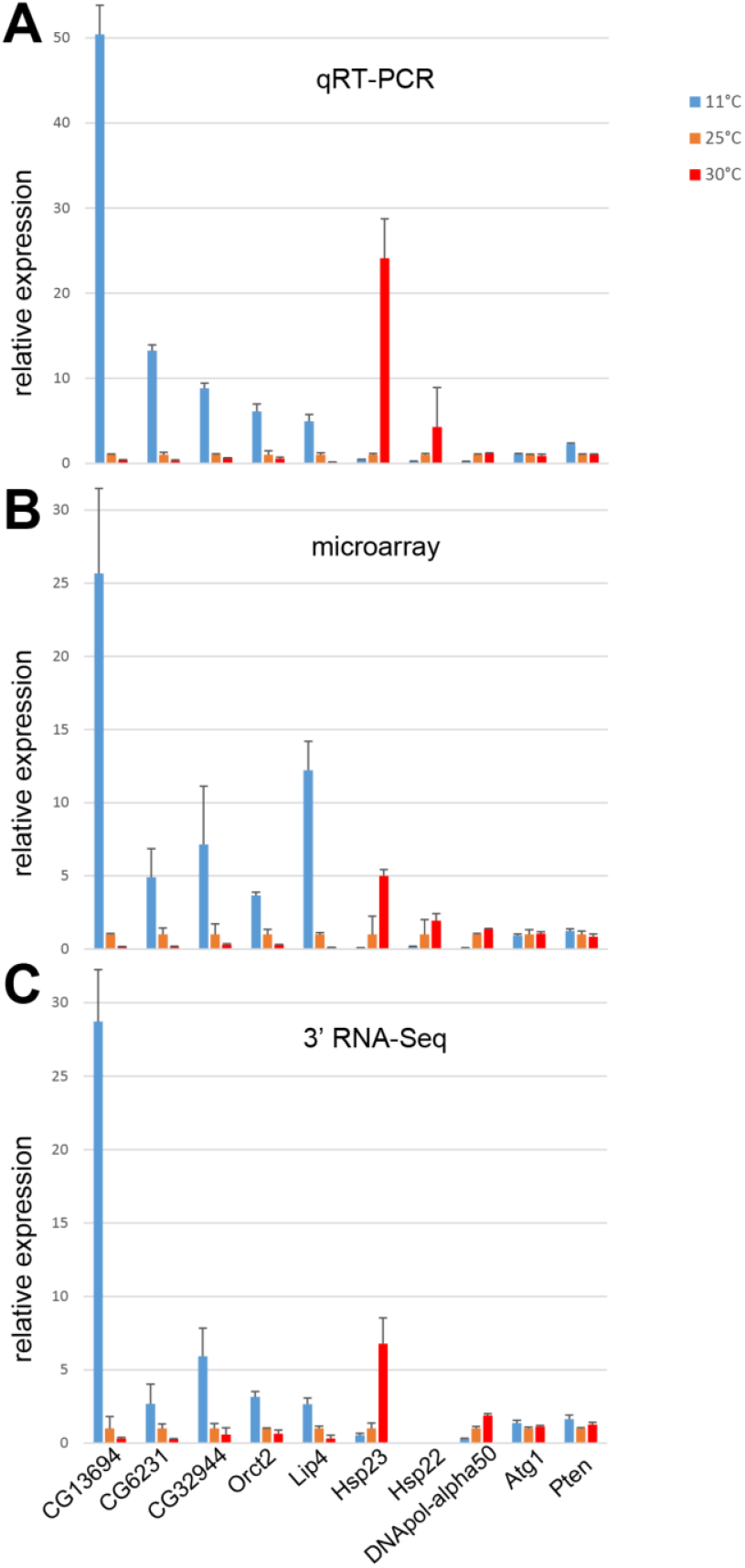
Concordance of transcript levels after quantification with different methods. (**A-C**) Temperature effects on transcript levels in S2R+ cells were analyzed using either qRT-PCR (**A**), microarrays (**B**) or 3’ RNA-Seq (**C**). The bar diagram displays relative expression levels at the different temperatures, as detected by the different methods. Expression at 25°C was set to 1. For each analysis, culture aliquots were shifted for 24 hours to the indicated temperatures (11, 25 and 30°C) before RNA isolation. Based on the results of the microarray data, representative CoolUp genes (*CG13694*, *CG6321*, *CG32944, Orct2*, *Lip4*) and CoolDown genes (*Hsp23*, *Hsp22*, *DNApol-α50*), as well as genes (*Atg1*, *Pten*), which were barely temperature regulated, were selected for validation by qRT-PCR. In case of qRT-PCR, mean and standard deviation of three technical replicates are displayed. In case of the microarray data, mean and standard deviation of three biological replicates and multiple probes, if present, are shown. In case of the 3’ RNA-Seq data, mean and standard deviation of three biological replicates are presented. *Hsp22* transcripts were not detected by 3’ RNA-Seq.

**S2 Fig.**
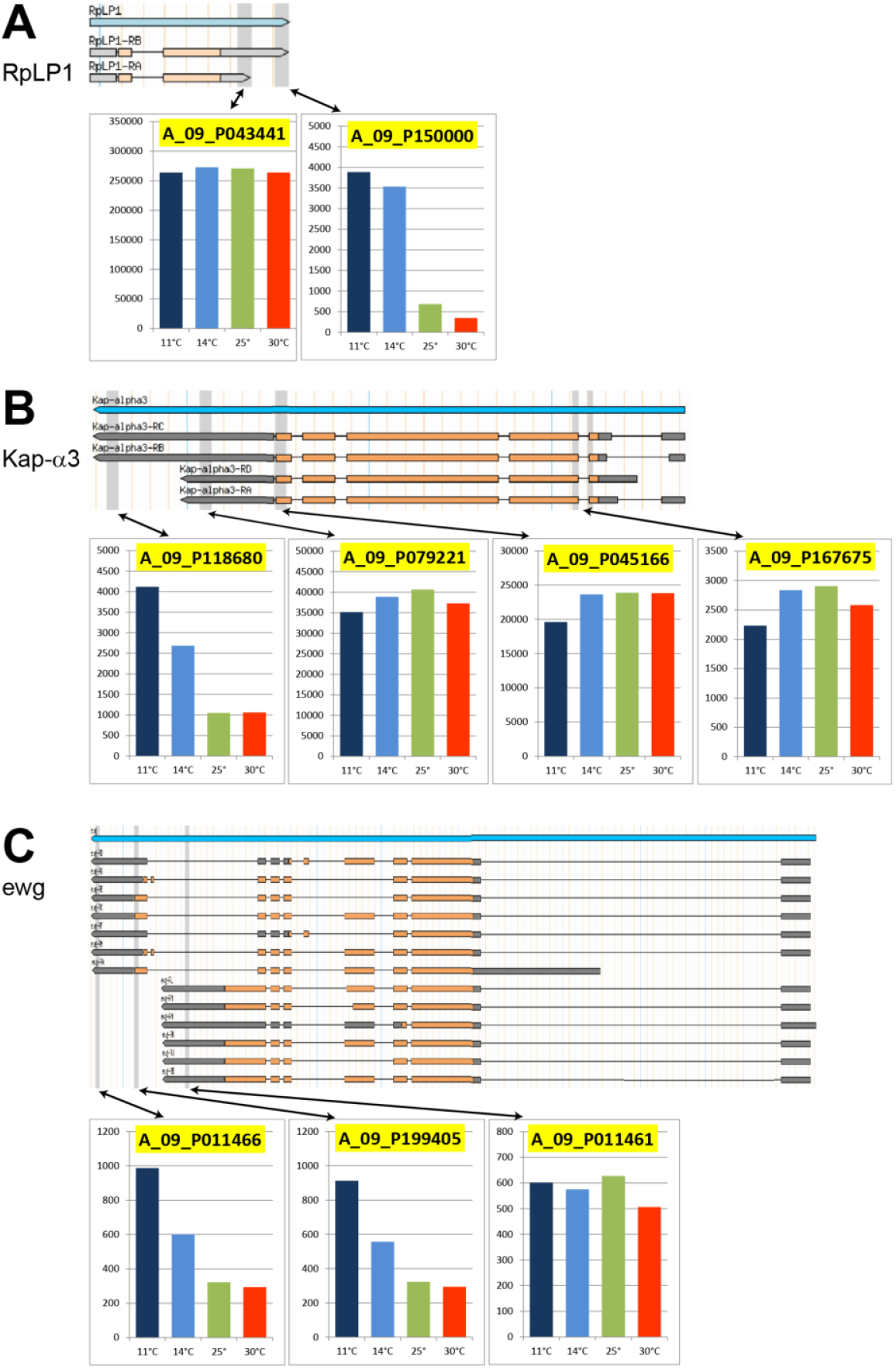
Increased use of distal polyadenylation sites at low temperature. (**A-C**) Microarray probes with pronounced CoolUp signals were observed to detect specifically the annotated transcripts with the longest 3’ untranslated region, as illustrated in case of the gene (**A**) *RpLP1*, (**B**) *Kap-α3* and (**C**) *ewg*. The positions recognized by different probes are indicated (vertical bars of light grey shading) in a scheme with the transcribed region and the annotated transcripts. The bar diagrams below display signal intensities (mean of three biological replicates) observed with these probes after incubation of S2R+ cells at different temperatures (11, 14, 25 and 30°C).

**S3 Fig.**
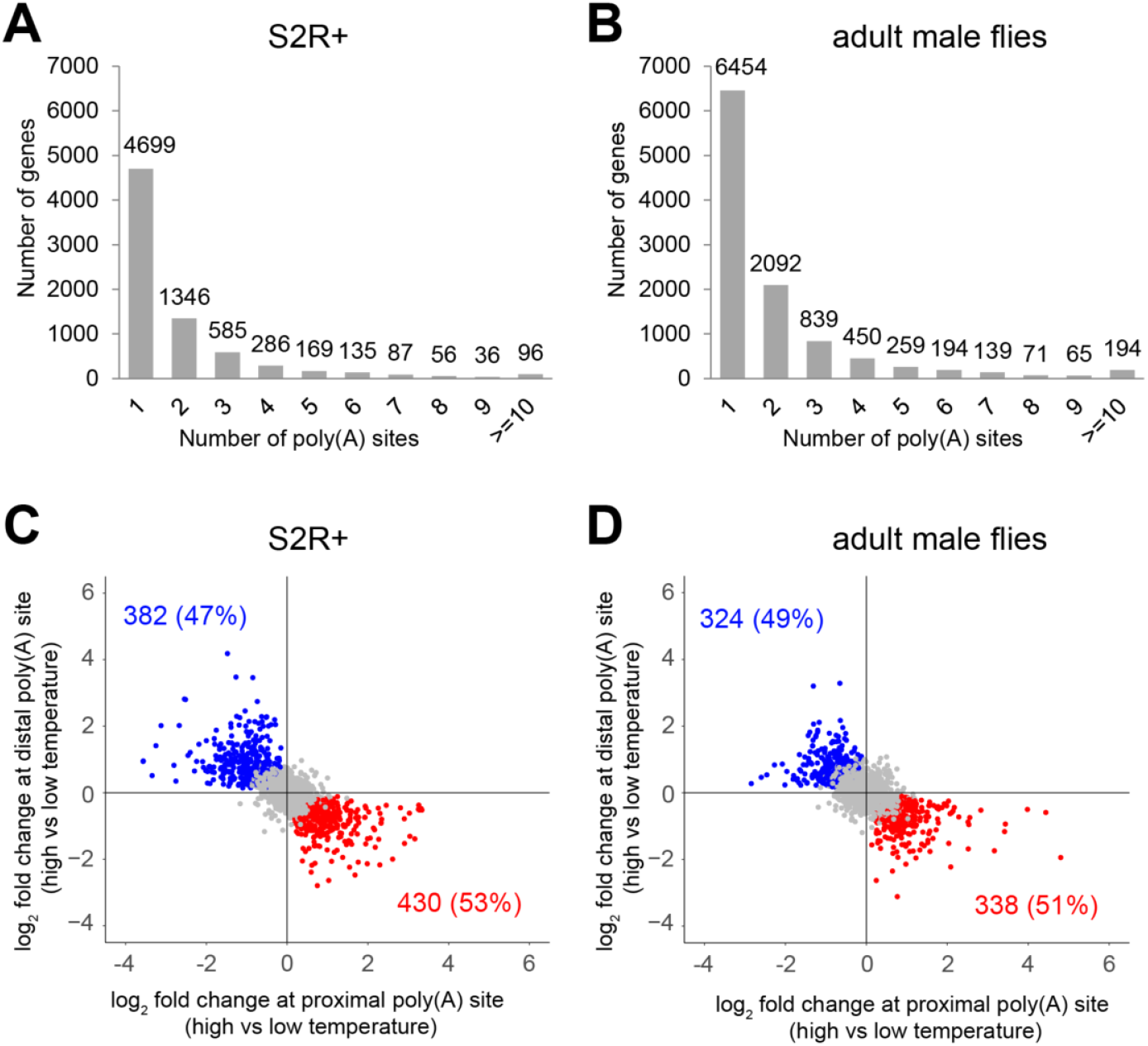
Temperature-regulated alternative polyadenylation. (**A-D**) 3’ RNA-Seq data obtained from S2R+ cells (**A, C**) and adult male flies (**B, D**) after 24 hours of incubation at different temperatures (11, 14, 25 and 30°C) was used for an analysis of temperature effects on the choice of polyadenylation sites (PASs). (**A,B**) Histograms illustrate the frequency of alternative polyadenylation (APA) in S2R+ cells (**A**) and adult male flies (**B**), as detected after pooling all data obtained at the different temperatures from the three biological replicates. In total, 14669 PASs were detected in S2R+ cells, and 22378 in adult male flies. These were assigned (see Materials and Methods) to a total of 7495 and 10757 expressed genes in S2R+ cells and adult male flies, respectively. (**C,D**) Temperature effects on APA. For genes with multiple PASs, we compared the two PASs with highest presence (read counts) across all conditions and required the log_2_ fold change to be in opposite directions when comparing the read counts at the high (25 and 30°C) with those at the low (11 and 14°C) temperatures. The scatter plots display the fold changes at the proximal and distal of these two most strongly affected PASs in S2R+ cells (**C**) and adult flies (**D**). Blue dots represent genes (class I), where preference of the distal over the proximal PAS at low temperature is significant in comparison to the high temperature. Red dots represent genes (class II), where temperature change has a significant effect in opposite direction.

**S4 Fig.**
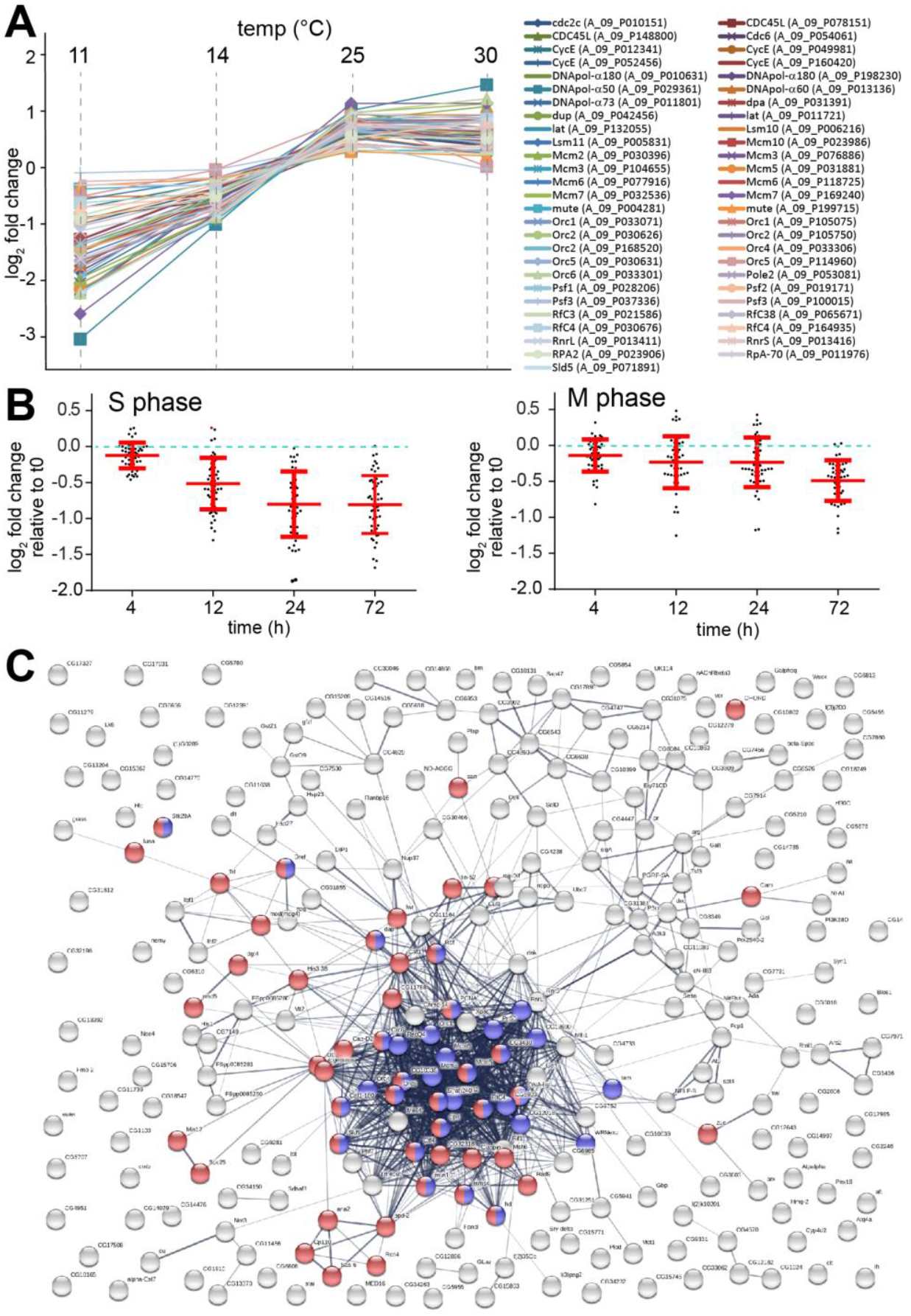
Downregulation of cell cycle genes in S2R+ cells at low temperature. (**A**) Temperature dependence of transcript levels of S phase genes. The S2R+ cell transcriptome was analyzed after 24 hours of incubation at different temperatures (11, 14, 25 and 30°C) using microarrays. Temperature dependence of signal intensities (mean of three biological replicates) observed with microarray probes for transcripts derived from a curated set of genes crucial for progression through S phase is displayed after baseline transformation. (**B**) Temporal dynamics S- and M phase gene downregulation after a shift from 25 to 14°C. Transcriptome changes were analyzed with microarrays at different times (0, 4, 12, 24 and 72 hours) after the temperature downshift. Fold change relative to expression at t = 0 (dashed green line) of signal intensities (mean of three biological replicates) observed with microarray probes for transcripts derived from a curated set of genes crucial for progression through S- and M phase was calculated. Mean and standard deviation are displayed in red. (**C**) Clustering of temperature-regulated genes according to their temporal expression dynamics after a 25->14°C shift using *k*-means identified a prominent cluster of genes with persistent downregulation (Fig. 3E, cluster 2). Functional interactions among the genes in this cluster, as revealed by analysis with the STRING database, are displayed. Genes associated with the GO term “DNA replication” (GO:0006260), the term most strongly enriched (FDR = 4.15×10^−26^) by the genes in this cluster, are marked in blue. Genes associated with “cell cycle” (GO:0007049, also strongly enriched, FDR = 3.72×10^−15^) are marked in red.

**S5 Fig.**
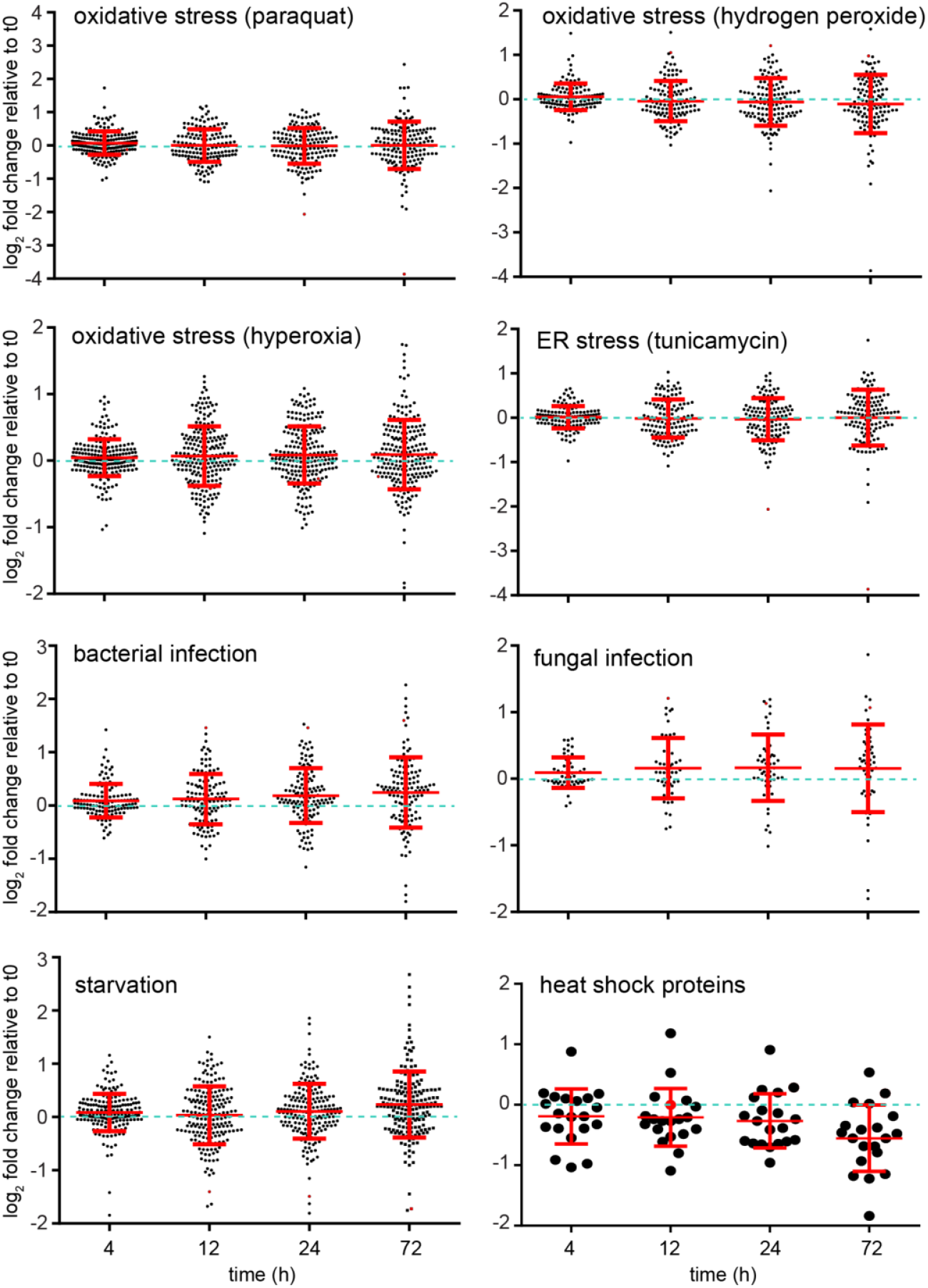
Absence of stress response gene induction after temperature downshift to 14°C. Microarray data from the time course analysis of S2R+ cell transcriptomes after a 25->14°C temperature shift were used for an analysis of the response of known stress response genes. Probes detecting transcripts of genes previously reported to be induced by the indicated stressors were identified. Signals obtained at t0 were set to 1 and fold change at different times after temperature downshift was calculated. Swarm plots display log_2_ values of the fold changes, as well as means and standard deviation in red.

**S6 Fig.**
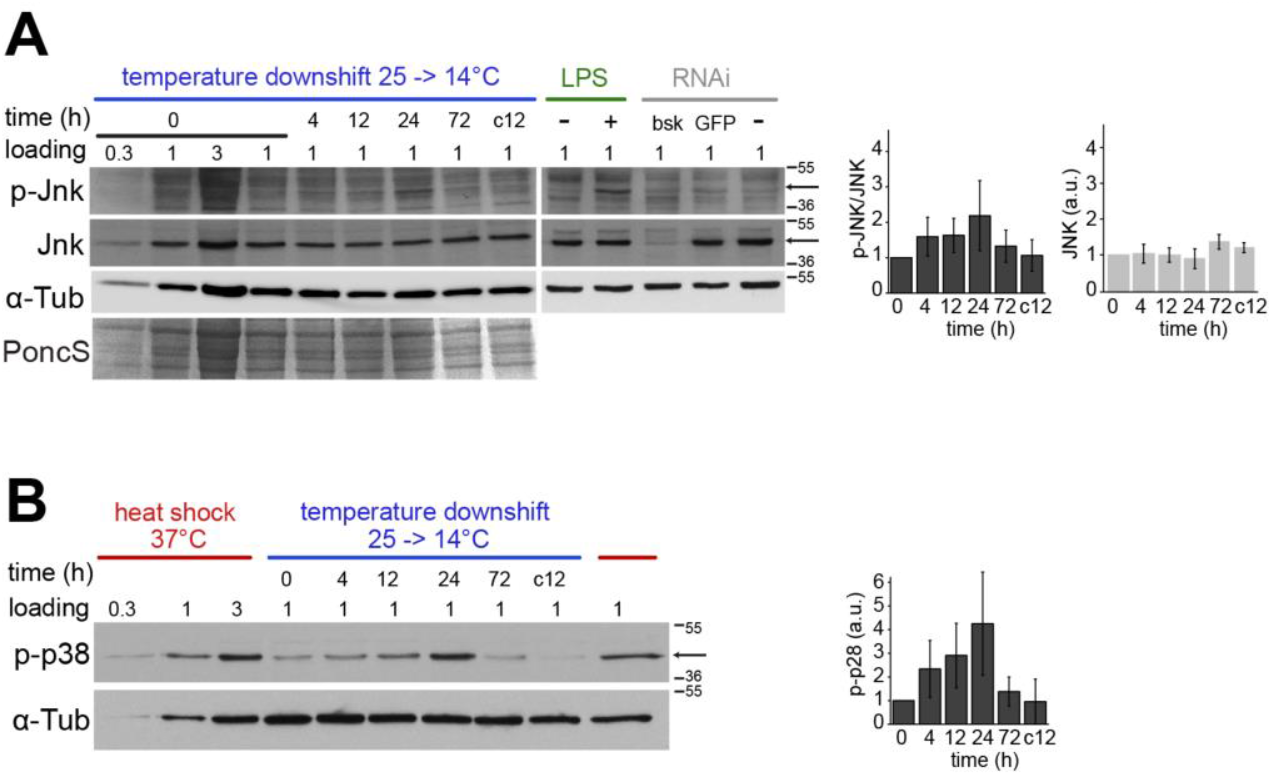
Transient activation of JNK and p38 protein kinases after temperature downshift to 14°C. (**A,B)**Immunoblotting with antibodies specific for the activated forms of JNK (phospho-JNK) and p38 (phospho-p38) were used for analysis. S2R+ cells were plated and grown at 25°C before a shift to 14°C at t = 0. Extracts were prepared at t = 0, 4, 12, 24, and 72 hours after downshift, and also from an aliquot maintained for 12 hours at 25°C after t = 0. Relative amounts of extracts analyzed by immunoblotting are indicated. Arrows mark the bands corresponding to phospho-JNK, JNK and phospho-p38, and dashes the position of molecular weight markers. (**A**) For control of antibody specificities, we also analyzed extracts from S2R+ cells treated with lipopolysaccharide (LPS) or depleted of *bsk* transcripts (coding for JNK) or GFP transcripts (for control) by RNAi. Immunoblots were stained with Ponceau S (PoncS) and probed with anti-phospho-JNK, anti-JNK and anti-α-tubulin. Signal intensities in the bands representing JNK and phospho-JNK were quantified in four replicates. A bar diagram (+/− s.d.) displays average intensities normalized to those at t = 0. (**B**) For comparison, we also analyzed extracts from S2R+ cells after exposure (45 min) to a heat shock at 37°C. Relative amounts of extracts analyzed by immunoblotting are indicated. Immunoblots were probed with anti-phospho-p38 and anti-α-tubulin. The arrow indicates the band corresponding to phospho-p38. Position of molecular weight markers are indicated. Signal intensities in the bands representing phospho-p38 were quantified in three replicates. A bar diagram (+/− s.d.) displays average intensities normalized to those at t = 0.

**S7 Fig.**
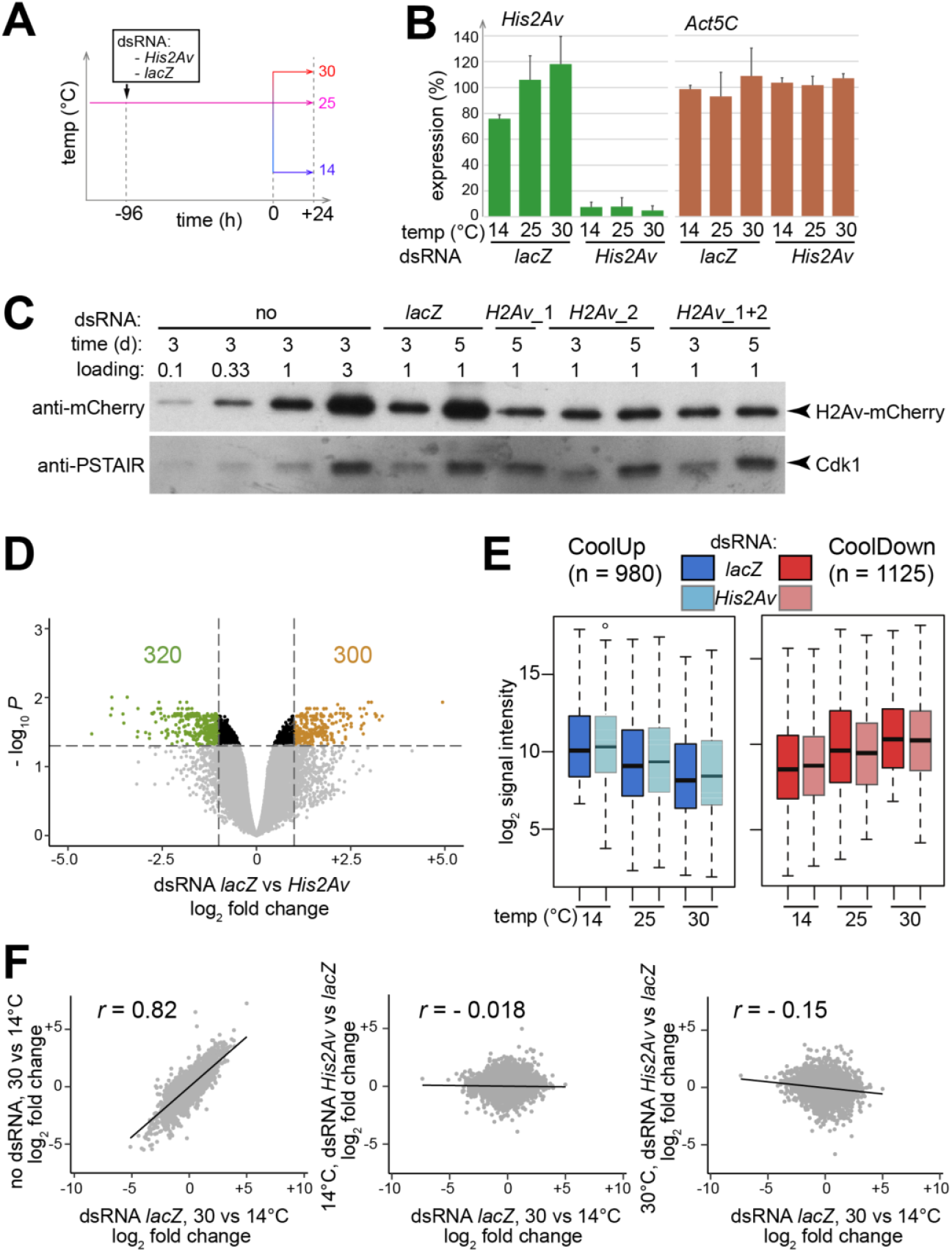
Evaluation of the role of histone H2Av (His2Av) in transcriptional control of temperature-regulated genes in S2R+ cells. (**A**) To address the role of His2Av in temperature-dependent regulation of gene expression in S2R+ cells, we added *His2Av* dsRNA or *lacZ* dsRNA for control to culture aliquots followed by a shift to the indicated temperatures after four days. Twenty-four hours after the shift, we isolated total RNA for analysis with microarrays. Three replicate experiments were performed. (**B**) Analysis of transcript levels of *His2Av* and *Act5C* (for control) confirmed successful depletion of *His2Av* transcripts. Bar diagrams display transcript levels (average of three biological replicates and standard deviation) at the analyzed temperatures. Average expression at the three temperatures observed after treatment with *lacZ* dsRNA was set to 100%. (**C**) Analysis of His2Av depletion by immunoblotting. S2R+_*MtnA*p-*His2Av*-mRFP cells (Lidsky et al. 2013) were treated with dsRNA derived from *lacZ* (for control) and two distinct *His2Av* amplicons (*H2Av*_1 and *H2Av*_2) for three and five days as indicated. Cell extracts were analyzed by immunoblotting with anti-mCherry (detecting His2Av-mRFP) and anti-PSTAIR (loading control). The combined treatment with both *His2Av* dsRNA preparations (*H2Av*_1+2) for five days was found to reduce His2Av-mRFP levels to 40% of controls according to quantification of signal intensities and normalization based on anti-PSTAIR signals. (**D**) Effects of *His2Av* depletion on the S2R+ cell transcriptome. A volcano plot is displayed with probes associated with signal intensities that were significantly (FDR < 0.05; fold change ≥ 2) down-(green dots, 320 probes) or upregulated (brown dots, 300 probes) by *His2Av* depletion in comparison to control (*lacZ* dsRNA treatment). The comparison based on signals detected at 14°C is shown. Analogous comparisons at 25 or 30°C resulted in comparable observations. (**E**) Effects of *His2Av* depletion on transcript levels of temperature-regulated genes. Probes associated with a fold change in signal intensity ≥ 2 in the comparison of 14 and 30°C in both control experiments (with *lacZ* dsRNA treatment and without any dsRNA treatment) were selected for further analysis. Plots summarize the signal intensities obtained with these probes, separated into CoolUp and CoolDown probes, after treatment with either *lacZ* or *His2Av* dsRNA at the three different temperatures. The comparison revealed at most subtle effects of *His2Av* depletion on transcript levels of temperature-regulated genes in S2R+ cells, in contrast to results from *Arabidopsis thaliana* where a reduction in the homolog His2A.Z has been found to result in a dramatic conversion of the expression of genes that are normally temperature-regulated into a constitutive warm-like expression (Kumar and Wigge 2010). (**F**) Fold change correlation analyses for further analysis of the effect of *His2Av* depletion on temperature-dependent gene regulation. Scatter plots of log2 fold changes for all probes with significant signals are displayed. *r* = Pearson’s correlation coefficient. As revealed by the left panel, the fold changes observed when comparing expression levels at 14 and 30°C were very strongly correlated in the two control experiments (with *lacZ* dsRNA treatment and without any dsRNA treatment). However, there was essentially no correlation between the fold changes resulting from temperature change and those resulting from His2Av depletion at 14°C (middle panel) or 30°C (right panel), in contrast to results from similar analyses in *Arabidopsis thaliana* (Kumar and Wigge 2010), where reduced loading of the homolog H2A.Z into chromatin (in *arp6-10* mutants) results in *r* = 0.68 for the correlation between transcriptome regulation by temperature and misregulation by reduced H2A.Z.

**S8 Fig.**
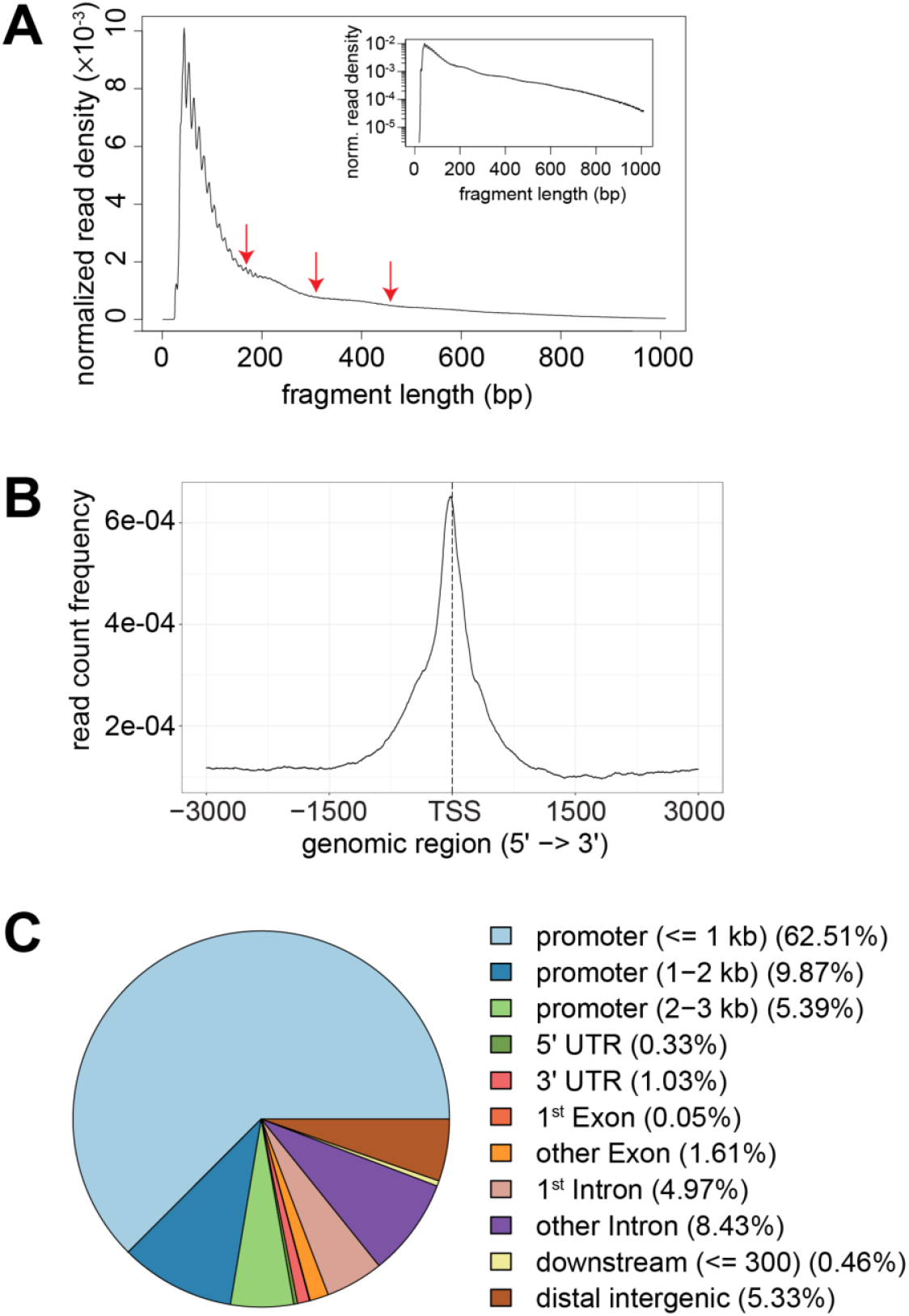
DNA accessibility in nuclear chromatin of S2R+ cells. For illustration of ATAC-Seq data quality, various analyses based on one of the samples (replicate of cells tagmented after growth at 25°C) are displayed. (**A**) Distribution of ATAC-Seq fragment lengths. Arrows indicate periodicity reflecting nucleosomal organization. (**B**) Enrichment of ATAC-Seq reads around annotated transcription start sites (TSS). Distance (bp) from the transcriptional start site (TSS) is indicated on the X axis. (**C**) Distribution of ATAC-Seq reads across annotated genome features.

S9 Fig. Assay for temperature-dependent enhancer activity with candidate CREs.

(**A**) Table summarizing location and activity of candidate CREs analyzed after RMCE with SR9rg cells.

(**B**) Data for the analyzed candidate CREs. Candidate CREs (designation in bold on the left side) were selected based on data obtained by ATAC-Seq (top three browser tracks) and 3’ RNA-Seq (bottom three browser tracks). Grey shading represents the region of the analyzed DNA fragment. Green arrowheads indicate the direction of transcription of the proximal gene. Scatter plots on the right side present green (x axis) and red (y axis) fluorescence intensities as determined by flow cytometry with SR9rg cells after RMCE and incubation at the indicated temperatures.

S10 Fig. Dissection of *pst_E1* region

(**A**) Schematic illustration of analyzed truncation series, with transcription start sites (kinked arrows), untranslated regions (grey boxes), coding region (yellow boxes) and introns (white boxes) indicated.

(**B**) Scatter plots of the results obtained by flow cytometry after RMCE with SR9rg cells and incubation at the indicated temperatures.

**S11 Fig.**
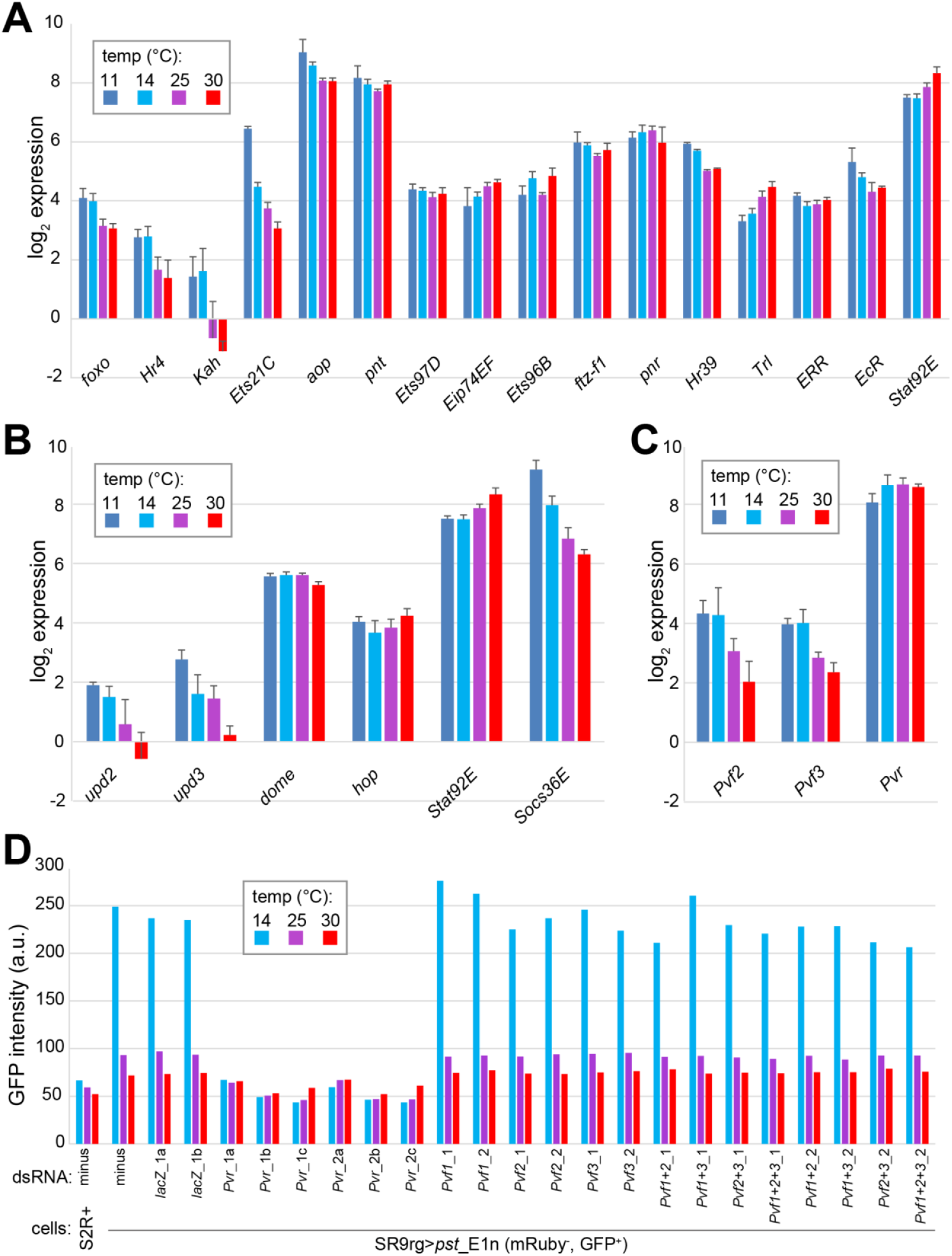
Temperature dependence of transcript levels of candidate regulators of *pst_E1n* enhancer activity and dependence on Pvf/Pvr. (**A-C**) Transcript levels at the indicated temperatures in S2R+ cells as detected by 3’ RNA-Seq. Bar diagrams display log2 of the average of the read counts (after normalization) and s.d. (n = 3 replicate experiments). (**A**) Transcript levels of TFs with predicted binding sites within *pst*_E1n and detectable expression in S2R+ cells. (**B**) Transcript levels of JAK/STAT signal transduction proteins. Expression of *upd1* is marginal at most in S2R+ cells and hence not included. (**C**) Transcript levels of the genes encoding the receptor tyrosine kinase Pvr and its known ligands Pvf2 and Pvf3. Expression of *Pvf1* is marginal at most in S2R+ cells and hence not included. (**D**) Dependence of *pst*_E1n enhancer activity on Pvfs and Pvr. The indicated dsRNAs were used for depletion in SR9rg>*pst*_E1n (mRuby^−^, GFP^+^) cells, which were then shifted in aliquots to the indicated temperatures and analyzed eventually by flow cytometry. Bars display median GFP signal intensity. For depletion of Pvr and Pvfs, two dsRNA preparations (1 or 2) generated from distinct amplicons, were used in multiple experiments (indicated by a, b or c) in some cases.

**S12 Fig.**
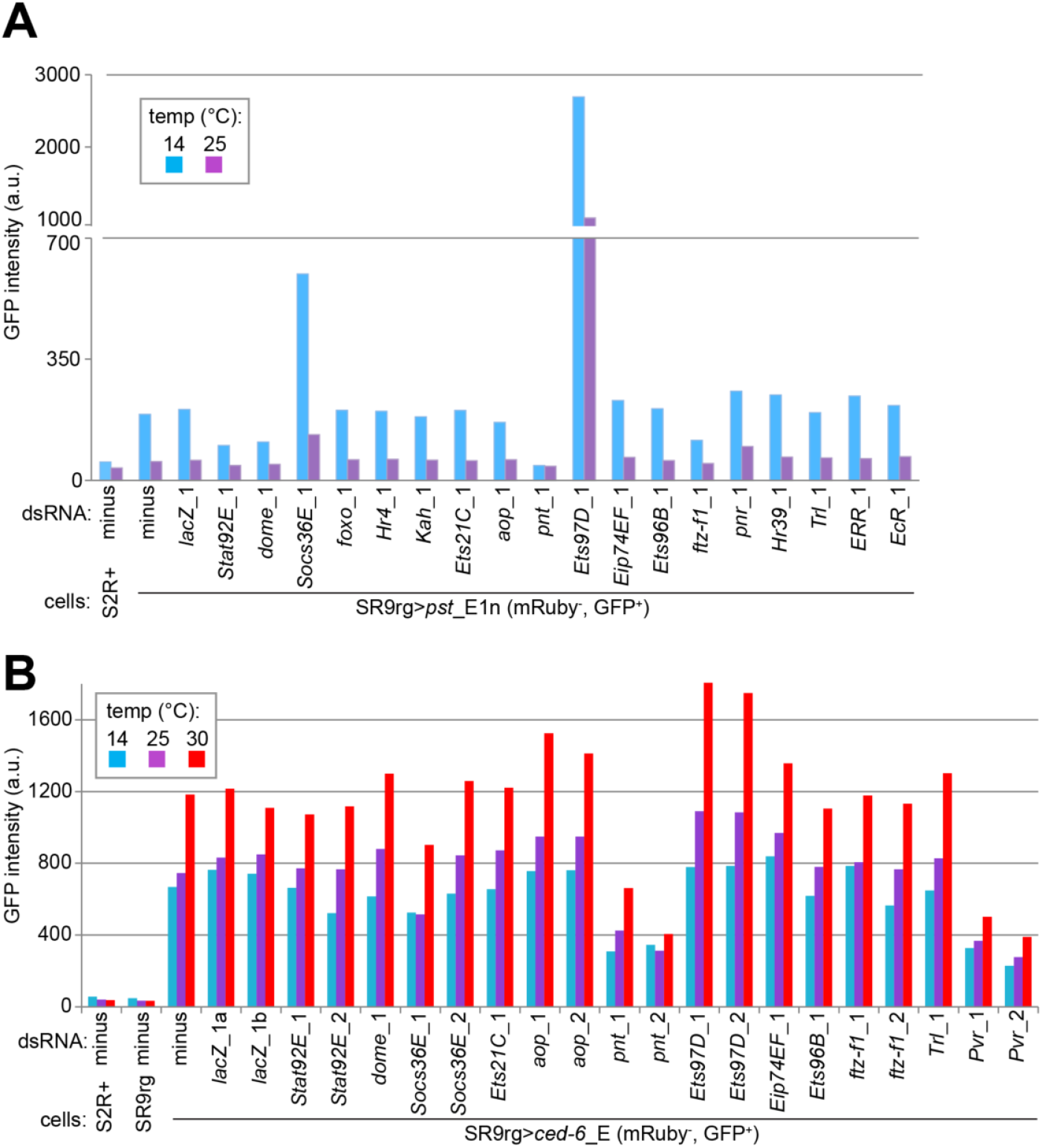
Enhancer activity after depletion of TFs, JAK/STAT and Pvr. (**A,B**) The indicated dsRNAs were used for depletion of TFs with predicted binding sites in *pst*_E1n. In addition, JAK/STAT signaling pathway proteins (Dome and Socs36E) and the receptor tyrosine kinase Pvr were depleted. In some cases, two dsRNA preparations (1 or 2) generated from distinct amplicons were used. *lacZ* ds RNA was used in two replicates (a and b). Depletions were performed in SR9rg>*pst*_E1n (mRuby^−^, GFP^+^) cells (**A**) and in SR9rg>*ced-6*_E (mRuby^−^, GFP^+^) cells (**B**). Depleted cells and control cell lines (S2R+ and SR9rg) were then shifted in aliquots to the indicated temperatures and analyzed eventually by flow cytometry. Bars display median GFP signal intensity.

**S13 Fig.**
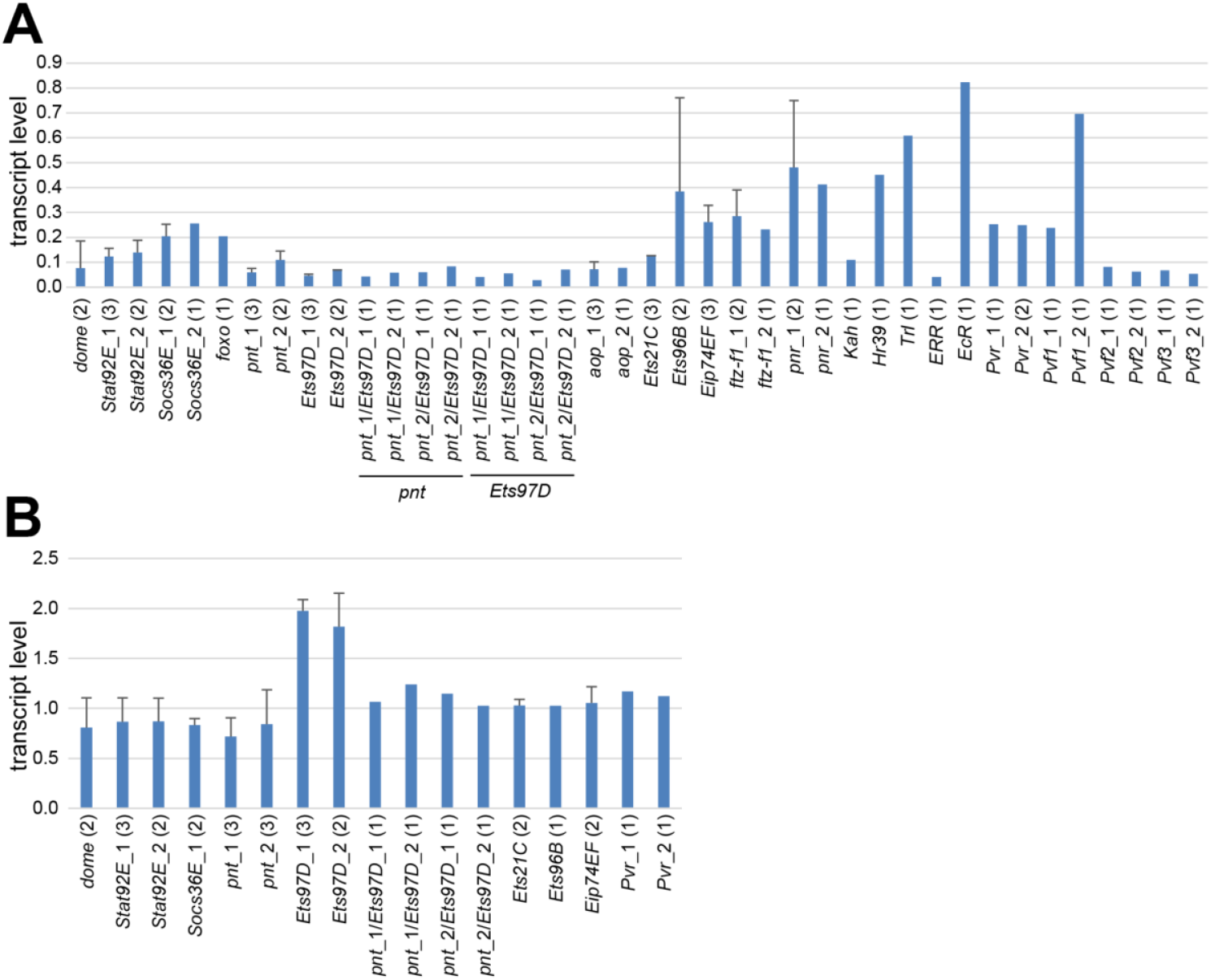
Depletion efficiency and effect on endogenous *pst* transcript levels. (**A,B**) The indicated dsRNAs were used for RNAi with SR9rg>*pst*_E1n (mRuby^−^, GFP^+^) cells. Two dsRNA preparations (1 or 2) generated from distinct amplicons were used for some targets. The number of independent experiments is indicated in brackets. In addition, *lacZ* dsRNA was used in parallel for control. Bars represent average transcript levels as determined by qRT-PCR relative to those detected after *lacZ* depletion, which were set to 1. Whiskers indicate s.d. in case of multiple independent experiments. (**A**) To assess RNAi efficiency, total RNA was isolated after four days of depletion at 25°C followed by analysis of the target transcript levels. The target analyzed in case of double depletion is indicated at the bottom. (**B**) The levels of transcripts derived from the endogenous *pst* gene were analyzed.

**S14 Fig.**
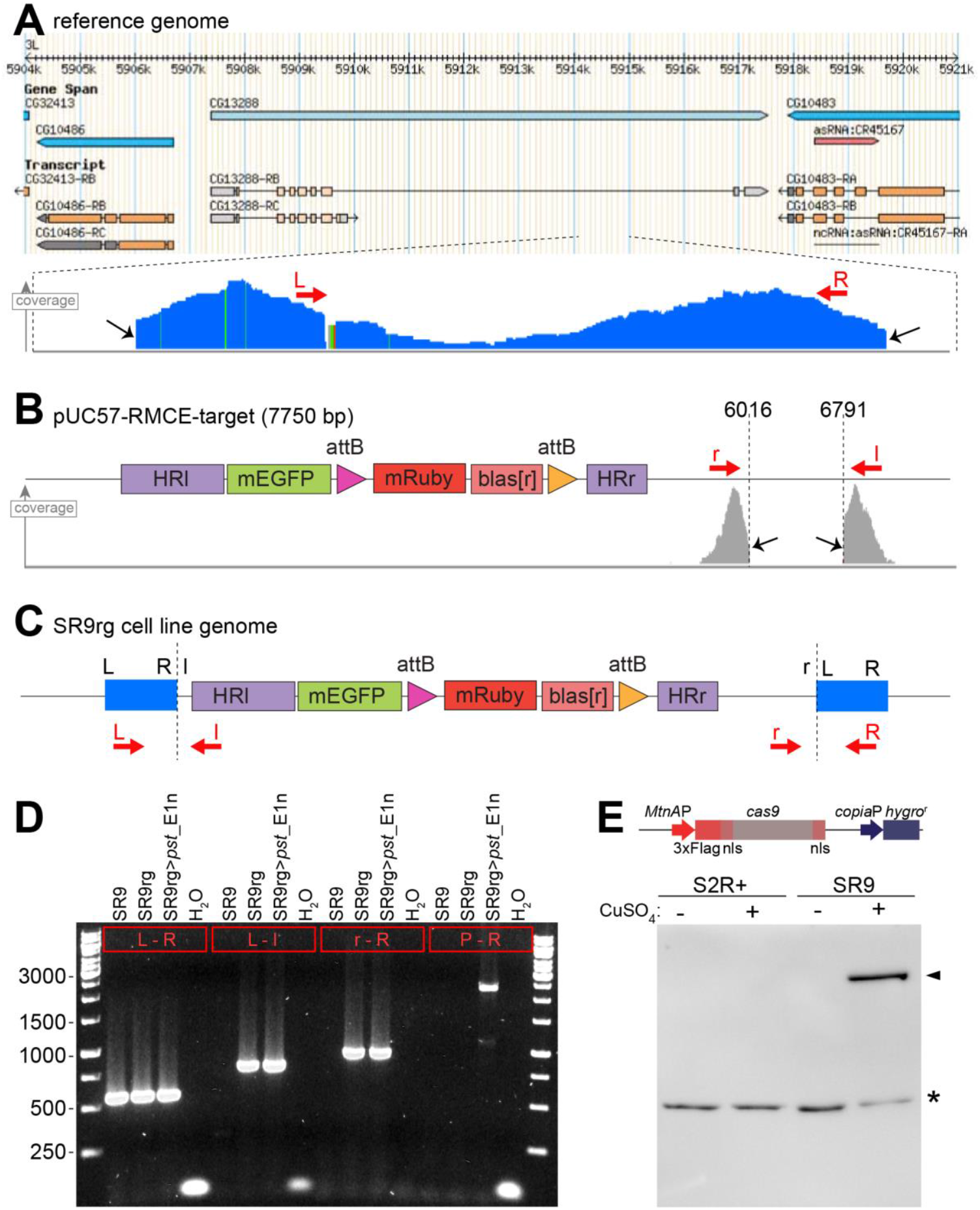
Characterization of SR9 and SR9rg cells. (**A-D**) Identification of a functional target locus for RMCE in SR9rg cells. Whole genome sequencing was used for the characterization of the clonal SR9rg cell line. Hybrid paired reads, where one read aligns to genomic sequences and the mate read to pUC57-RMCE-target plasmid sequences were identified and used for mapping chromosomal insertions of pUC57-RMCE-target. An insertion that is competent for RMCE was identified on chromosome 3L in an intron of the *CG13288* gene. (**A**) The genomic region (dmel_r6.19, chr3L:5904000 to chr3L:5921000) with *CG13288* as visualized by GBrowse 2.0 is displayed on top. The cumulative base coverage by hybrid paired reads mapping to this region is indicated by the blue bars below (from 0 to 55 reads/bp). Black arrows emphasize the distal clustering of the abrupt end of the alignments on either side, corresponding to the genome-plasmid junctions. Primers “L” and “R” (red arrows) with sequences close to the left and right ends of this genomic region were used for confirmation of the insertion by PCR (see panel D). (**B**) A linearized representation of pUC57-RMCE-target is shown on top. The grey bars below display the cumulative base coverage (from 0 to 51 reads/bp) by the hybrid paired reads mapping to the *CG132888* region. Black arrows and dashed lines indicate breakpoints (abrupt clustered ends of read alignments). Primers “r” and “l” (red arrows) with sequences close to the right and left ends of the plasmid region were used for confirmation of the insertion by PCR (see panel D). (**C**) Proposed organization of pUC57-RMCE-target insertion in the *CG13288* region. The plasmid pUC57-RMCE-target is inserted almost completely, starting from breakpoint “l” and extending to the breakpoint “r” (see panel B). The inserted plasmid region is flanked by a tandem duplication of the genome region between “L” and “R” (see panel A). Primers used for confirmation of the proposed organization are indicated (red arrows). (**D**) PCR assay for confirmation of the proposed organization of the pUC57-RMCE-target insertion in the *CG13288* region and of its competence for RMCE. Genomic DNA isolated from the indicated cells was analyzed by PCR with the primer pairs indicated in the red boxes. Primer P in the last pair anneals to the *pst*_E1n region, which is only present after RMCE, i.e., only in the SR9rg>*pst*_E1n cells. L = oEC264, R = oEC267, l = oEC263, r = oEC265, P = YB654. (**E**) Characterization of SR9 cells. These cells were obtained after transfection of S2R+ cells with the plasmid pMT-cas9-hygro^r^ schematically shown on top. For confirmation of CuSO_4_-inducible expression of *cas9* in the SR9 cells, they were cultured in parallel with the parental S2R+ cells for 24 hours without or with 500 μM CuSO_4_ as indicated. Total extracts were probed by immunoblotting with an anti-FLAG. Ponceau S staining (not shown) indicated comparable sample loading, as also indicated by a non-specific anti-FLAG band (*). The specific signal representing 3xFlag-nls-Cas9-nls (predicted molecular weight 164.67 kDa) is indicated as well (arrowhead).

## Acknowledgements

We would like to thank Andre Koch for initial contributions, Björn Handke for the generation of HB10 cells, Jessica Bader for technical help in particular during the generation of SR9rg cells, Helen Lindsay and Michael Frochaux for support during initial ATAC-Seq data analysis and prediction of TF binding sites, and Sina Moser for technical support all along. We are also very grateful to the Functional Genomics Center Zurich and its staff for their support of the microarray, RNA-Seq and ATAC-Seq analyses.

## Supplementary Information

### Supplementary tables

- S1 Table: temperature dependence of the S2R+ cell transcriptome as detected by DNA microarray analysis

- S2 Table: log2 values of the fold changes in transcript levels observed when comparing low and high temperatures by DNA microarray analysis with S2R+ cells (worksheet 1), 3’ RNA-Seq with S2R+ cells (worksheet 2), 3’ RNA-Seq with adult male flies (worksheet 3) and DNA microarray analysis with HB10 cells (worksheet 4). Expression at low temperature represents the mean observed at 11 and 14°C, while expression at high temperature the mean at 25 and 30°C, except for the analyses with HB10 cells where expression at 14 and 24°C is compared.

- S3 Table: temperature dependence of the S2R+ cell transcriptome as detected by 3’ RNA-Seq

- S4 Table: temperature dependence of the transcriptome of adult flies as detected by 3’ RNA-Seq

- S5 Table: temperature dependence of the HB10 cell transcriptome as detected by DNA microarray analysis

- S6 Table: temperature effects on the use of alternative polyadenylation sites for genes in S2R+ cells (worksheet 1) and adult male flies (worksheet 2)

- S7 Table: temporal dynamics of transcriptome changes in S2R+ cells after a shift to 14°C as detected by DNA microarray analysis

- S8 Table: Effects of RNAi with *His2Av* dsRNA or *lacZ* dsRNA (for control) on temperature dependence of the S2R+ cell transcriptome as detected by DNA microarray analysis.

- S9 Table: temperature dependence of DNA accessibility in S2R+ cells as detected by ATAC-Seq

- S10 Table: synthetic DNA (oligos and gene blocks)

- S11 Table: source data

