## Supplementary material for "A *cis*-regulatory element promoting increased transcription at low temperature in cultured ectothermic *Drosophila* cells": All Figures and Supplementary Figures

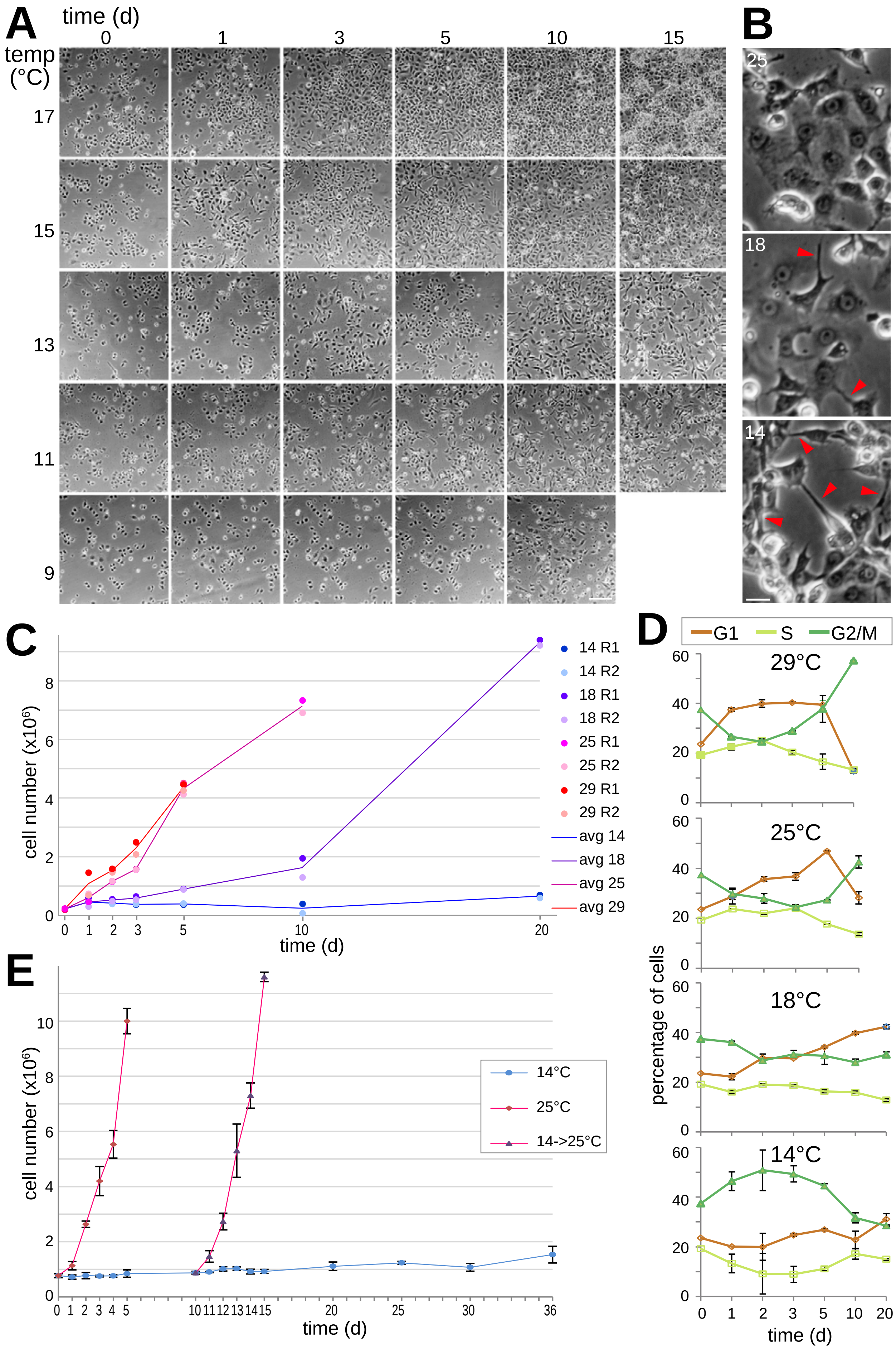
A
time (d)
0 1 3 5 10 15
temp (°C)
17
15
13
11
9
B
25
18
14
C
cell number (x10⁶)
time (d)
14 R1
14 R2
18 R1
18 R2
25 R1
25 R2
29 R1
29 R2
avg 14
avg 18
avg 25
avg 29
D
G1
S
G2/M
29°C
25°C
18°C
14°C
percentage of cells
time (d)
E
cell number (x10⁶)
14°C
25°C
14->25°C
time (d)

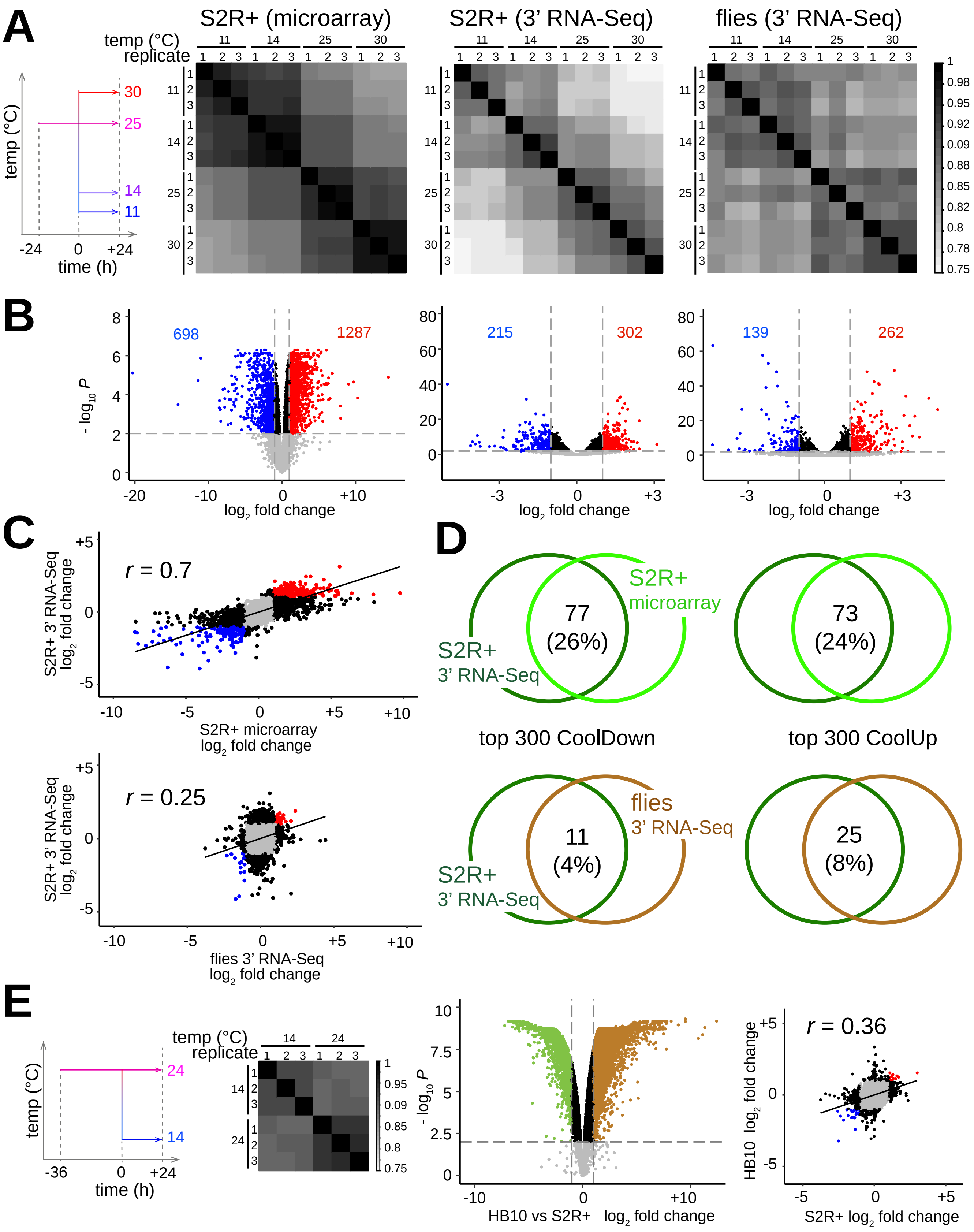
A
S2R+ (microarray)
S2R+ (3' RNA-Seq)
flies (3' RNA-Seq)
temp (°C)
replicate
time (h)
B
log2 fold change
- log10 P
698
1287
215
302
139
262
C
r = 0.7
S2R+ 3' RNA-Seq log2 fold change
S2R+ microarray log2 fold change
r = 0.25
flies 3' RNA-Seq log2 fold change
D
S2R+ microarray
S2R+ 3' RNA-Seq
77 (26%)
73 (24%)
top 300 CoolDown
top 300 CoolUp
flies 3' RNA-Seq
11 (4%)
25 (8%)
E
HB10 vs S2R+ log2 fold change
r = 0.36
HB10 log2 fold change
S2R+ log2 fold change

A

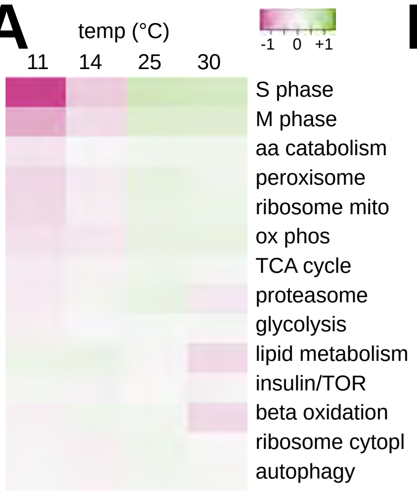


B

| term | | S2R+ | | HB10 | |
|---|---|---|---|---|---|
| | | enrichment (FDR) | | enrichment (FDR) | |
| MCM complex | GO:0042555 | 4.35 (3.93e-07) | D | ns | |
| ORC complex | GO:0005664 | 3.02 (1.87e-05) | D | ns | |
| Ndc80 complex | GO:0031262 | 2.12 (1.30e-04) | D | ns | |
| V-type ATPase | GO:0000221 | 1.89 (6.40e-04) | U | ns | |
| leucine degradation | GO:0006551 | 2.13 (2.90e-03) | D | 2.89 (1.40e-03) | D |
| branched aa degradation | GO:000981 | 2.24 (3.90-e04) | D | 2.86 (1.00e-04) | D |
| TCA cycle | GO:0006099 | ns | | 1.12 (5.00e-04) | D |
| mito electron transport | GO:0006119 | ns | | 1.10 (2.71e-05) | D |

ns = not significant D = CoolDown U = CoolUp

C

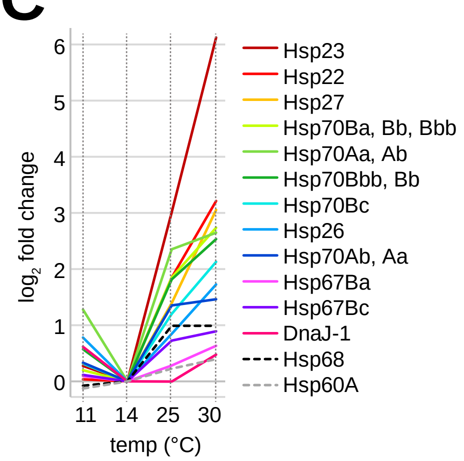


D

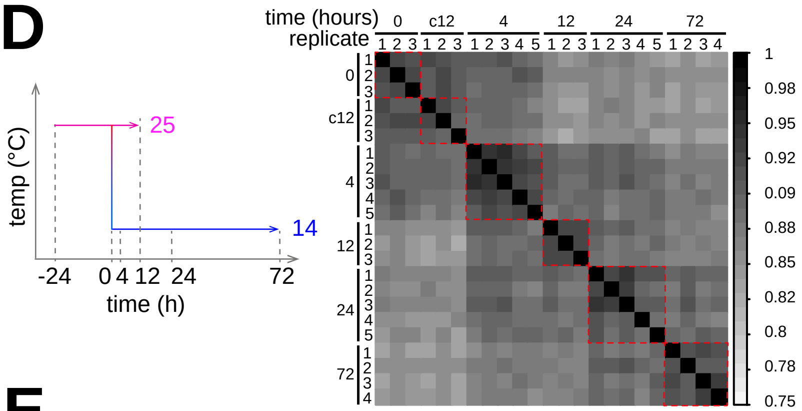


E

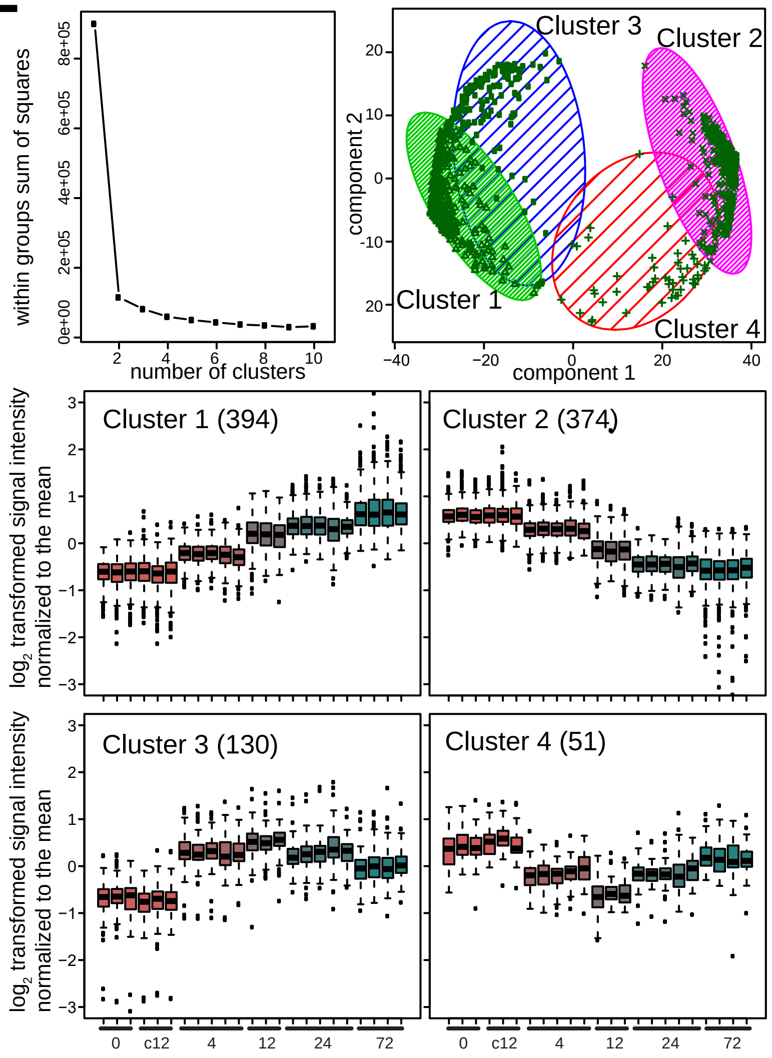


F

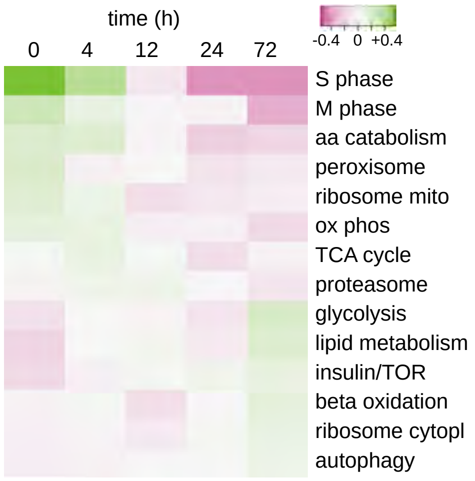

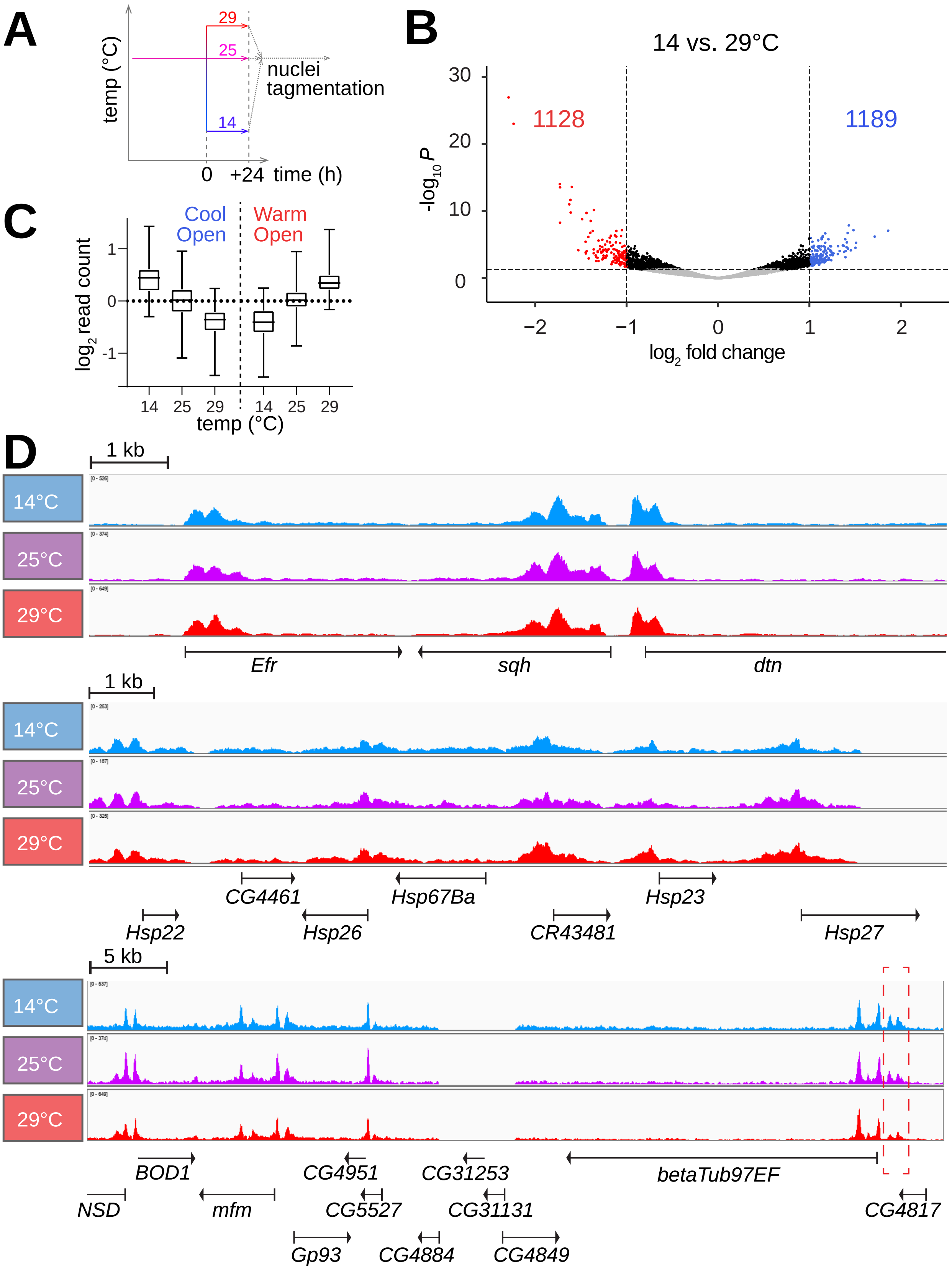
A
temp (°C)
29
25
14
nuclei tagmentation
0
+24
time (h)
B
14 vs. 29°C
$-log_{10}P$
1128
1189
$log_2$ fold change
C
Cool Open
Warm Open
$log_2$ read count
temp (°C)
D
1 kb
14°C
25°C
29°C
Efr
sqh
dtn
1 kb
CG4461
Hsp67Ba
Hsp23
Hsp22
Hsp26
CR43481
Hsp27
5 kb
BOD1
CG4951
CG31253
betaTub97EF
NSD
mfm
CG5527
CG31131
CG4817
Gp93
CG4884
CG4849

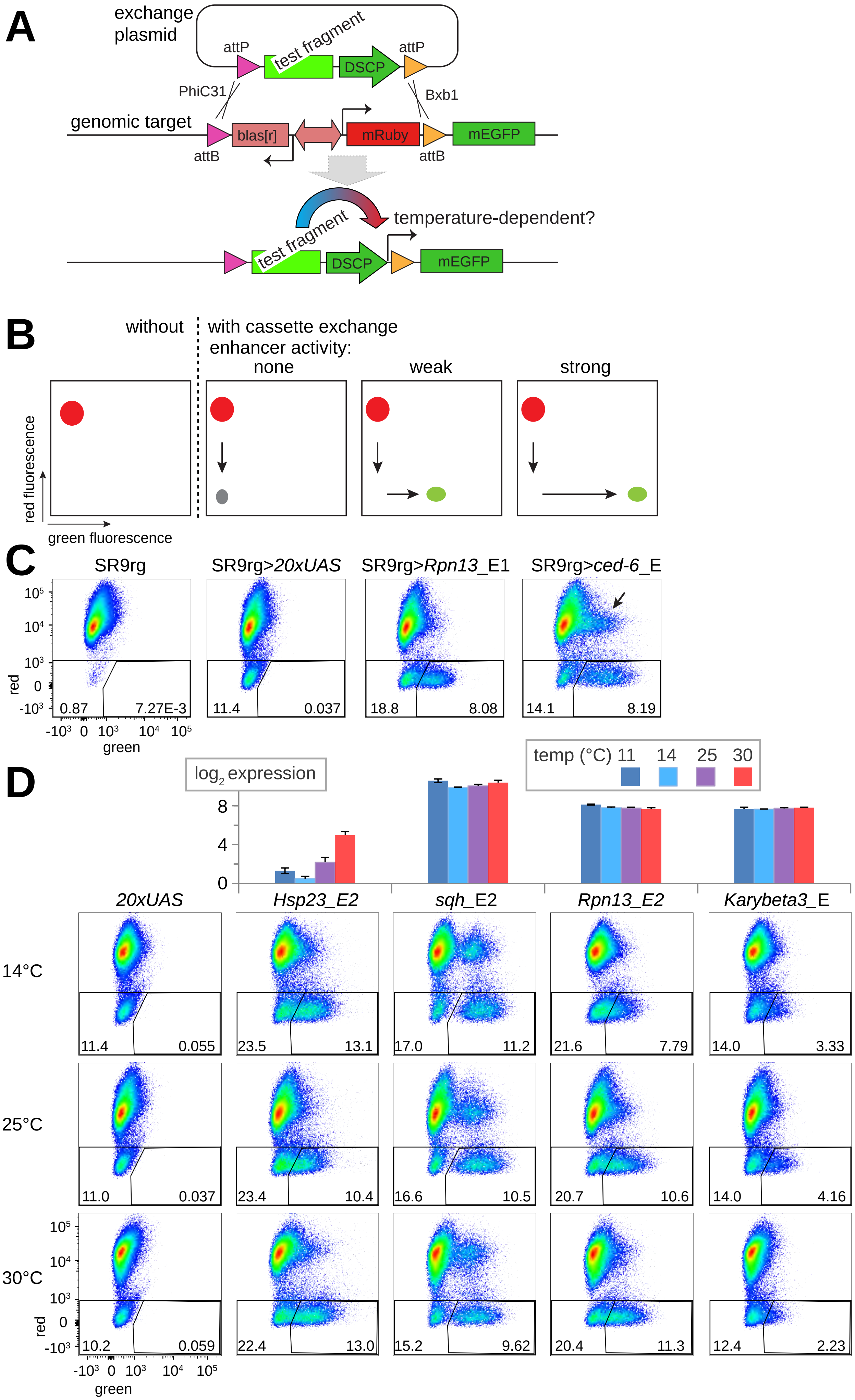
A
exchange plasmid
attP
test fragment
DSCP
attP
PhiC31
Bxb1
genomic target
blas[r]
mRuby
mEGFP
attB
attB
temperature-dependent?
test fragment
DSCP
mEGFP
B
without
with cassette exchange
enhancer activity:
none
weak
strong
red fluorescence
green fluorescence
C
SR9rg
SR9rg>20xUAS
SR9rg>Rpn13_E1
SR9rg>ced-6_E
red
green
0.87
7.27E-3
11.4
0.037
18.8
8.08
14.1
8.19
D
log2 expression
temp (°C) 11 14 25 30
20xUAS
Hsp23_E2
sqh_E2
Rpn13_E2
Karybeta3_E
14°C
25°C
30°C
11.4
0.055
23.5
13.1
17.0
11.2
21.6
7.79
14.0
3.33
11.0
0.037
23.4
10.4
16.6
10.5
20.7
10.6
14.0
4.16
10.2
0.059
22.4
13.0
15.2
9.62
20.4
11.3
12.4
2.23

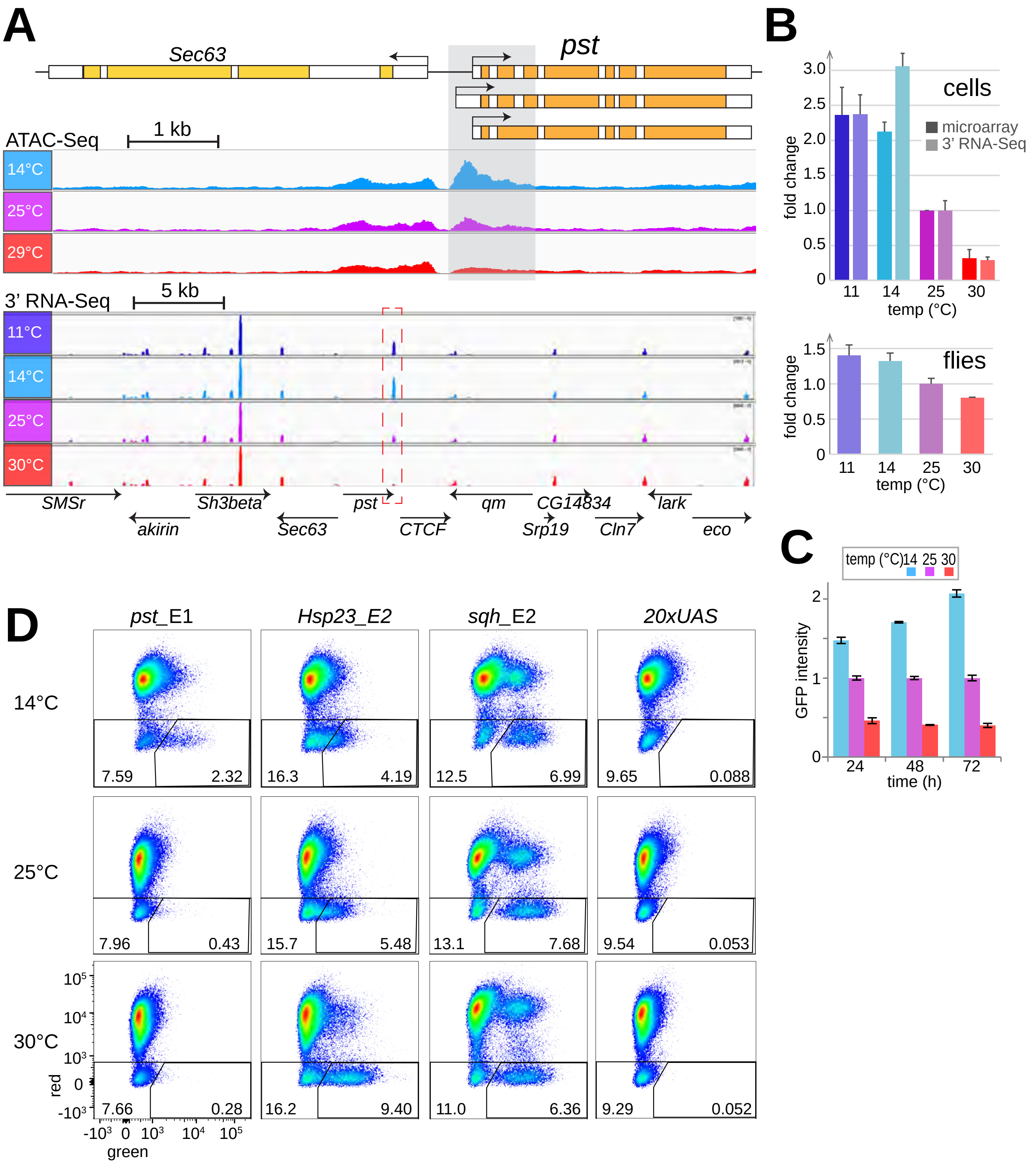
A
Sec63
pst
ATAC-Seq
1 kb
14°C
25°C
29°C
3' RNA-Seq
5 kb
11°C
14°C
25°C
30°C
SMSr
akirin
Sh3beta
Sec63
pst
CTCF
qm
CG14834
Srp19
Cln7
lark
eco
B
cells
microarray
3' RNA-Seq
fold change
temp (°C)
flies
C
temp (°C) 14 25 30
GFP intensity
time (h)
D
pst_E1
Hsp23_E2
sqh_E2
20xUAS
14°C
25°C
30°C
7.59
2.32
16.3
4.19
12.5
6.99
9.65
0.088
7.96
0.43
15.7
5.48
13.1
7.68
9.54
0.053
7.66
0.28
16.2
9.40
11.0
6.36
9.29
0.052
red
green

Bai et al., Fig. 7

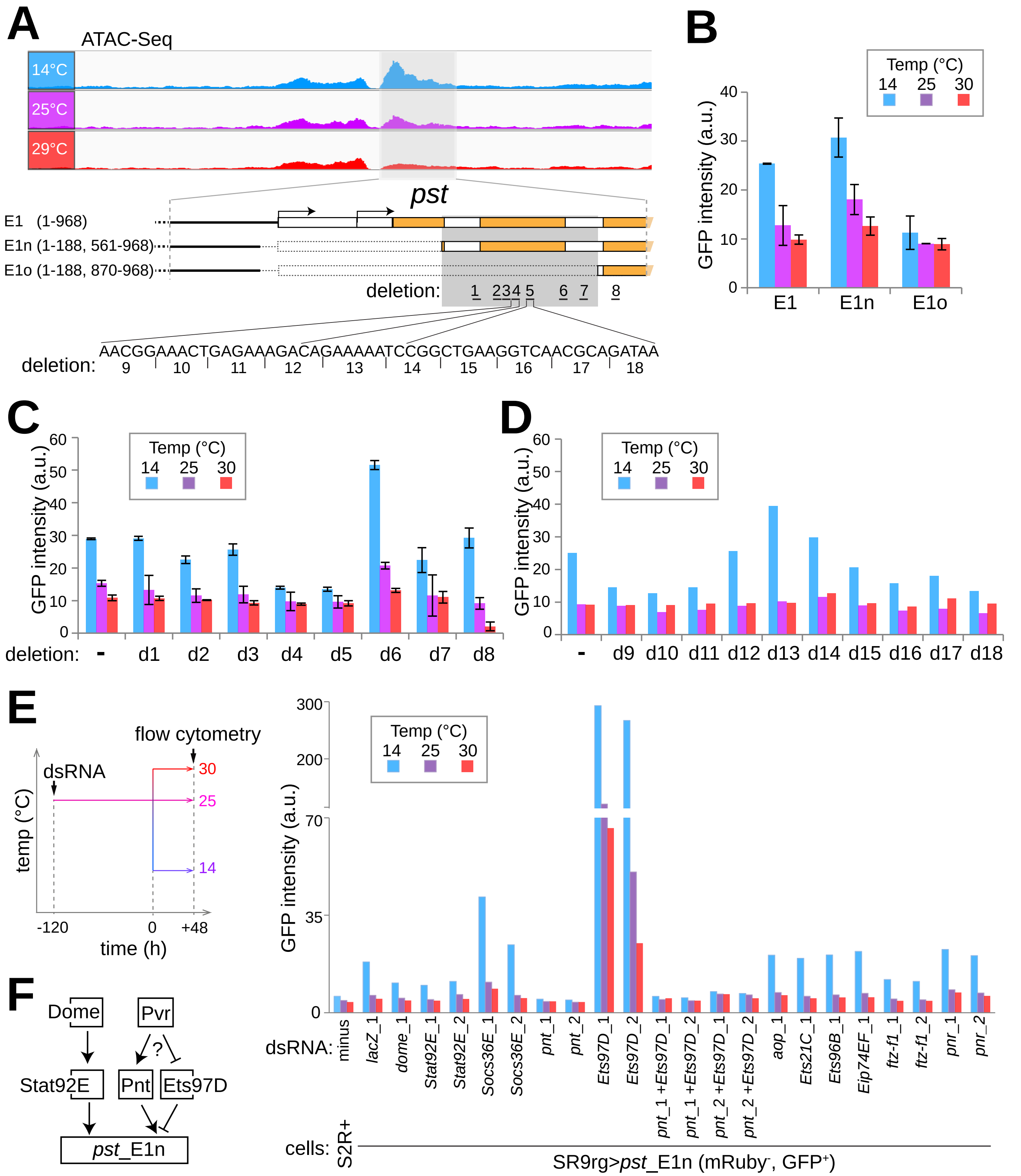

Bai et al., S1 Fig.

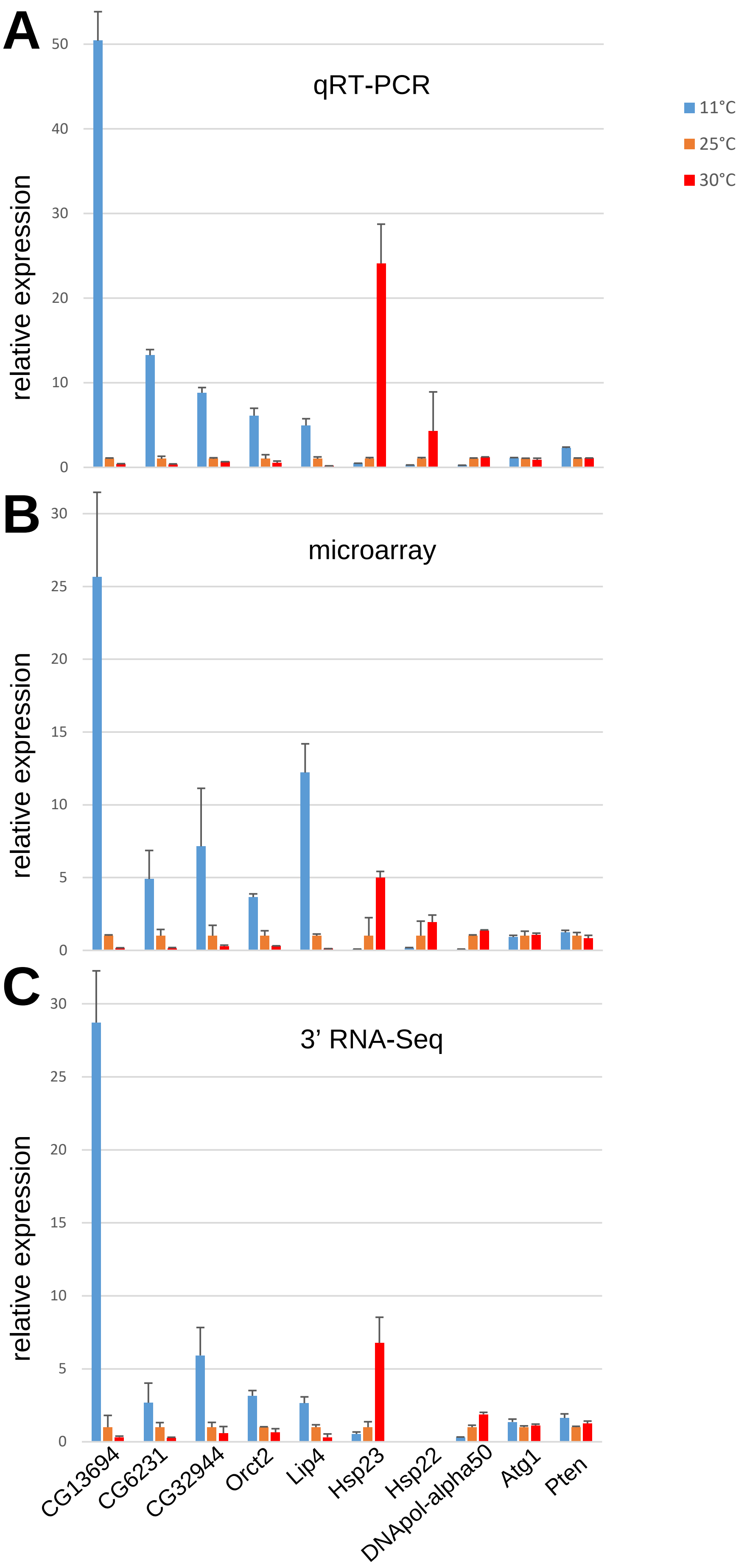

Bai et al., S2 Fig.

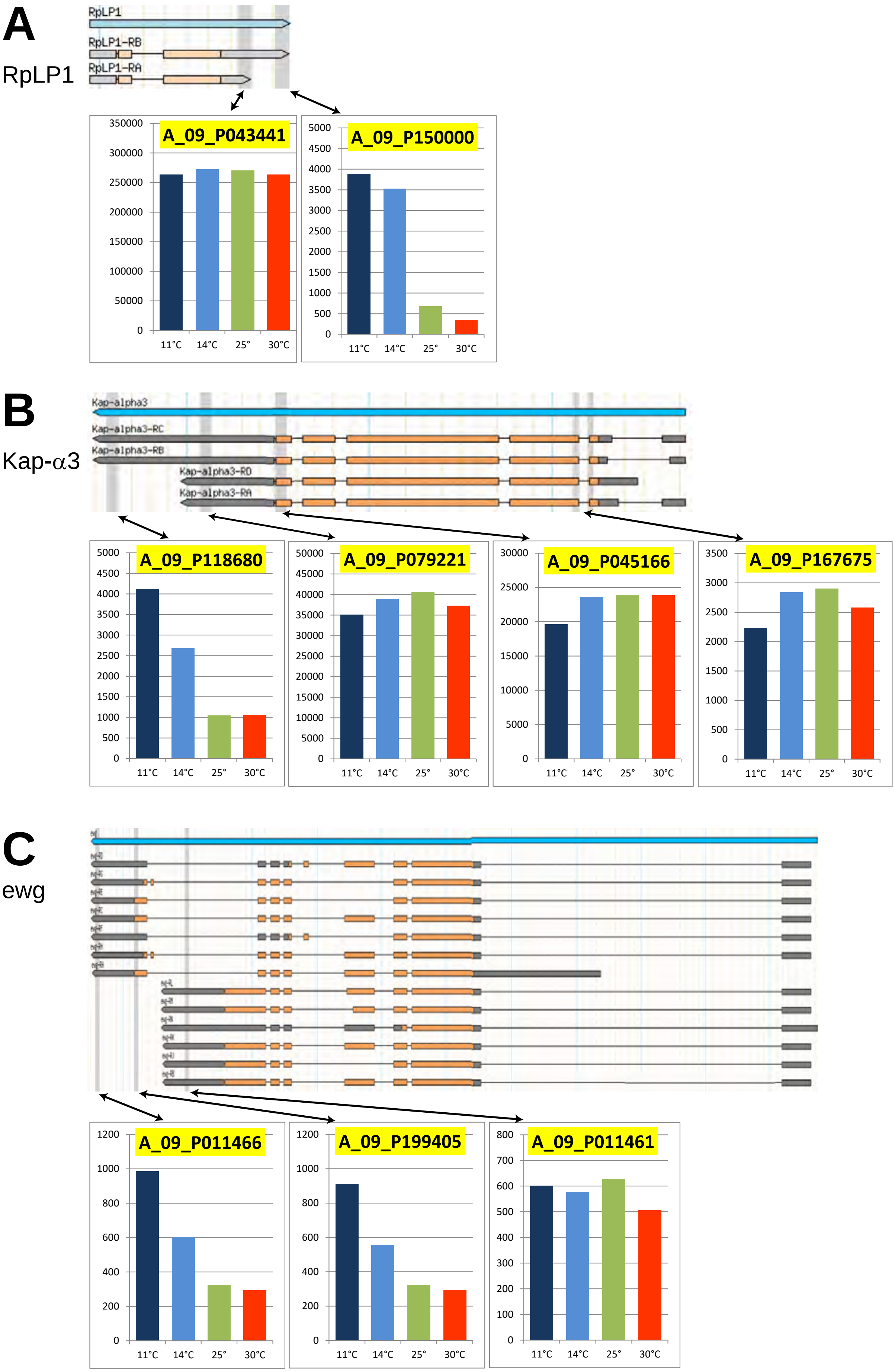

Bai et al., S3 Fig.

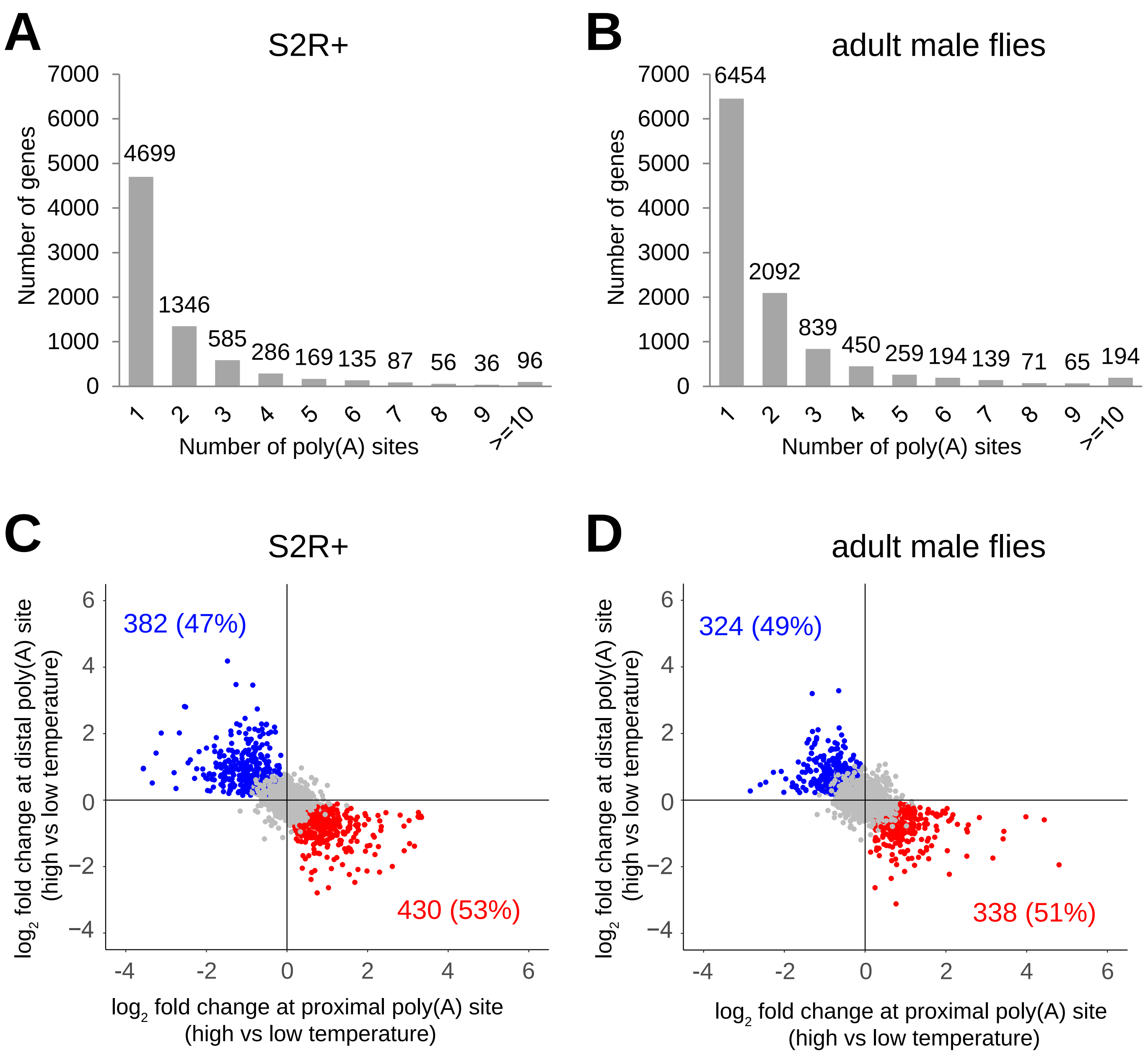

Bai et al., S4 Fig.

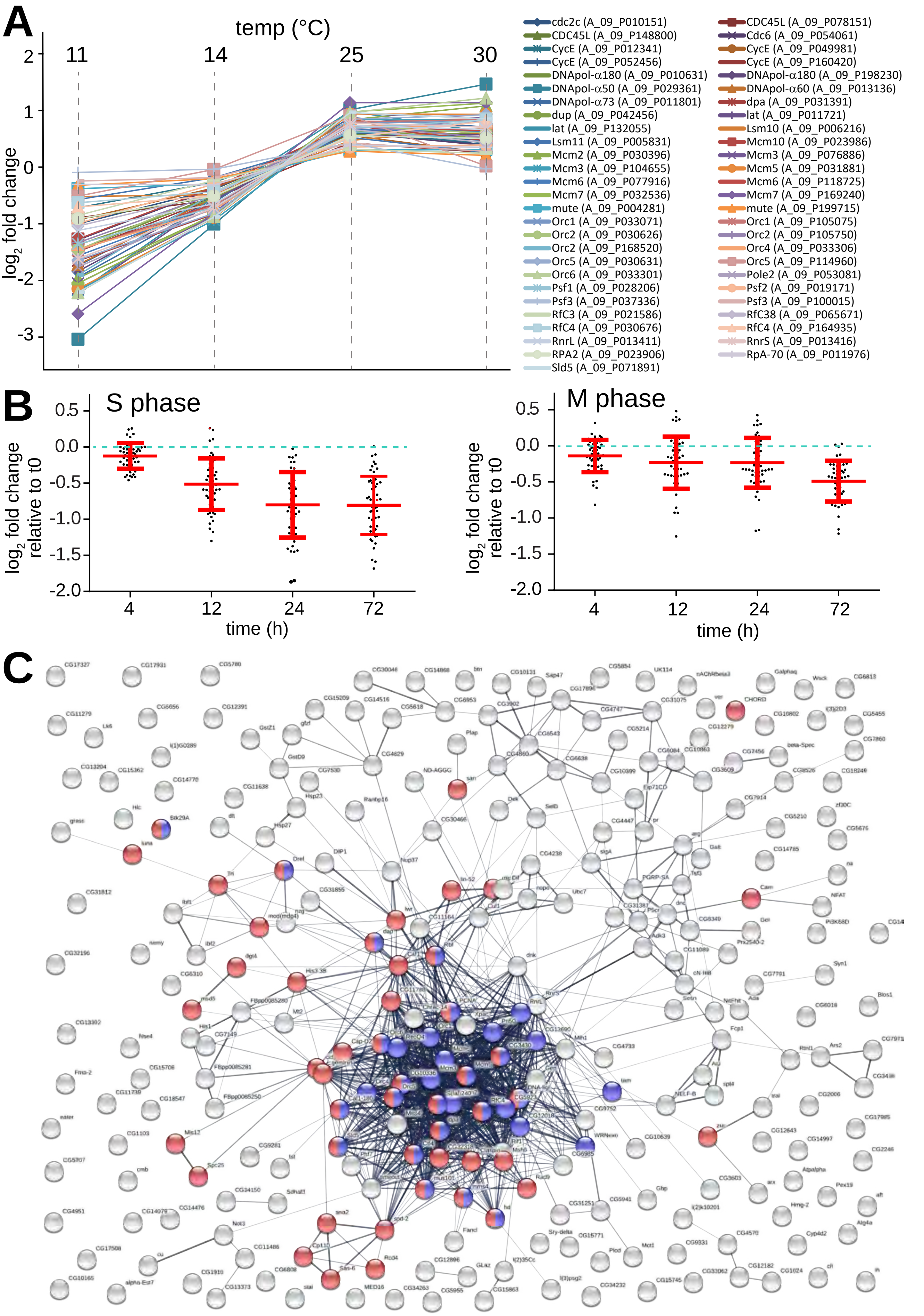

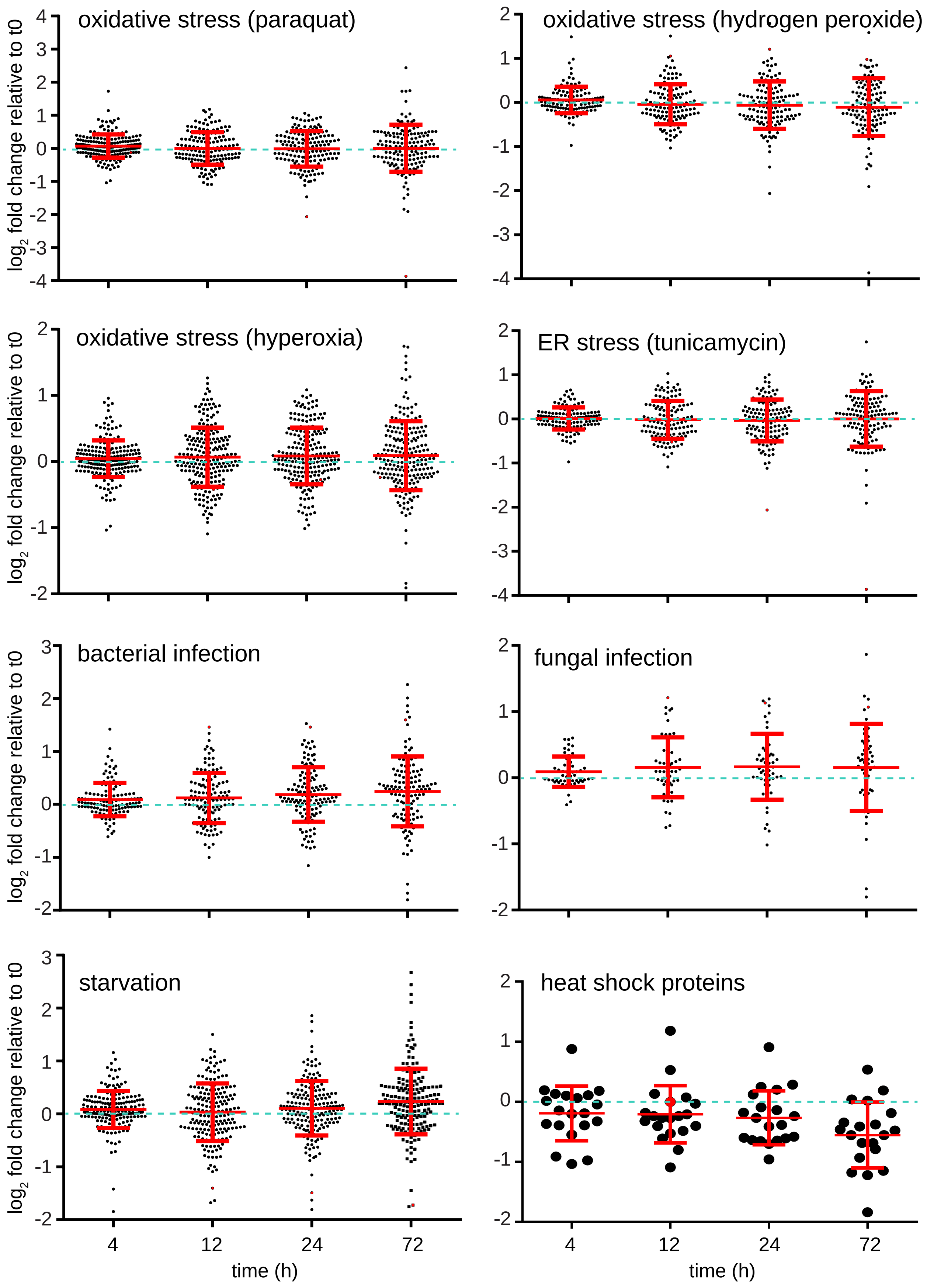
oxidative stress (paraquat)
oxidative stress (hydrogen peroxide)
oxidative stress (hyperoxia)
ER stress (tunicamycin)
bacterial infection
fungal infection
starvation
heat shock proteins
log2 fold change relative to t0
time (h)
4
12
24
72

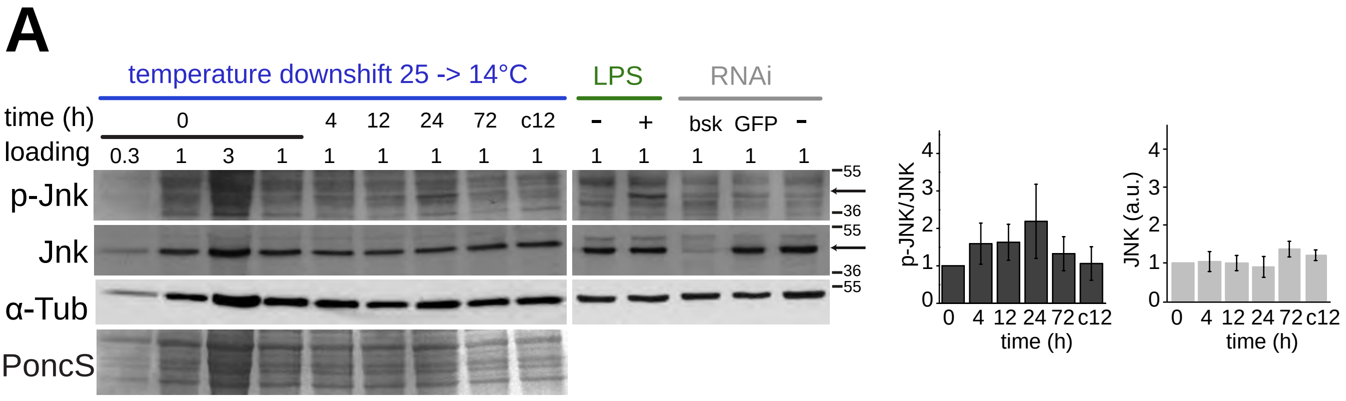
A
temperature downshift 25 -> 14°C
LPS
RNAi
time (h)
0
4
12
24
72
c12
-
+
bsk
GFP
-
loading
0.3
1
3
1
1
1
1
1
1
1
1
1
1
1
p-Jnk
Jnk
α-Tub
PoncS
55
36
55
36
55
p-JNK/JNK
JNK (a.u.)
time (h)
0 4 12 24 72 c12

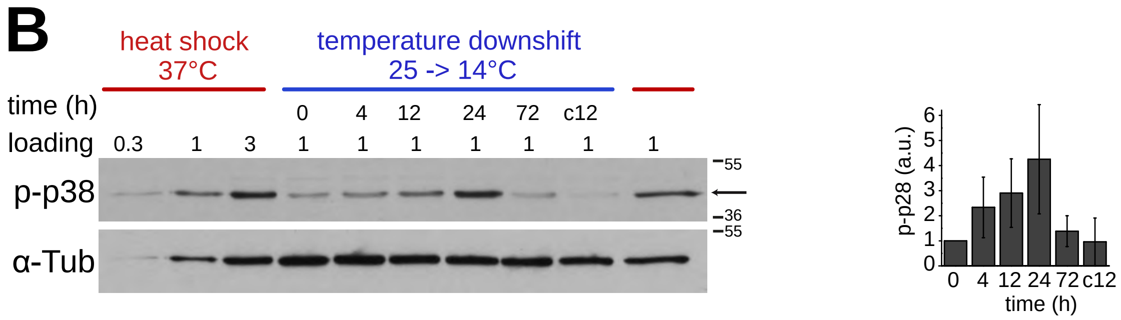
B
heat shock
37°C
temperature downshift
25 -> 14°C
time (h)
0
4
12
24
72
c12
loading
0.3
1
3
1
1
1
1
1
1
1
p-p38
α-Tub
55
36
55
p-p28 (a.u.)
time (h)
0 4 12 24 72 c12

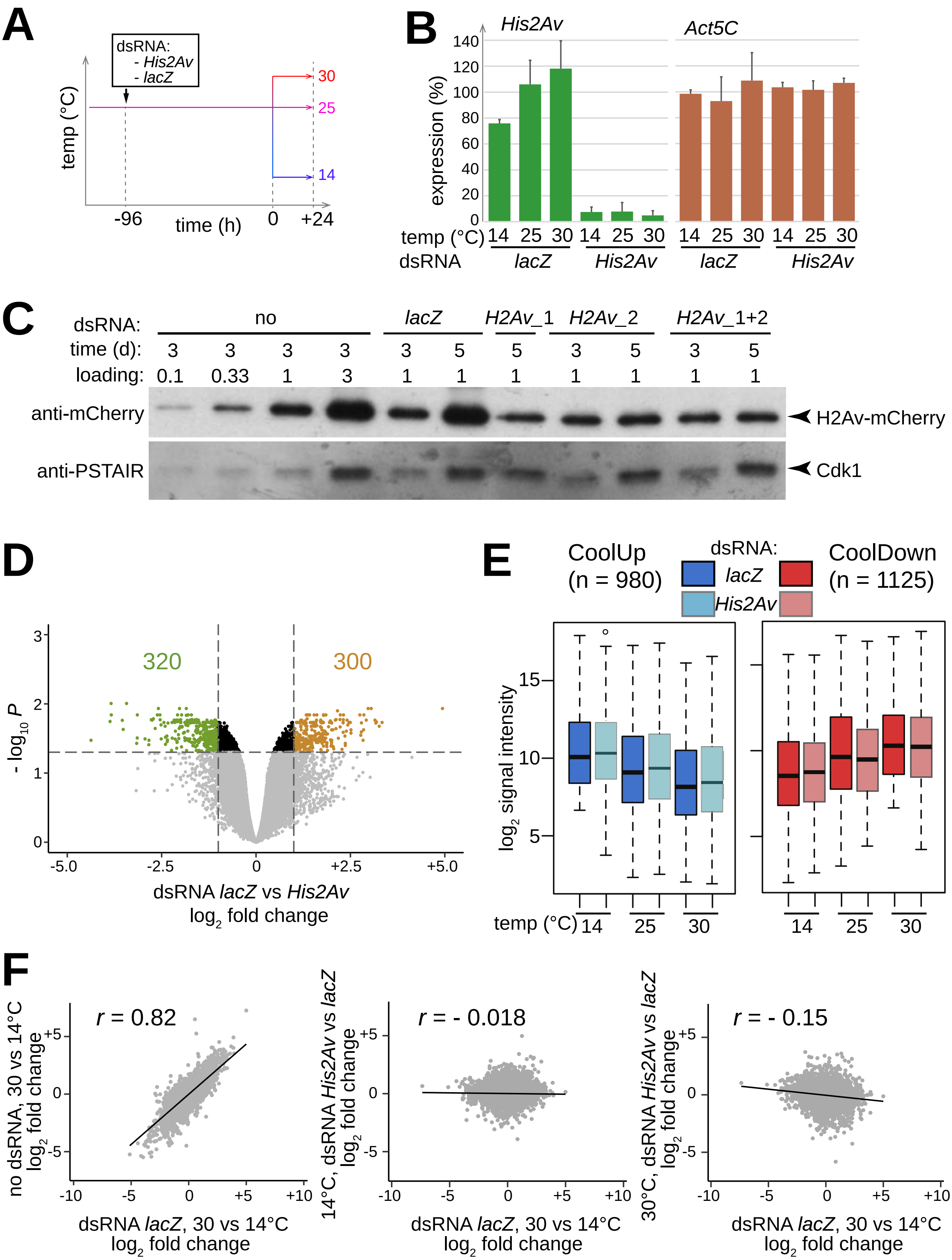
A
dsRNA:
- His2Av
- lacZ
temp (°C)
30
25
14
-96
time (h)
0
+24
B
His2Av
Act5C
expression (%)
140
120
100
80
60
40
20
0
temp (°C) 14 25 30 14 25 30 14 25 30 14 25 30
dsRNA lacZ His2Av lacZ His2Av
C
dsRNA: no lacZ H2Av_1 H2Av_2 H2Av_1+2
time (d): 3 3 3 3 3 5 5 3 5 3 5
loading: 0.1 0.33 1 3 1 1 1 1 1 1 1
anti-mCherry
H2Av-mCherry
anti-PSTAIR
Cdk1
D
320
300
- log10 P
dsRNA lacZ vs His2Av
log2 fold change
-5.0
-2.5
0
+2.5
+5.0
E
CoolUp
(n = 980)
dsRNA:
lacZ
His2Av
CoolDown
(n = 1125)
log2 signal intensity
15
10
5
temp (°C) 14 25 30
14 25 30
F
r = 0.82
no dsRNA, 30 vs 14°C
log2 fold change
dsRNA lacZ, 30 vs 14°C
log2 fold change
r = - 0.018
14°C, dsRNA His2Av vs lacZ
log2 fold change
r = - 0.15
30°C, dsRNA His2Av vs lacZ
log2 fold change
-10
-5
0
+5
+10

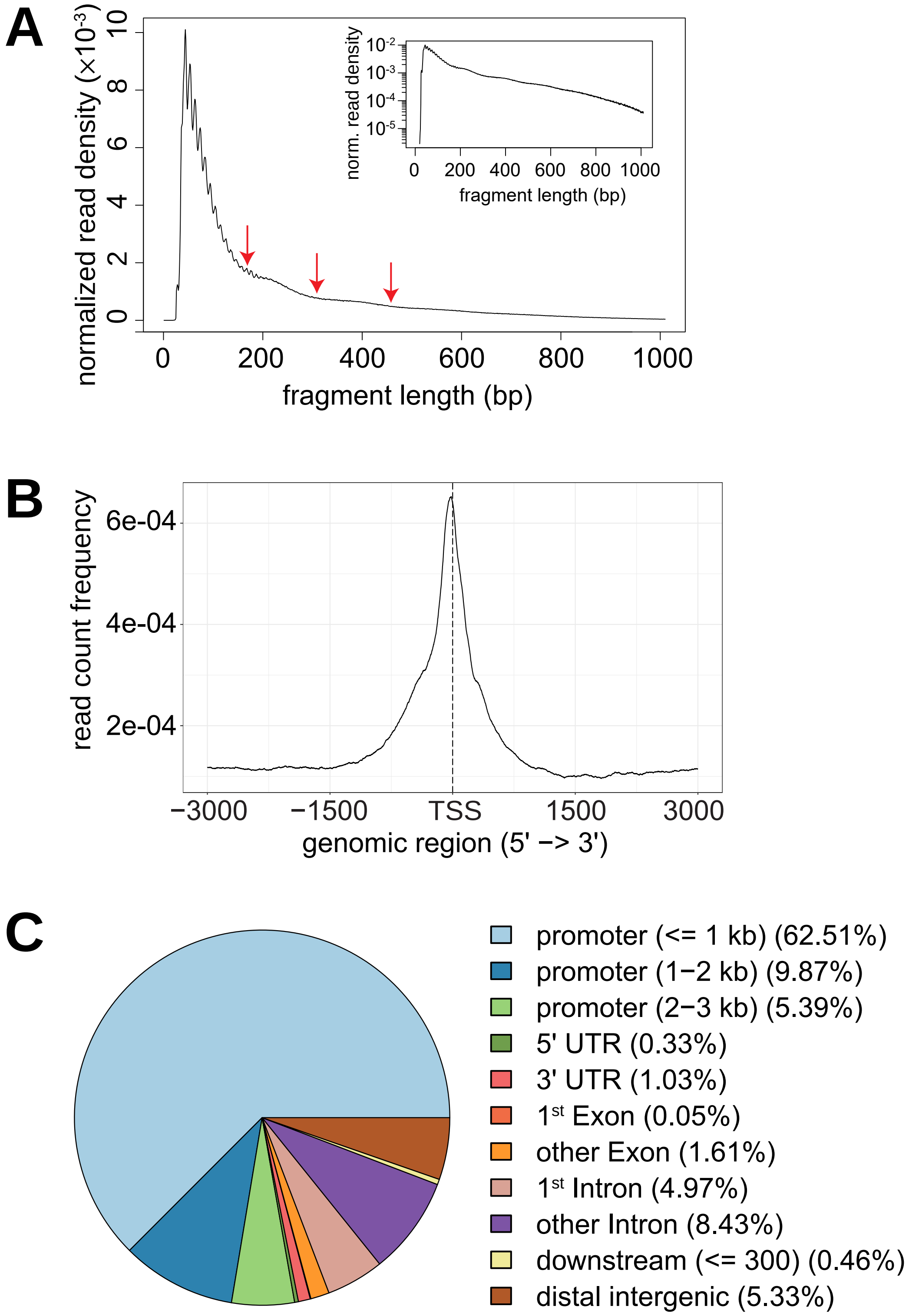
A
normalized read density (×10⁻³)
0
2
4
6
8
10
0
200
400
600
800
1000
fragment length (bp)
norm. read density
10⁻²
10⁻³
10⁻⁴
10⁻⁵
0
200
400
600
800
1000
fragment length (bp)
B
read count frequency
6e-04
4e-04
2e-04
−3000
−1500
TSS
1500
3000
genomic region (5' −> 3')
C
promoter (<= 1 kb) (62.51%)
promoter (1−2 kb) (9.87%)
promoter (2−3 kb) (5.39%)
5' UTR (0.33%)
3' UTR (1.03%)
1st Exon (0.05%)
other Exon (1.61%)
1st Intron (4.97%)
other Intron (8.43%)
downstream (<= 300) (0.46%)
distal intergenic (5.33%)

Bai et al., S9 Fig.

# A

| name | transcript level | test fragment[a)] | promoter[b)] | enhancer activity | temperature dependence [c)] |
|---|---|---|---|---|---|
| *Prx2540-1*_E | WarmUp | chr2R:10423096-10424598 | - | ++ | WarmUp |
| *Mcm2*_E | WarmUp | chr3R:8264489-8264947 | + | (+) | |
| *CG31075*_E_inverted | WarmUp | chr3R:26988420-26989034 | + | - | |
| *CG13694*_E | CoolUp | chr2L:116969-116021 | - | - | |
| *mthl14*_E | CoolUp | chr3L:245757-248102 | - | + | |
| *CG13893*_E | | chr3L:604500-600889 | - | + | |
| *CG6231*_E1 | CoolUp | chr3R:19622194-19619140 | - | ++ | WarmUp |
| *CG6231*_E2 | CoolUp | chr3R:19630067-19627647 | + | - | |
| | CoolUp | | - | - | |
| *CG6231*_E3 | CoolUp | chr3R:19633419-19632370 | + | - | |
| | CoolUp | | - | - | |
| *Kah*_E1 | CoolUp | chr3L:590671-587637 | + | - | |
| *Kah*_E2 | CoolUp | chr3L:589326-588826 | + | (+) | |
| *Orct2*_E | CoolUp | chr3R:24271000-24273602 | + | - | |
| *Lip4*_E | CoolUp | chr2L:10527000-10529600 | + | (+) | |
| | CoolUp | | - | - | |

| | | | | | |
|---|---|---|---|---|---|
| *Lip4*_E_inverted | CoolUp | chr2L:10529600-10527000 | + | - | |
| *cv-2*_E1 | CoolUp | chr2R:21369718-21367280 | + | + | (CoolUp) |
| *cv-2*_E2 | CoolUp | chr2R:21383738-21378813 | + | +++ | |
| *Pask*_E | CoolUp | chr2R:23987549-23989100 | - | + | CoolUp |
| *pirk*_E | CoolUp | chr2R:21663200-21661544 | - | - | |
| *mthl5*_E | CoolUp | chr3R:11888215-11886762 | - | +++ | (WarmUp) |
| *betaTub97E*F_E1_inverted | CoolUp | chr3R:27986500-27990003 | + | - | |
| *betaTub97EF*_E2 | CoolUp | chr3R:27990000-27988130 | - | - | |
| *CR45607/upd2*_E | CoolUp | chrX:18246571-18250087 | + | - | |
| *Socs36E*_E1 | CoolUp | chr2L:18146300-18142749 | + | - | |
| *Socs36E*_E2 | CoolUp | chr2L:18142600-18141068 | + | - | |
| *GILT2*_E | (CoolUp) | chr3R:23678729-23679439 | + | - | |
| | (CoolUp) | | - | (+) | |
| *Atf3*_E | | chrX:1249873-1248024 | + | (+) | |
| *CG11403*_E | | chrX:1243130-1239626 | + | - | |
| *CG32809*_E | | chrX:1692228-1691362 | + | - | |
| *ACC*_E | | chr2R:7992450-7991766 | + | (+) | |
| | | | - | - | |
| *pain*_E | | chr2R:24920931-24920217 | + | + | |
| *Rho1*_E | | chr2R:16106015-16106899 | + | + | |
| *CG43149*_E | | chr3L:56221-55198 | + | + | |

| | | | | | |
|---|---|---|---|---|---|
| *CG6888*_E | | chr3L:15205814-15207085 | + | (+) | |
| *CG14810*_E | | chrX:1821507-1820886 | + | - | |
| *CG15482*_E | | chr2L:13211885-13211259 | + | - | |
| *CR45693*_E | | chr2L:14516828-14514926 | + | + | |
| *CR43263*_E | | chr2L:1218362-1217849 | + | + | |
| *RNaseX25*_E1 | | chr3L:7969603-7970384 | + | +++ | CoolUp |
| *RNaseX25*_E2 | | chr3L:7968618-7970699 | + | ++ | CoolUp |
| | | | - | + | |
| *RNaseX25*_E3 | | chr3L:7968618-7970384 | + | ++ | CoolUp |
| *RNaseX25*_E4 | | chr3L:7969603-7970699 | + | ++ | |
| | | | - | ++ | |

a) chromosomal coordinates of the tested DNA fragments according to genome release dm6.
b) some candidate fragments, which included an endogenous promoter region, were inserted into an exchange plasmid lacking the DSCP. Accordingly, the presence (+) or absence (-) of the DSCP in the exchange plasmid is indicated.
c) the effect of temperature on enhancer activity is specified.

# B

Bai et al., S9 Fig.

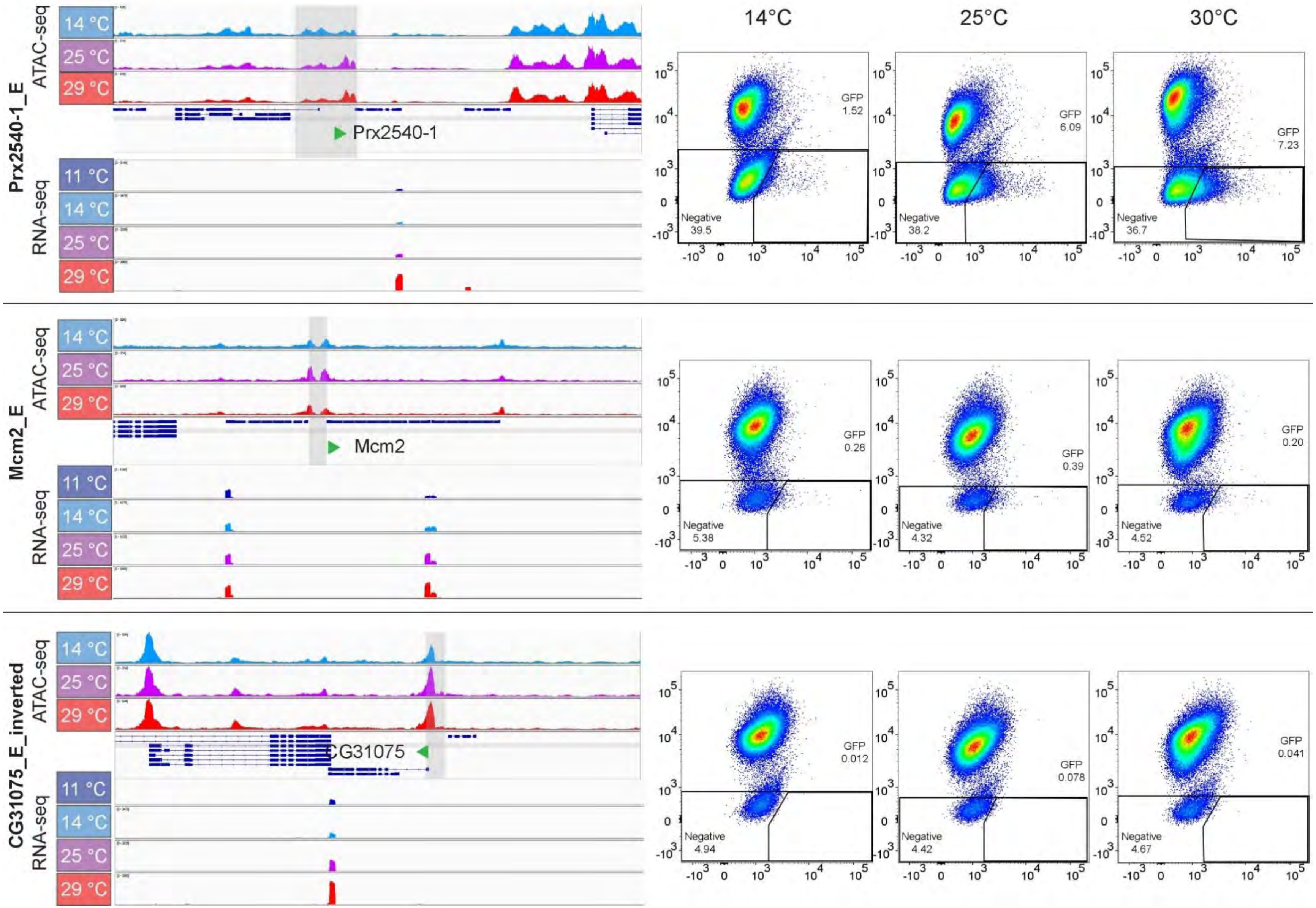

Bai et al., S9 Fig.

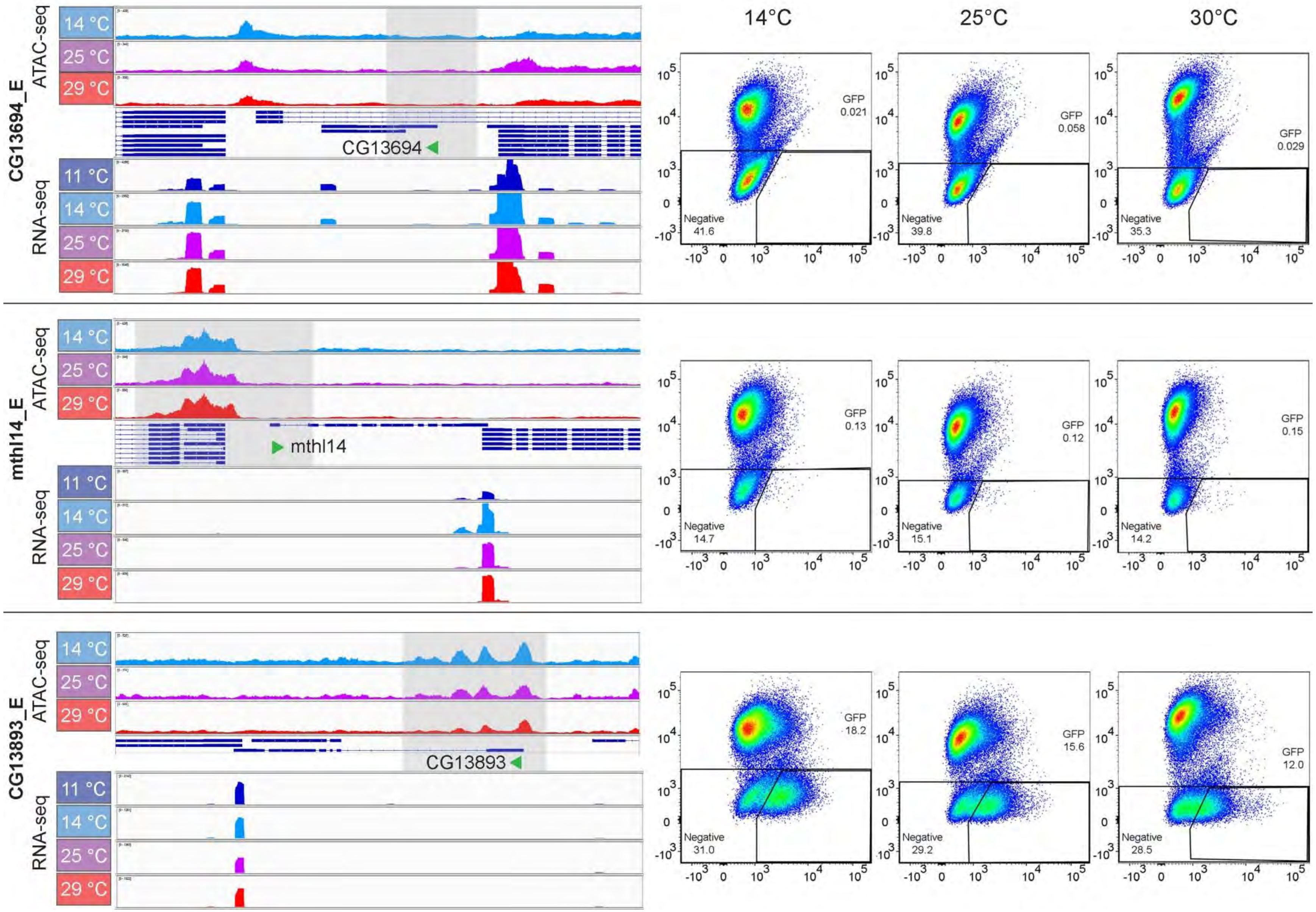

Bai et al., S9 Fig.

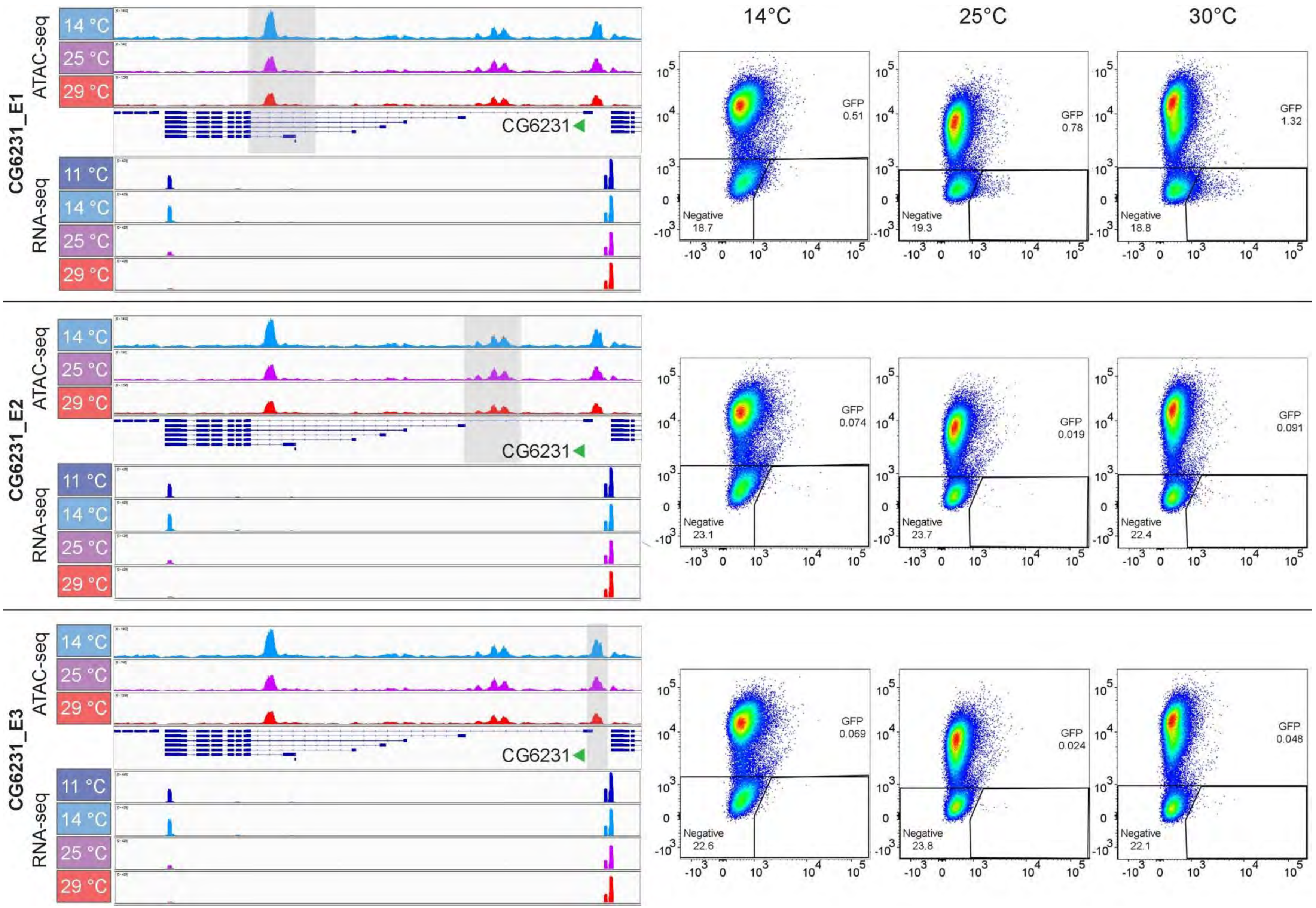

Bai et al., S9 Fig.

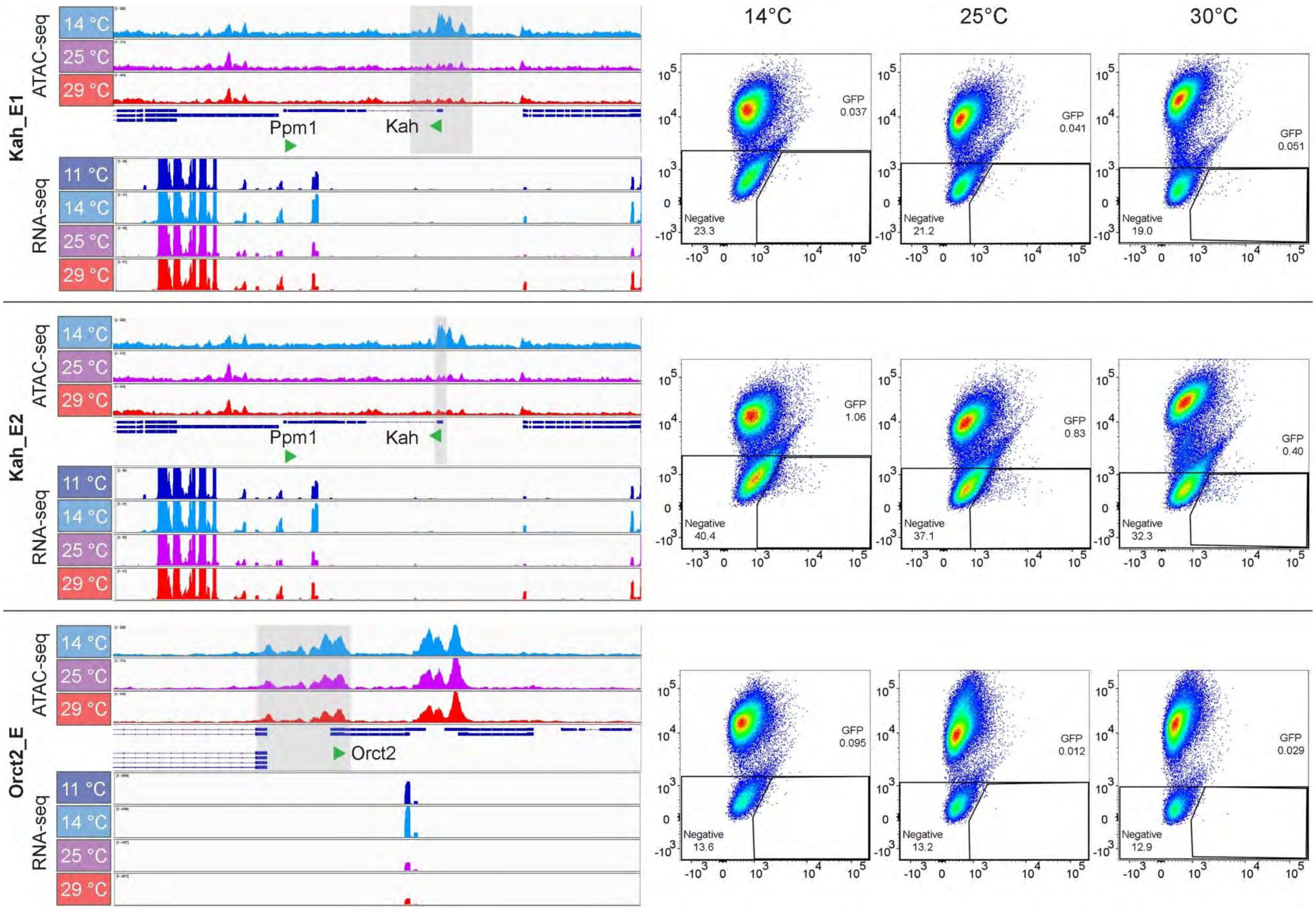

Bai et al., S9 Fig.

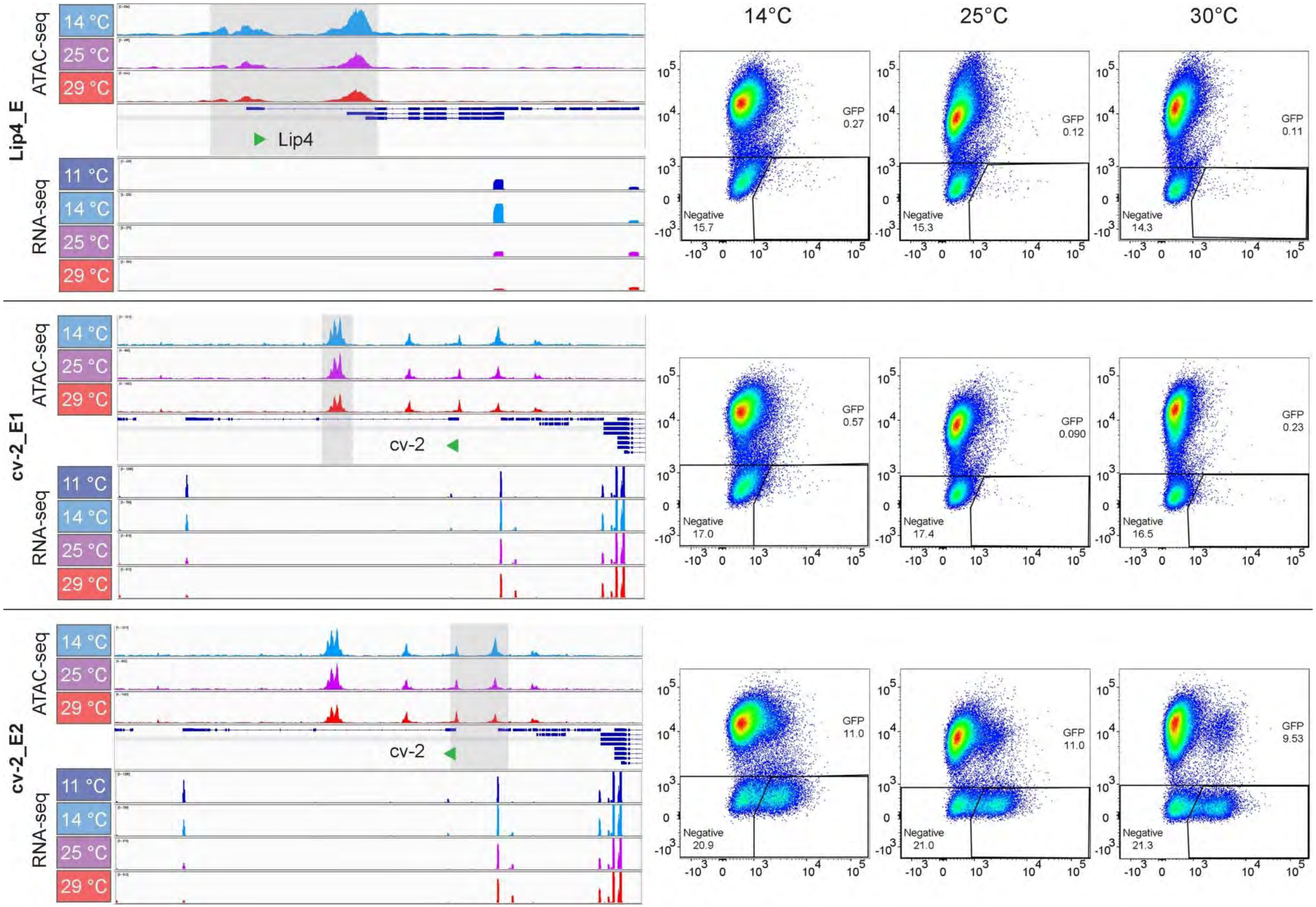

Bai et al., S9 Fig.

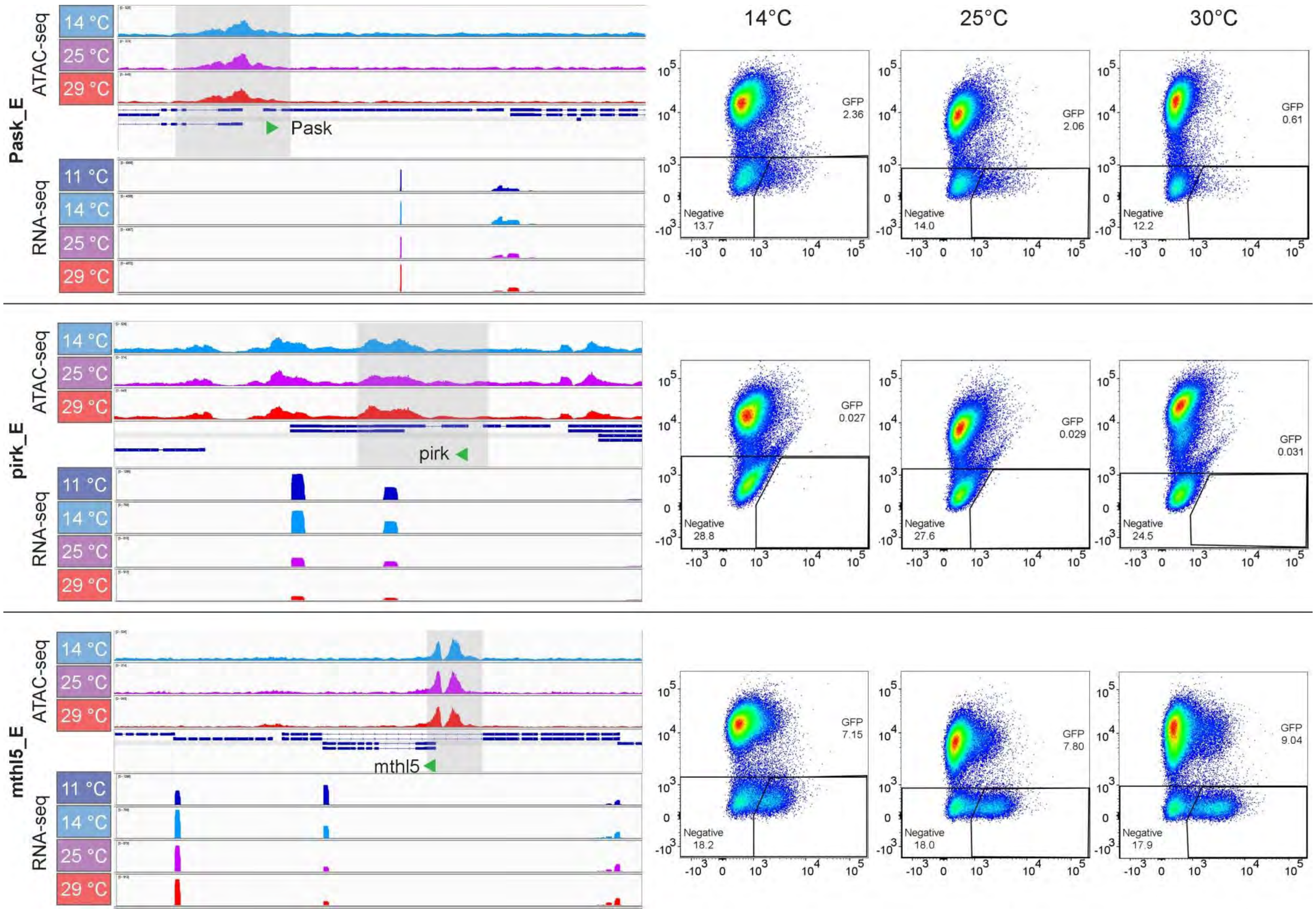

Bai et al., S9 Fig.

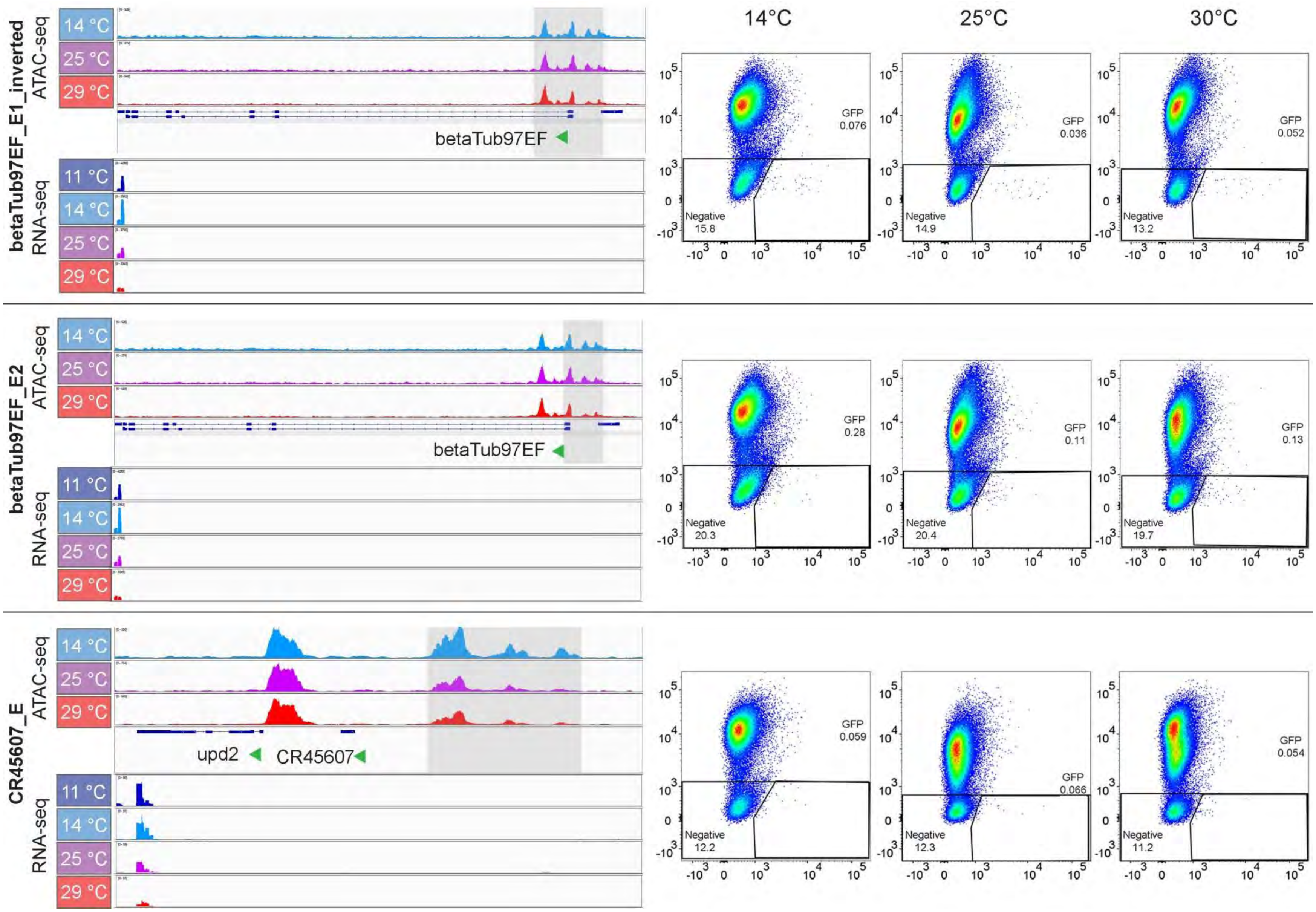

Bai et al., S9 Fig.

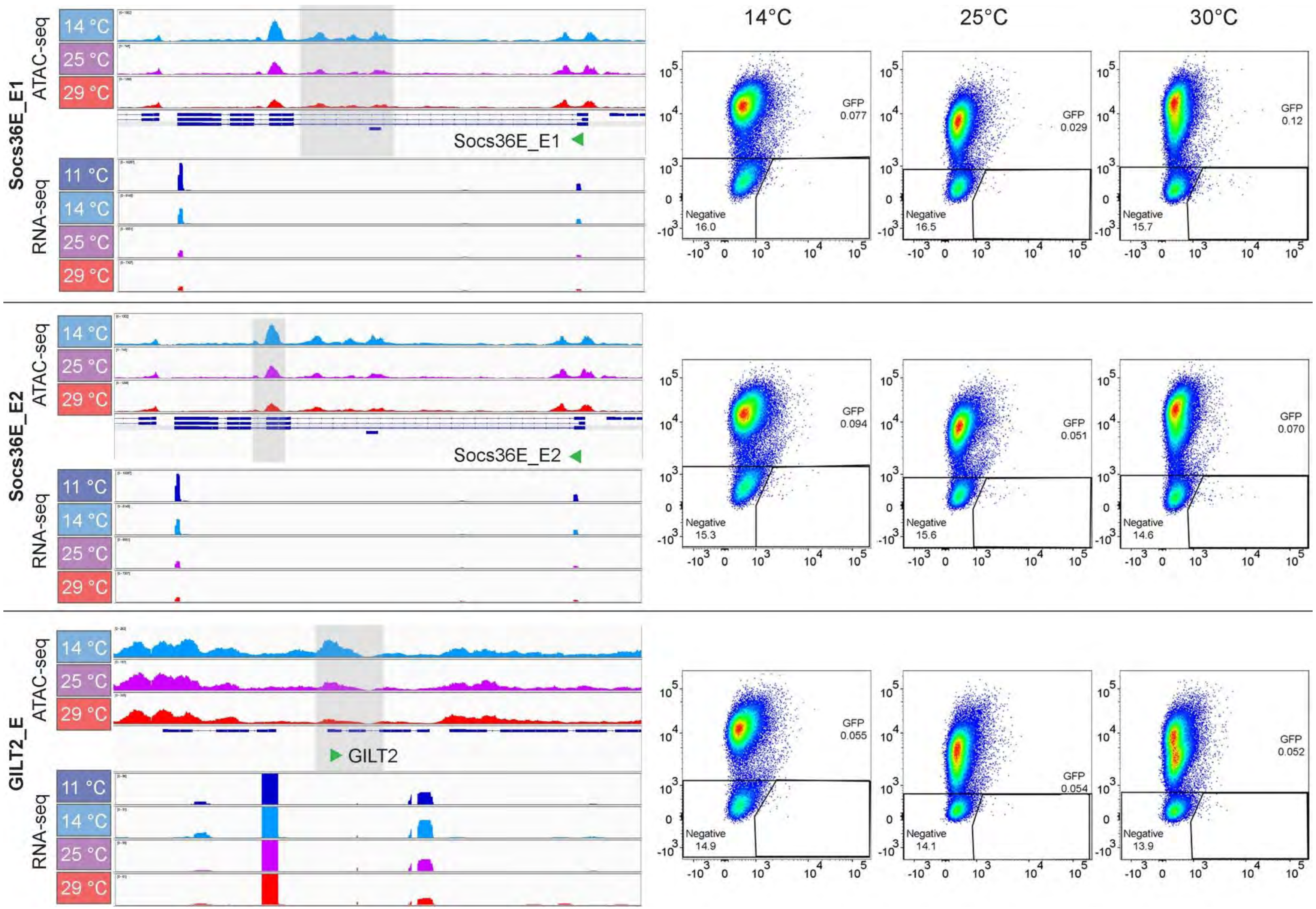

Bai et al., S9 Fig.

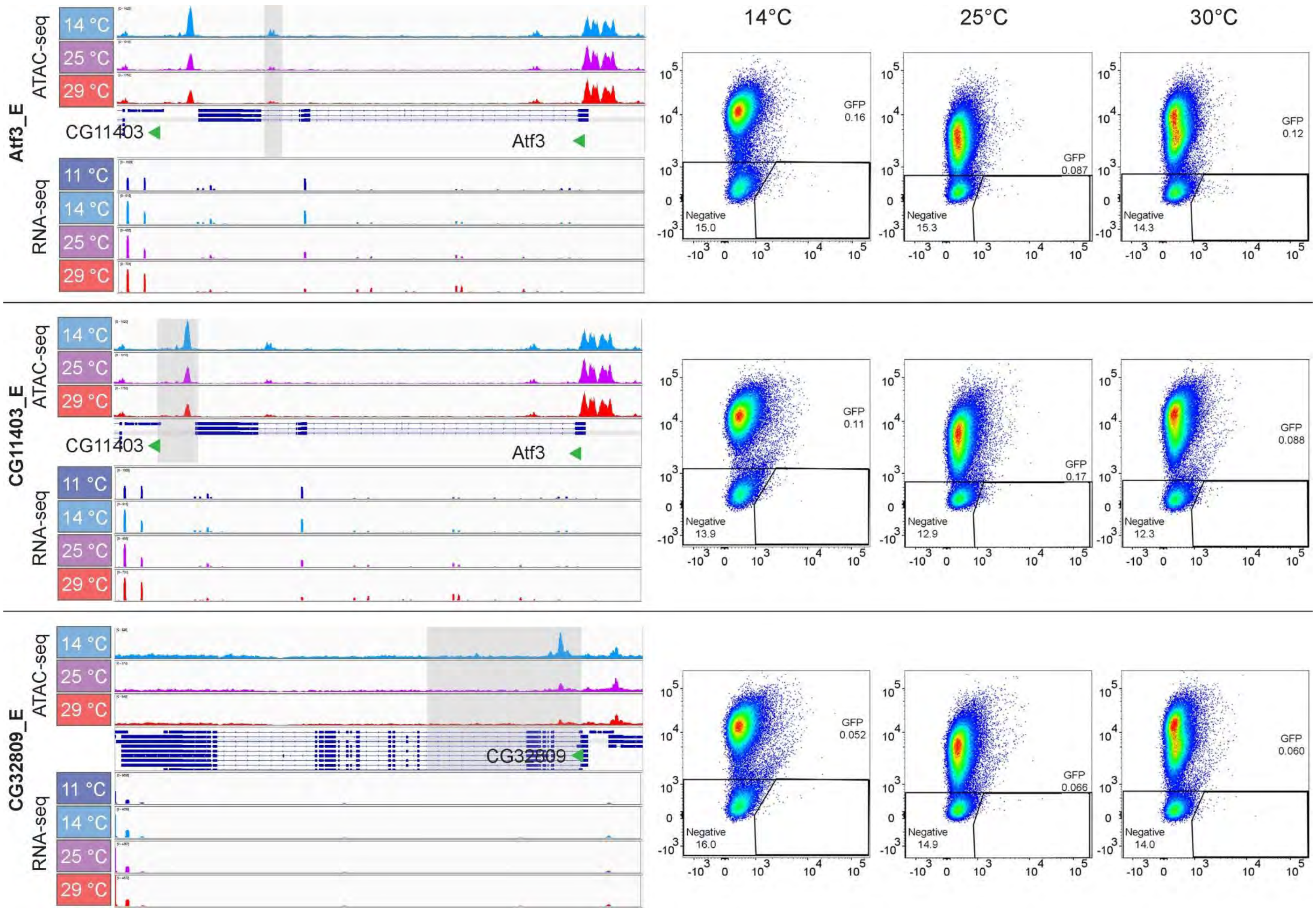

Bai et al., S9 Fig.

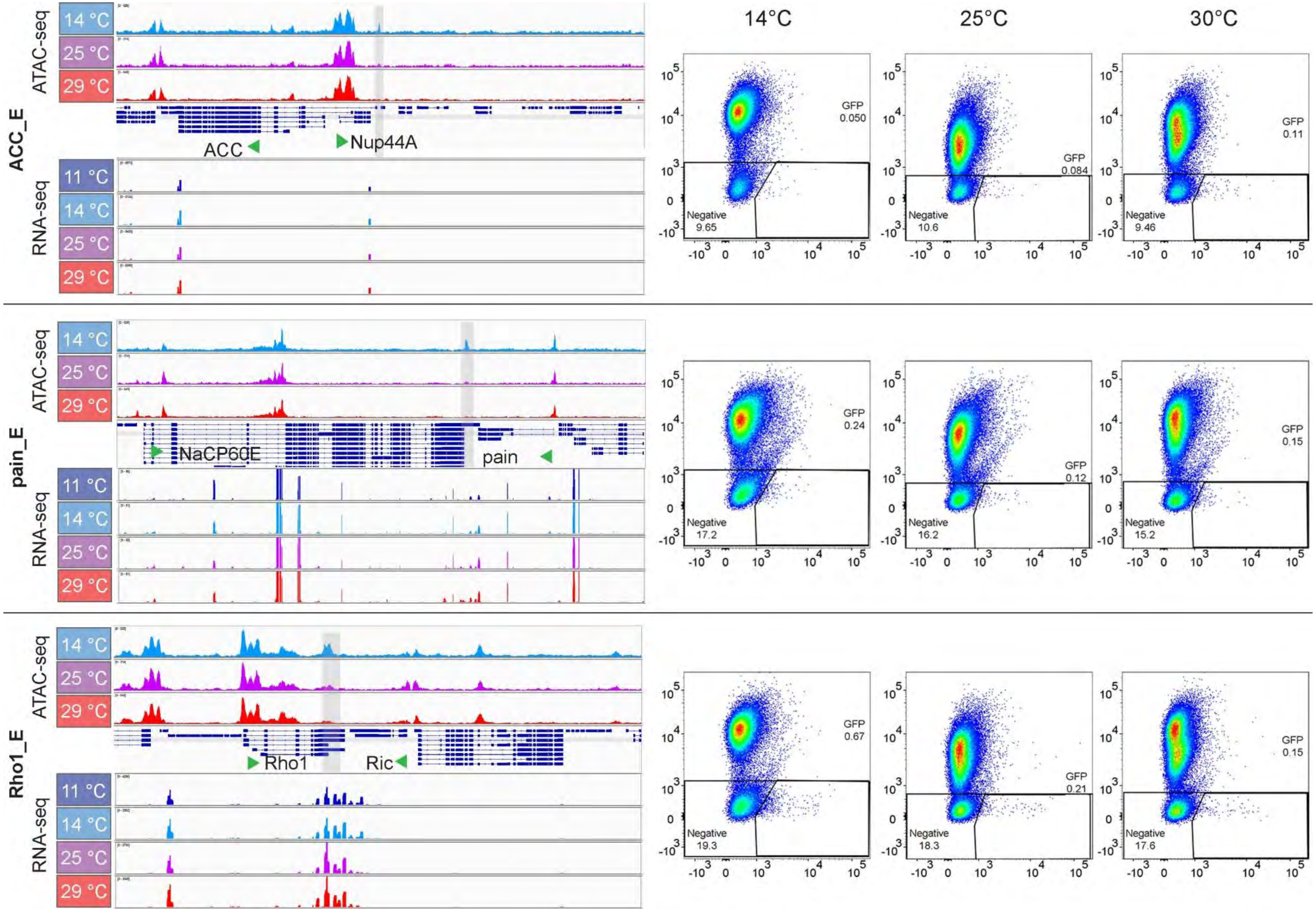

Bai et al., S9 Fig.

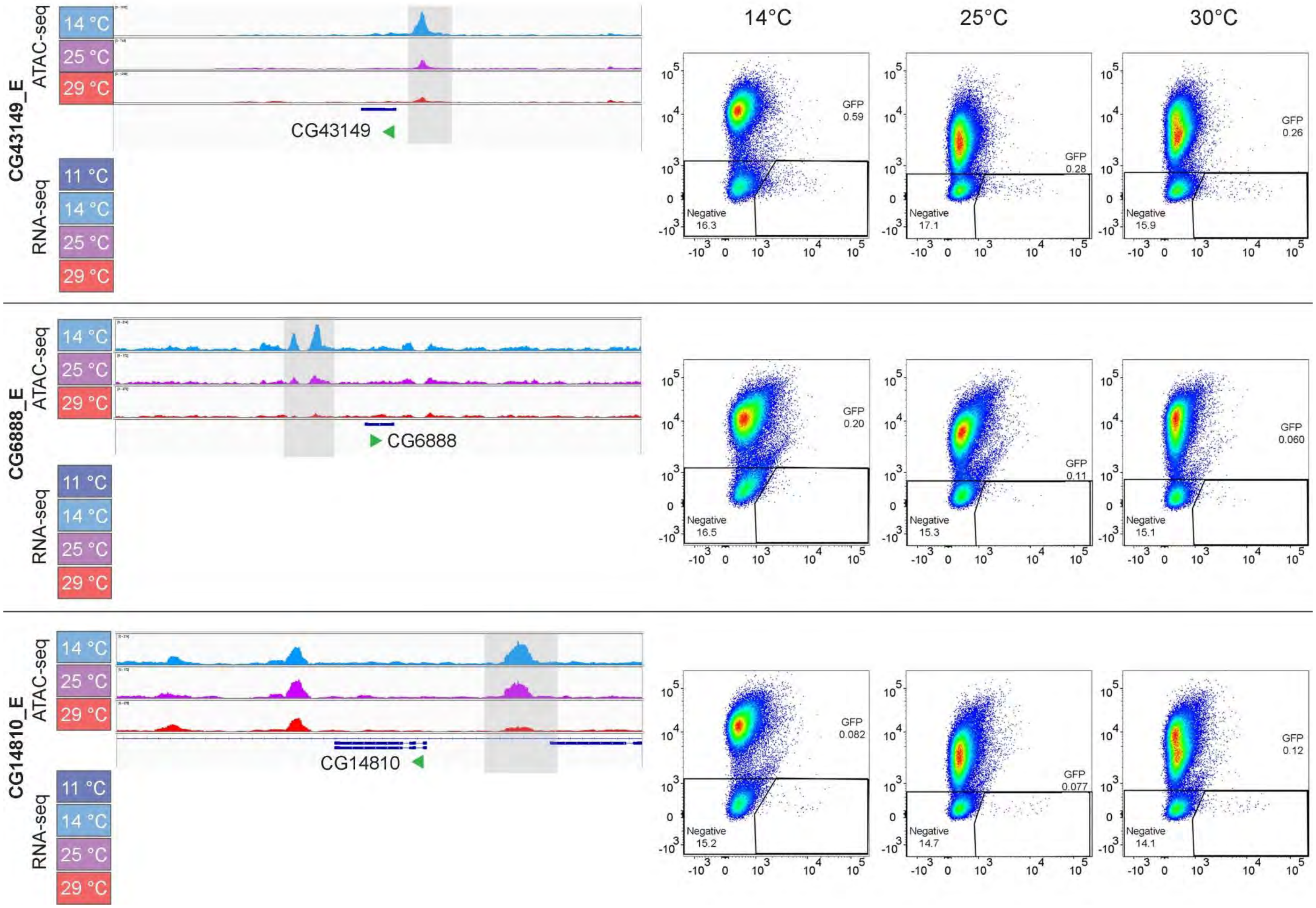

Bai et al., S9 Fig.

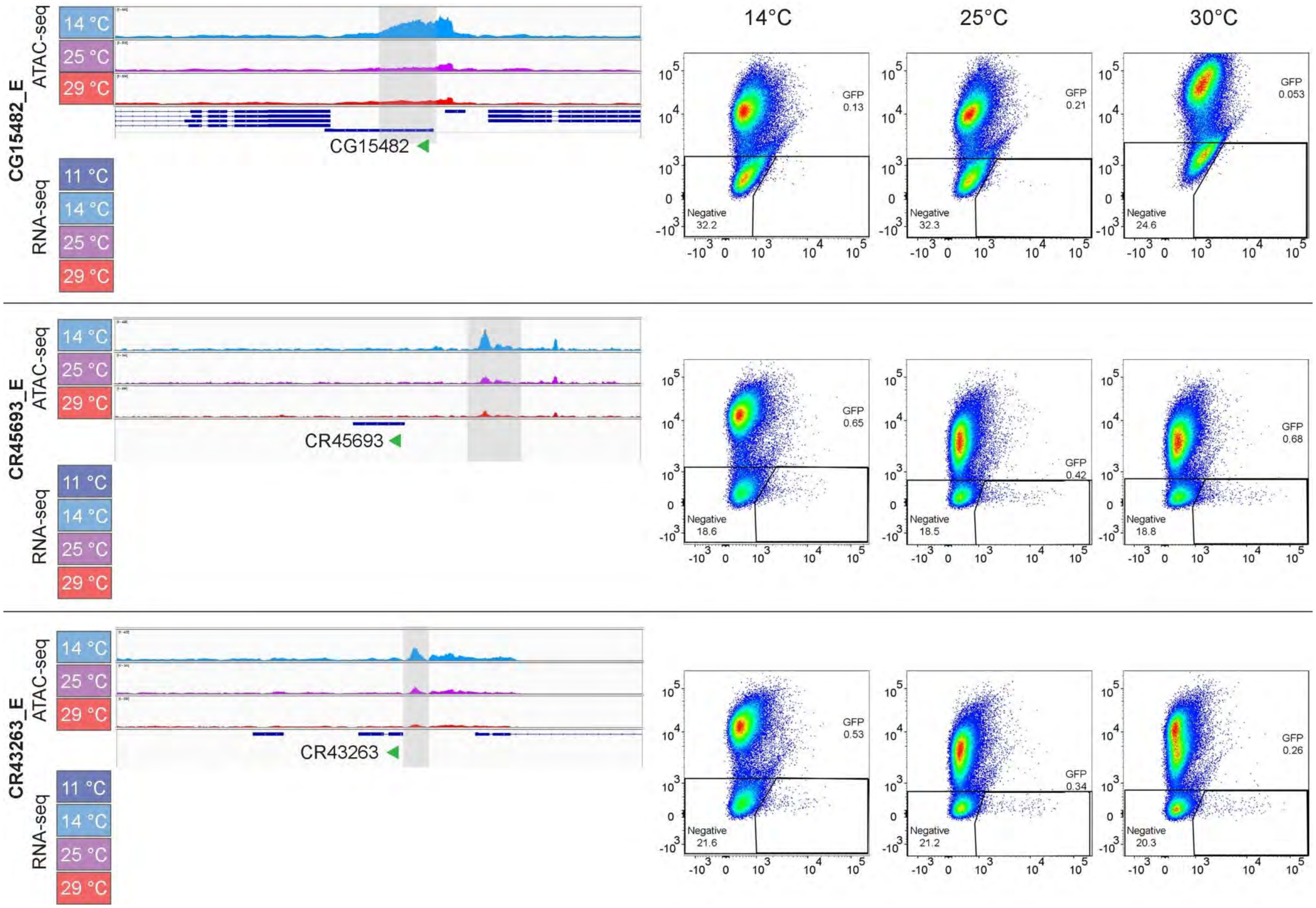

Bai et al., S9 Fig.

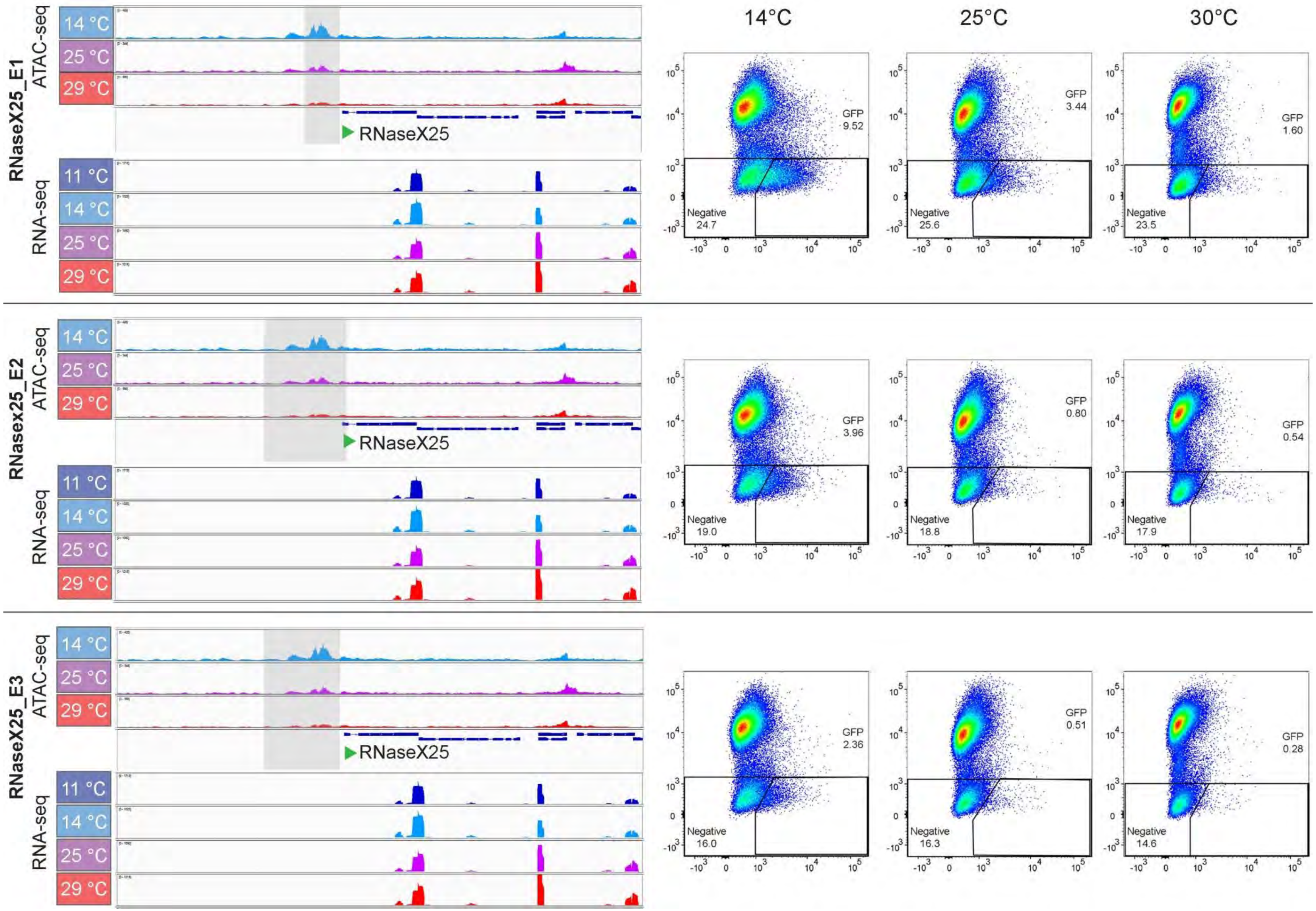

Bai et al., S9 Fig.

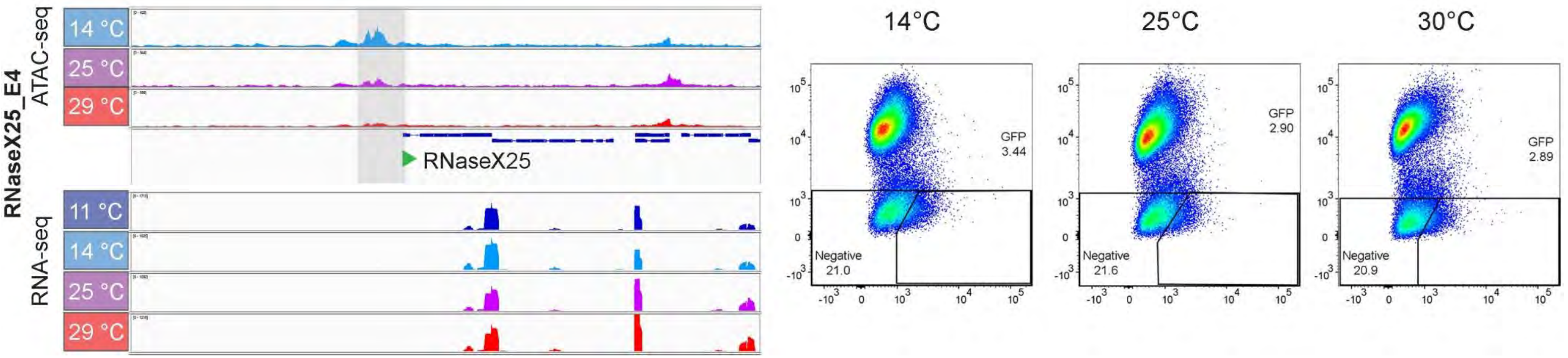


**S9 Fig. Assay for temperature-dependent enhancer activity with candidate CREs.**
(**A**) Table summarizing location and activity of candidate CREs analyzed after RMCE with SR9rg cells.
(**B**) Data for the analyzed candidate CREs. Candidate CREs (designation in bold on the left side) were selected based on data obtained by ATAC-seq (top three browser tracks) and 3' RNA-seq (bottom three browser tracks). Grey shading represents the region of the analyzed DNA fragment. Green arrowheads indicate the direction of transcription of the proximal gene. Scatter plots on the right side present green (x axis) and red (y axis) fluorescence intensities as determined by flow cytometry with SR9rg cells after RMCE and incubation at the indicated temperatures.

Bai et al., S10 Fig.

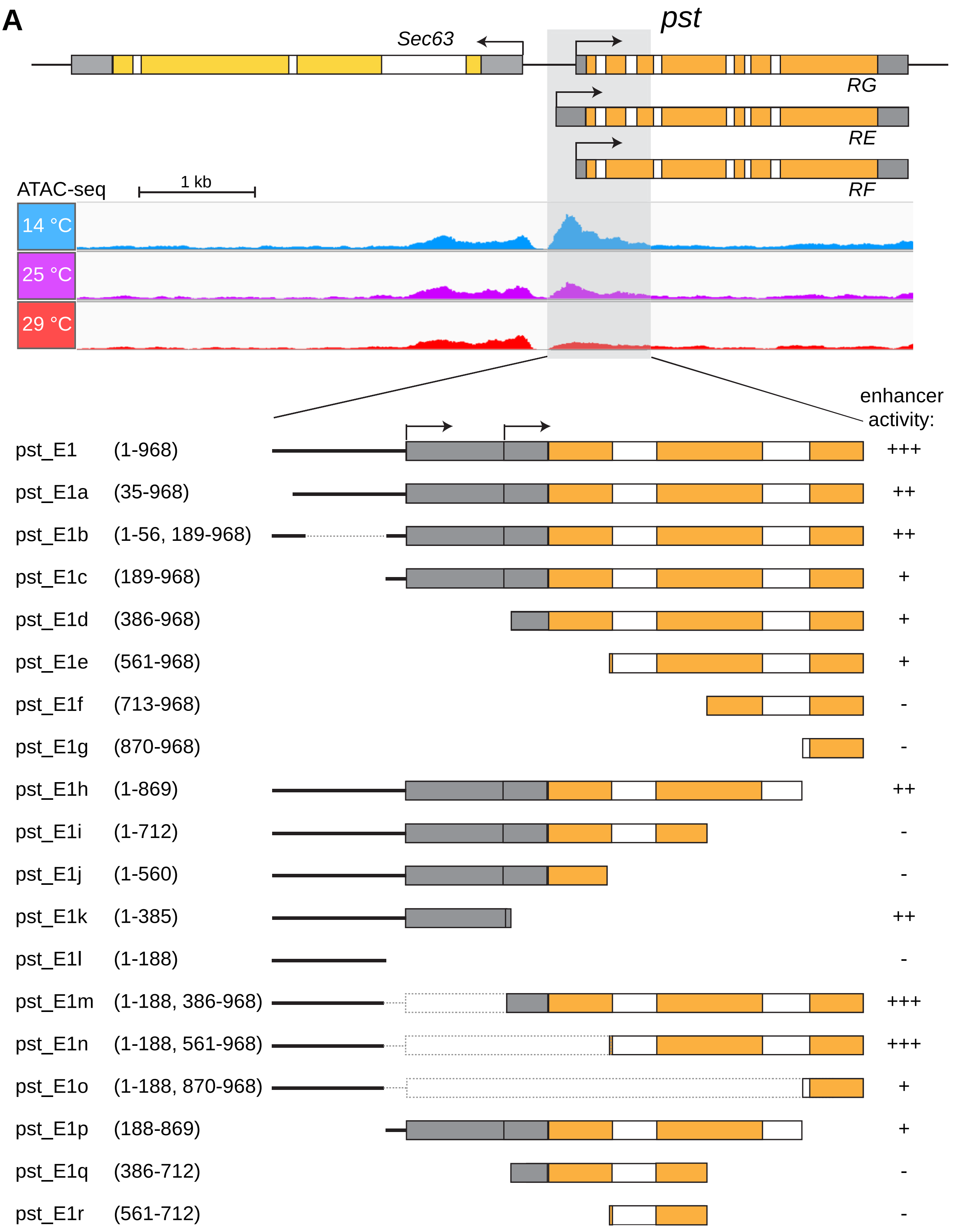

Bai et al., S10 Fig.

**B**

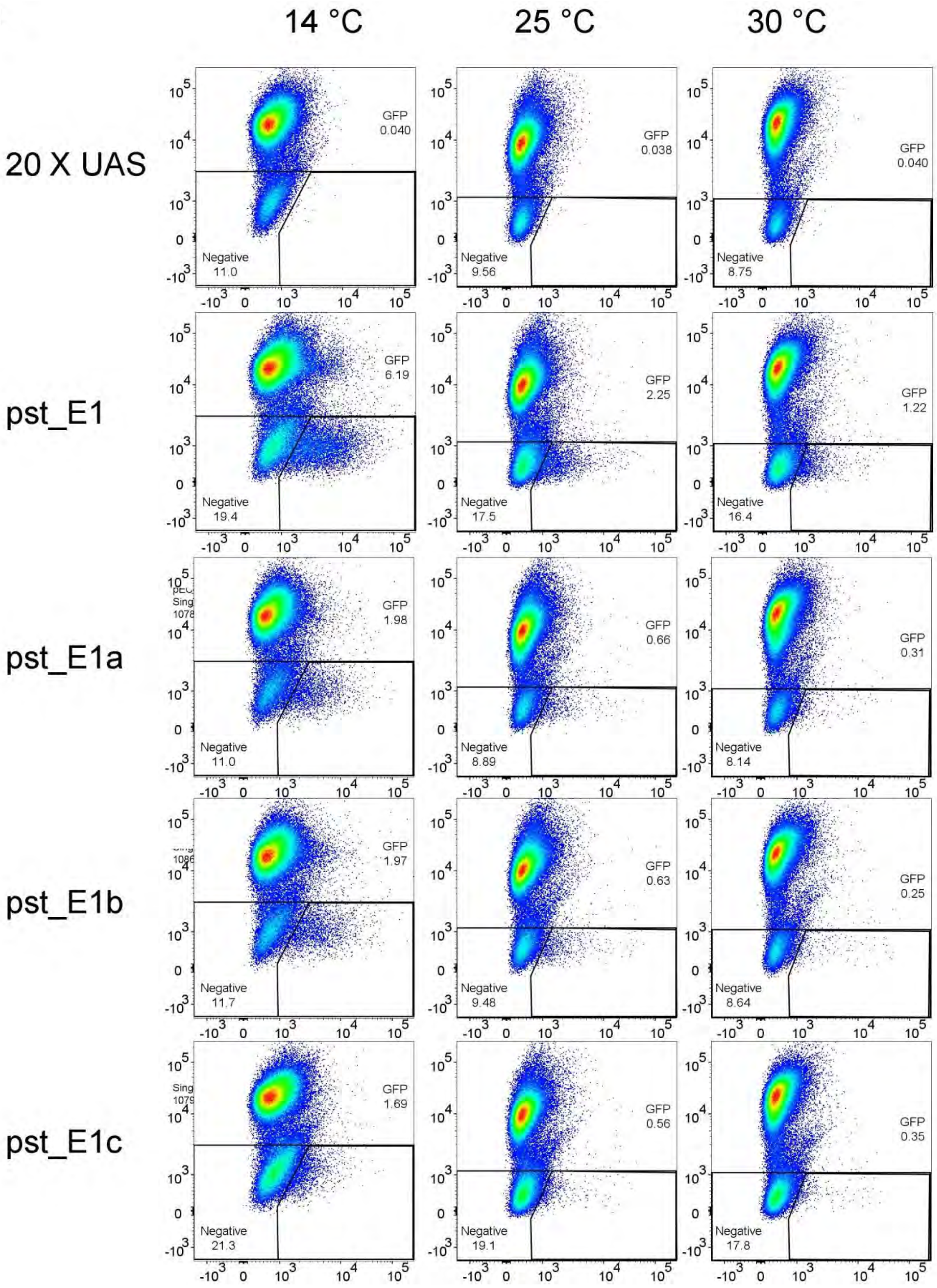

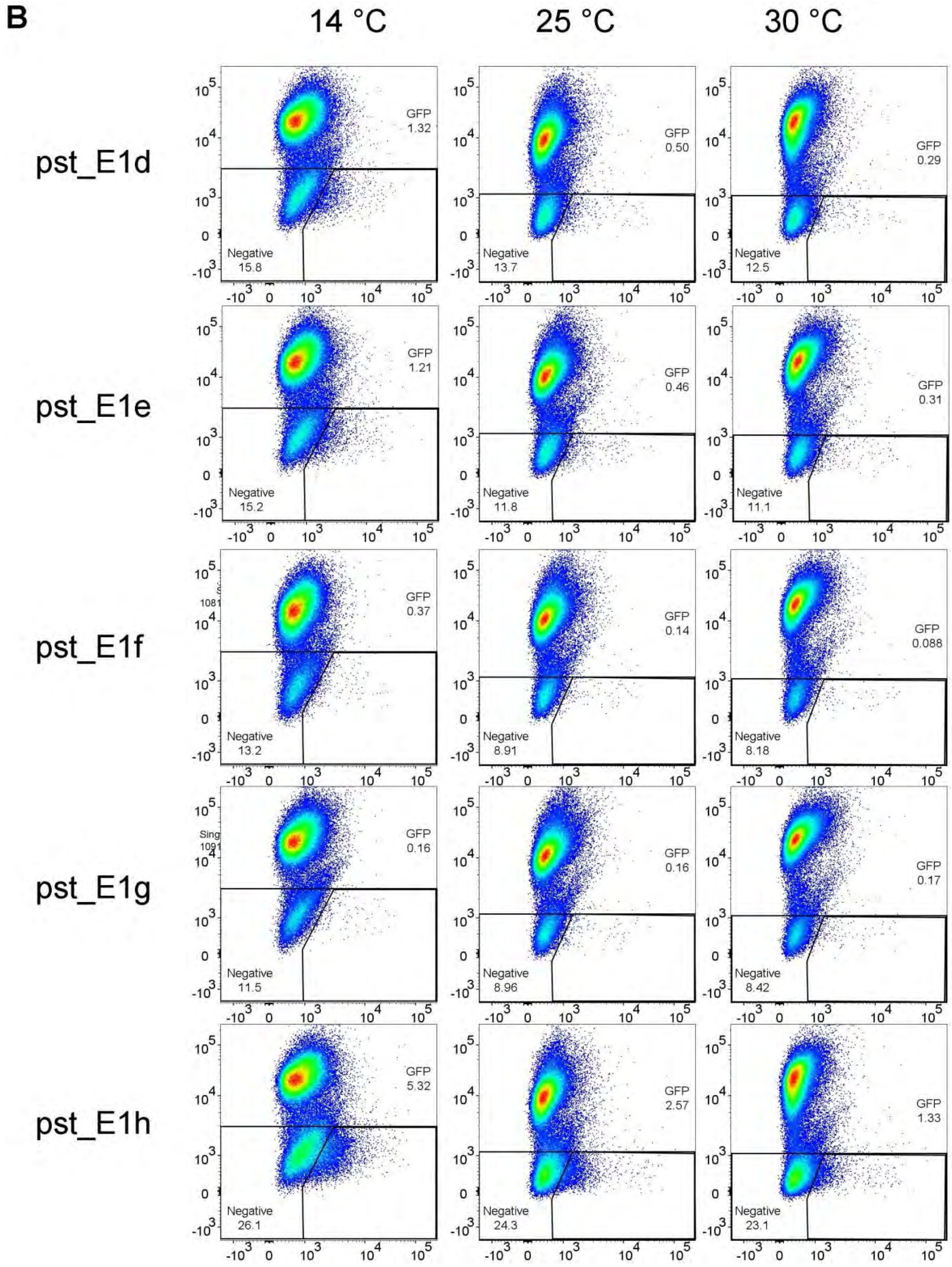
B
14 °C
25 °C
30 °C
pst_E1d
pst_E1e
pst_E1f
pst_E1g
pst_E1h
GFP 1.32
Negative 15.8
GFP 0.50
Negative 13.7
GFP 0.29
Negative 12.5
GFP 1.21
Negative 15.2
GFP 0.46
Negative 11.8
GFP 0.31
Negative 11.1
GFP 0.37
Negative 13.2
GFP 0.14
Negative 8.91
GFP 0.088
Negative 8.18
GFP 0.16
Negative 11.5
GFP 0.16
Negative 8.96
GFP 0.17
Negative 8.42
GFP 5.32
Negative 26.1
GFP 2.57
Negative 24.3
GFP 1.33
Negative 23.1

Bai et al., S10 Fig.

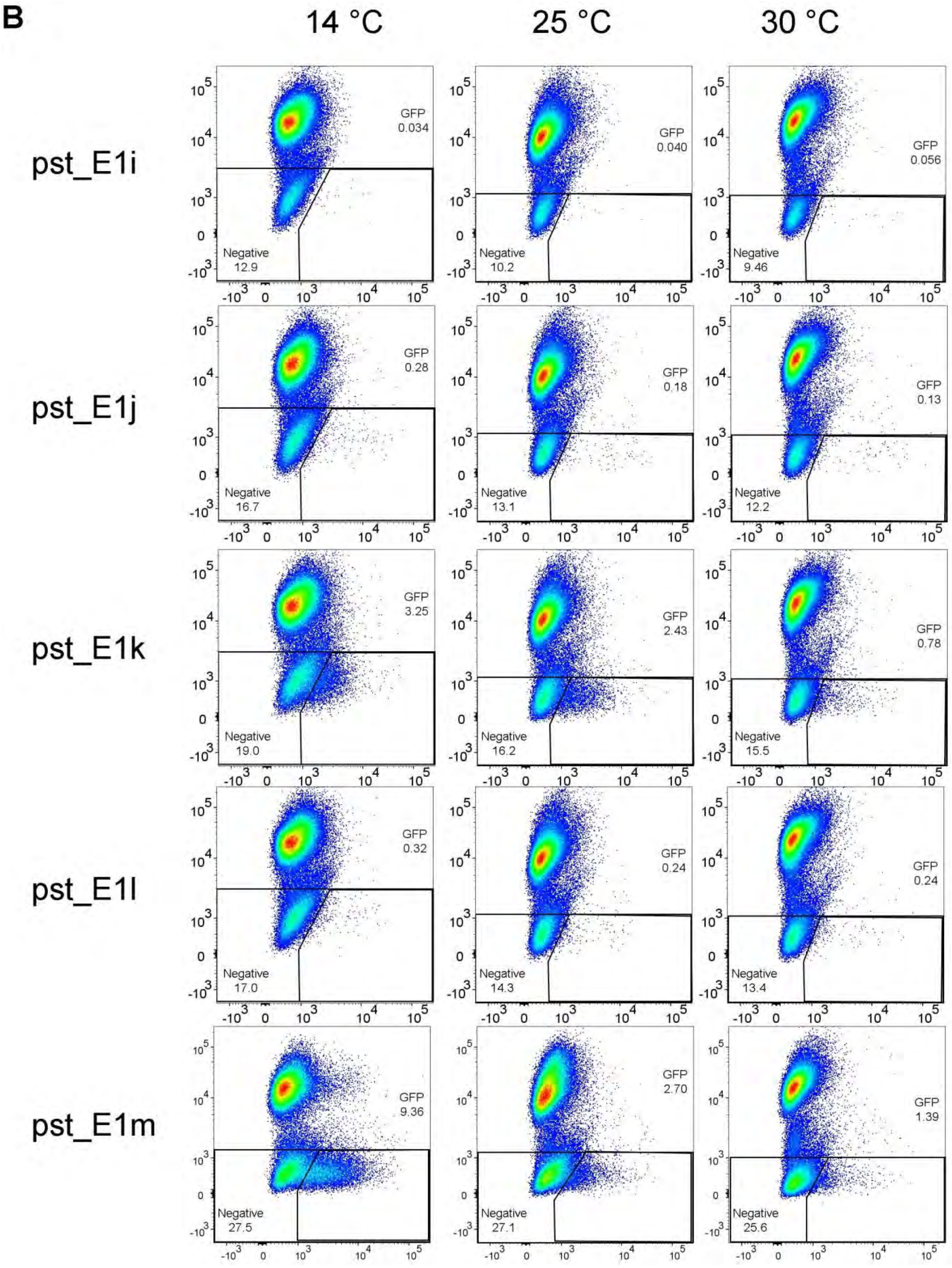

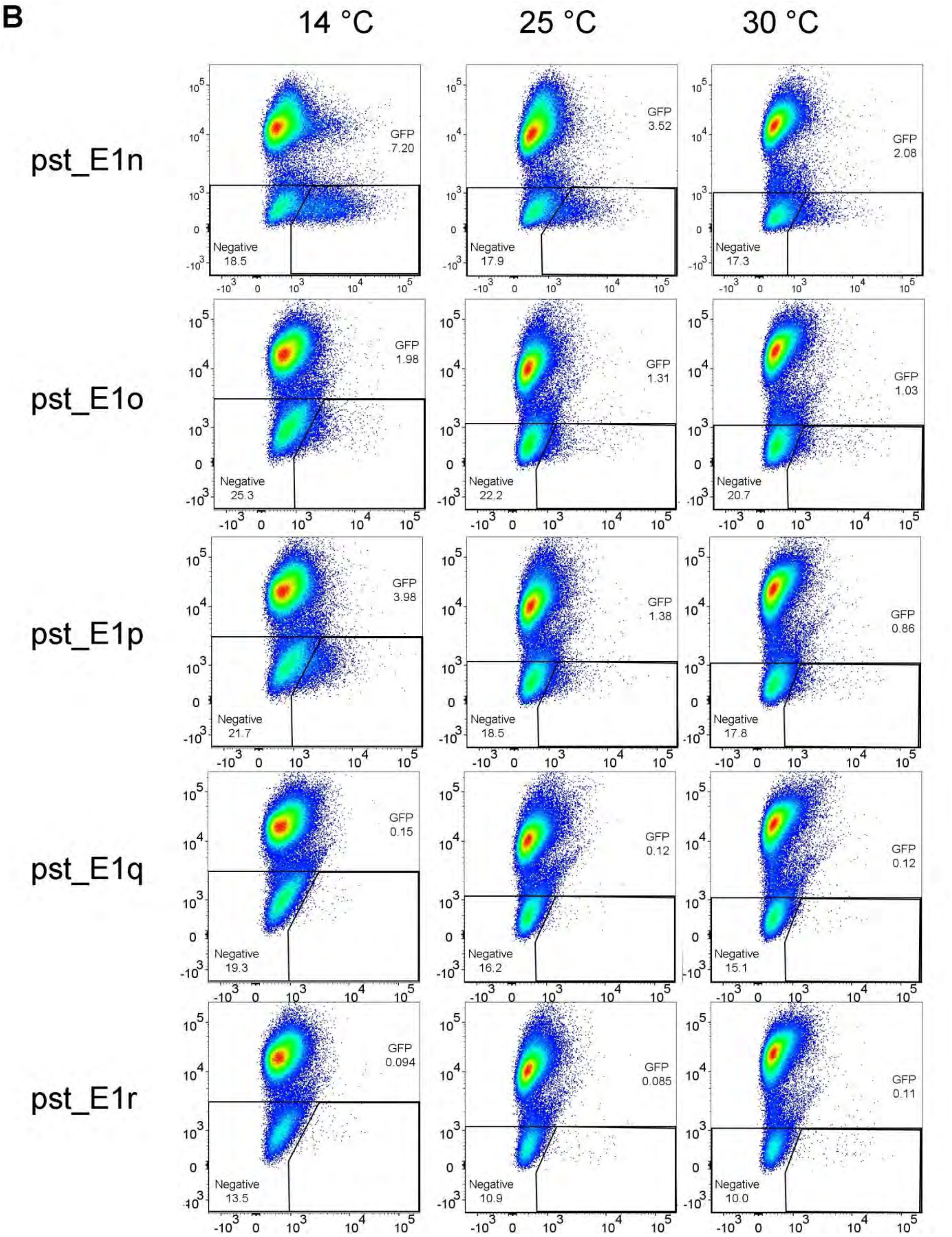


**S10 Fig. Dissection of *pst*_E1 region**

(**A**) Schematic illustration of analyzed truncation series, with transcription start sites (kinked arrows), untranslated regions (grey boxes), coding region (yellow boxes) and introns (white boxes) indicated.

(**B**) Scatter plots of the results obtained by flow cytometry after RMCE with SR9rg cells and incubation at the indicated temperatures.

Bai et al., S11 Fig.

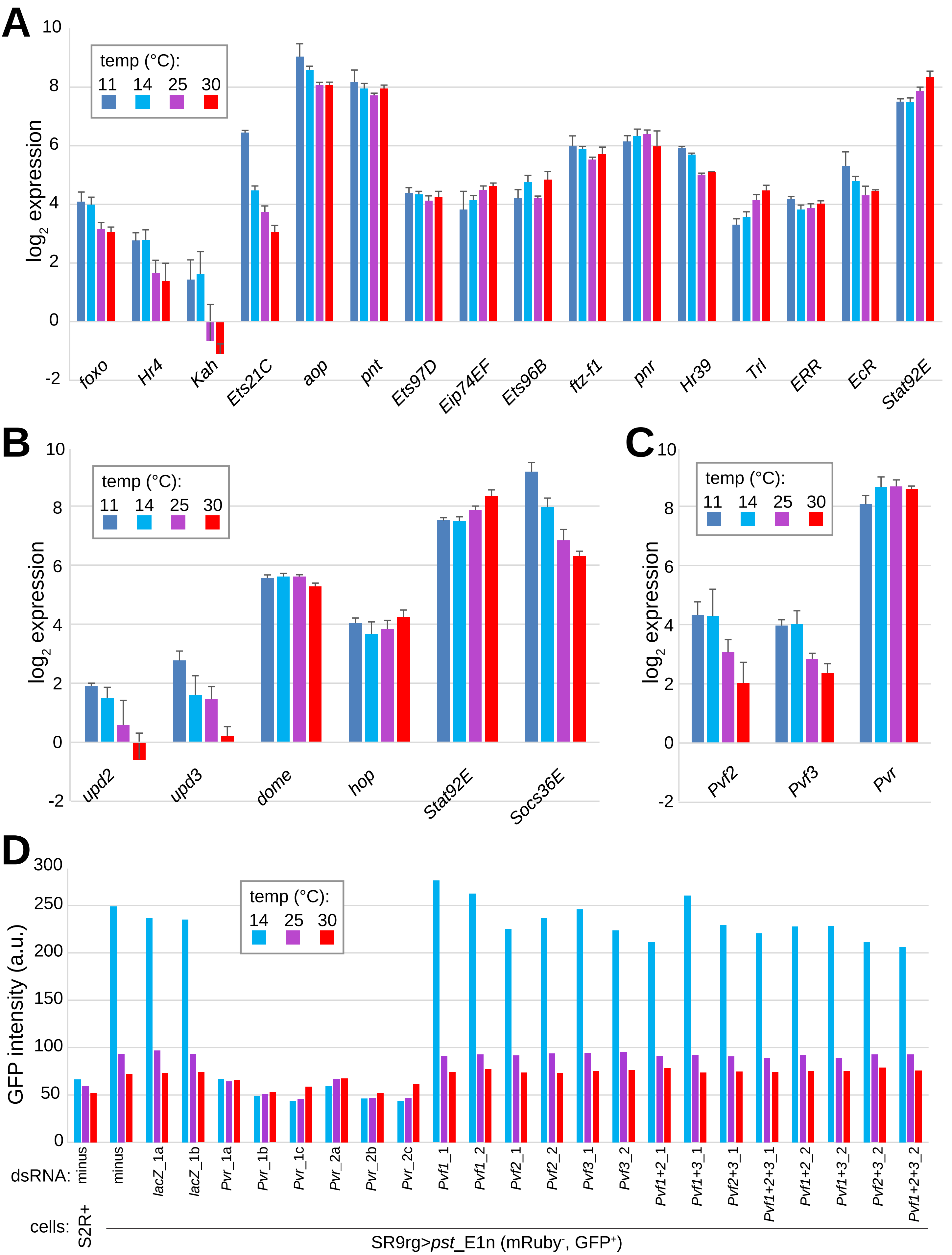

Bai et al., S12 Fig.

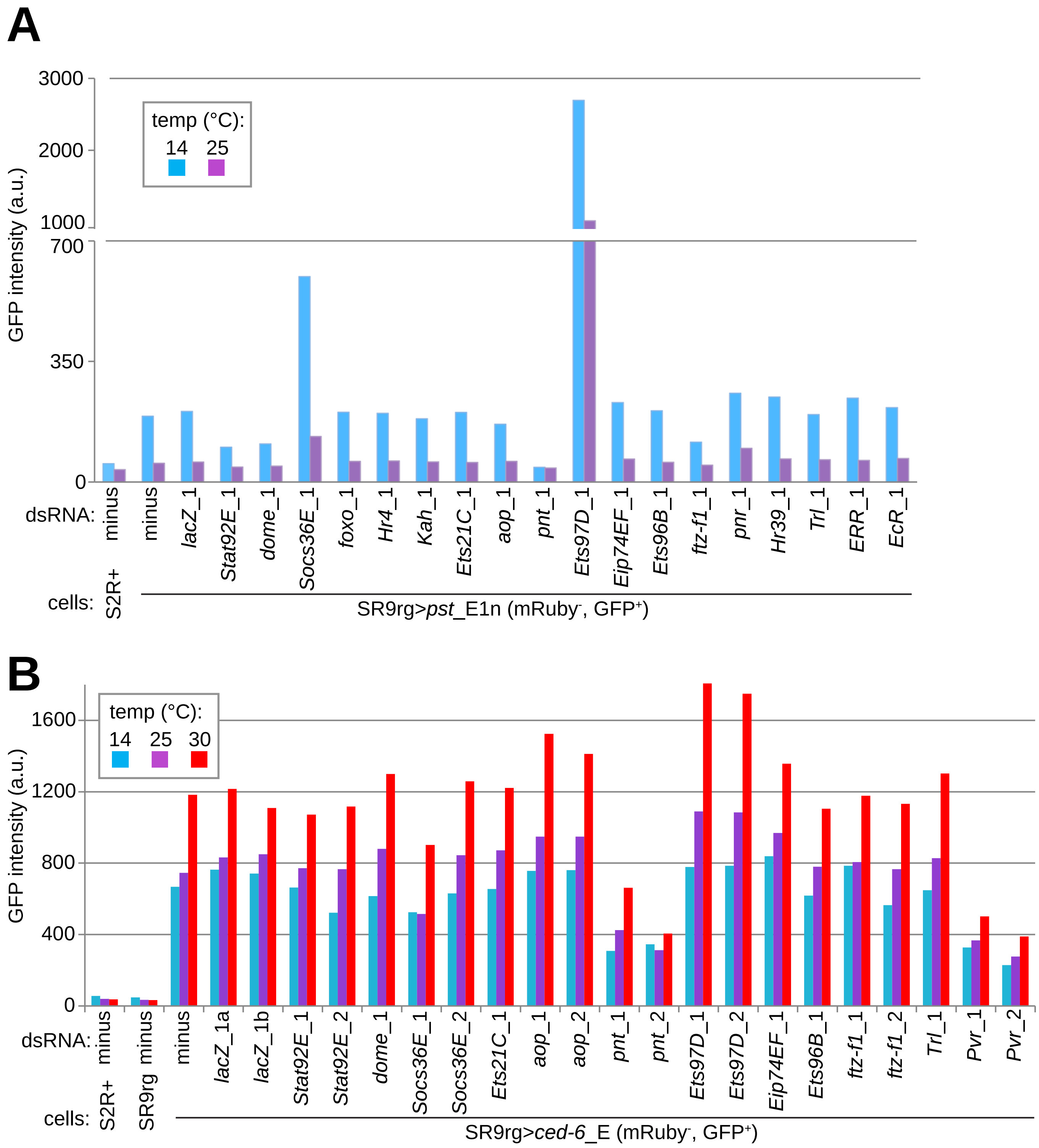

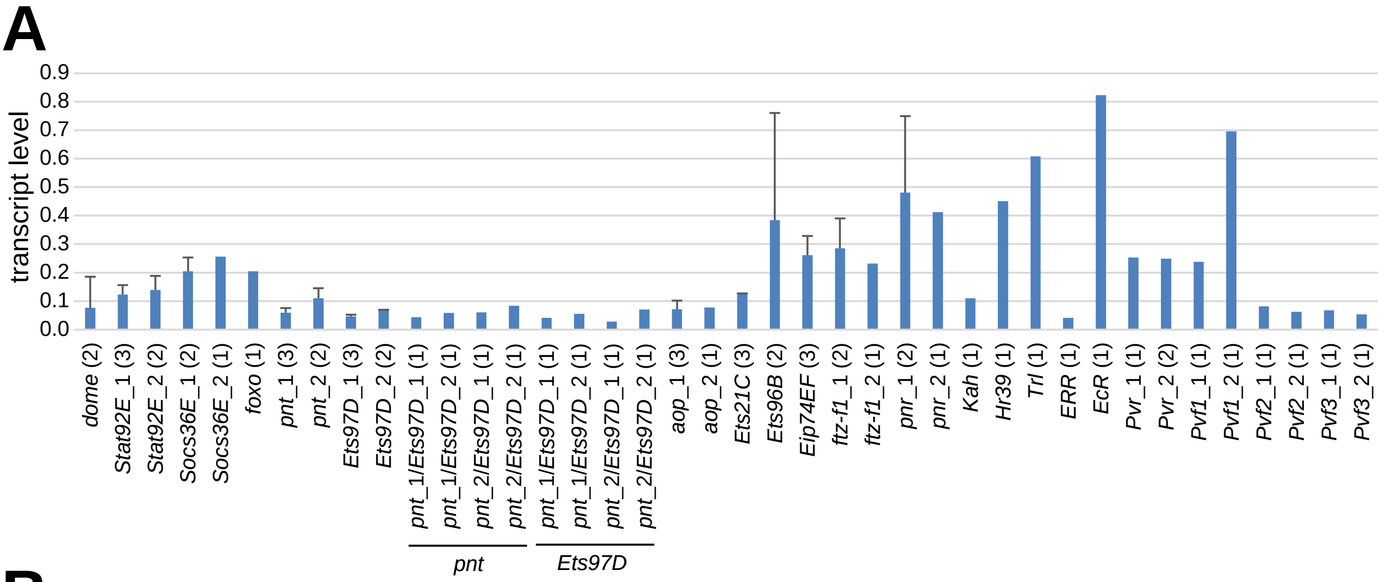
A
transcript level
0.0
0.1
0.2
0.3
0.4
0.5
0.6
0.7
0.8
0.9
dome (2)
Stat92E_1 (3)
Stat92E_2 (2)
Socs36E_1 (2)
Socs36E_2 (1)
foxo (1)
pnt_1 (3)
pnt_2 (2)
Ets97D_1 (3)
Ets97D_2 (2)
pnt_1/Ets97D_1 (1)
pnt_1/Ets97D_2 (1)
pnt_2/Ets97D_1 (1)
pnt_2/Ets97D_2 (1)
pnt_1/Ets97D_1 (1)
pnt_1/Ets97D_2 (1)
pnt_2/Ets97D_1 (1)
pnt_2/Ets97D_2 (1)
aop_1 (3)
aop_2 (1)
Ets21C (3)
Ets96B (2)
Eip74EF (3)
ftz-f1_1 (2)
ftz-f1_2 (1)
pnr_1 (2)
pnr_2 (1)
Kah (1)
Hr39 (1)
Trl (1)
ERR (1)
EcR (1)
Pvr_1 (1)
Pvr_2 (2)
Pvf1_1 (1)
Pvf1_2 (1)
Pvf2_1 (1)
Pvf2_2 (1)
Pvf3_1 (1)
Pvf3_2 (1)
pnt
Ets97D

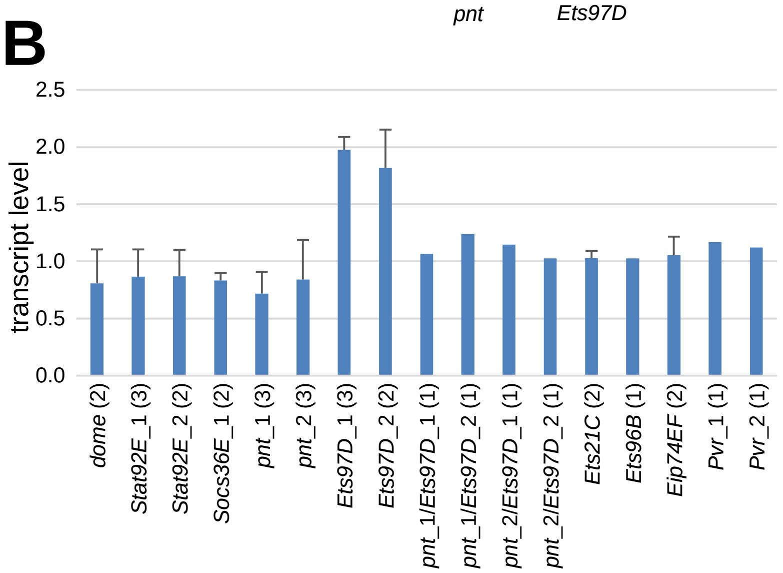
B
transcript level
0.0
0.5
1.0
1.5
2.0
2.5
dome (2)
Stat92E_1 (3)
Stat92E_2 (2)
Socs36E_1 (2)
pnt_1 (3)
pnt_2 (3)
Ets97D_1 (3)
Ets97D_2 (2)
pnt_1/Ets97D_1 (1)
pnt_1/Ets97D_2 (1)
pnt_2/Ets97D_1 (1)
pnt_2/Ets97D_2 (1)
Ets21C (2)
Ets96B (1)
Eip74EF (2)
Pvr_1 (1)
Pvr_2 (1)

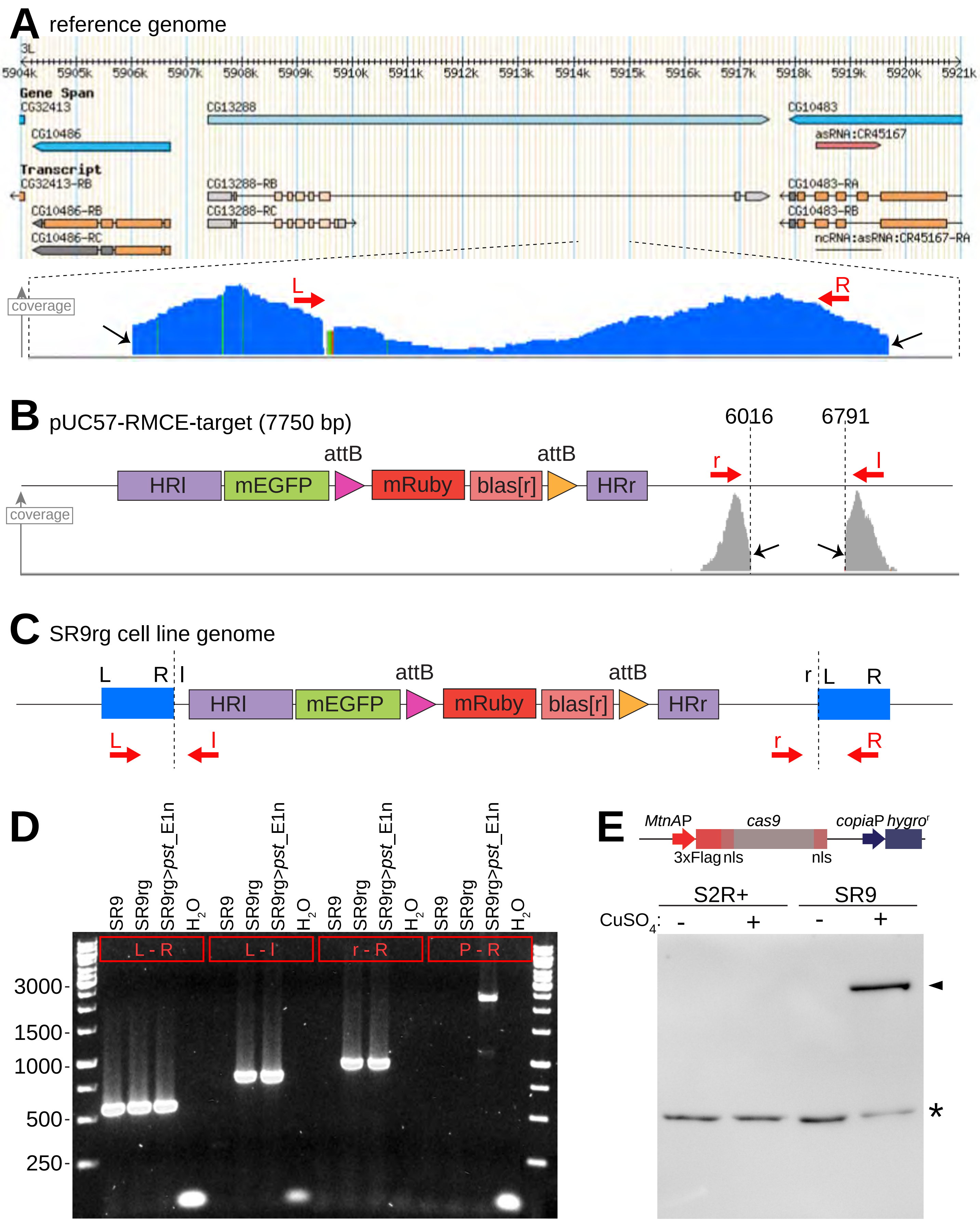
A reference genome
3L
5904k 5905k 5906k 5907k 5908k 5909k 5910k 5911k 5912k 5913k 5914k 5915k 5916k 5917k 5918k 5919k 5920k 5921k
Gene Span
CG32413
CG13288
CG10483
CG10486
asRNA:CR45167
Transcript
CG32413-RB
CG13288-RB
CG10483-RA
CG10486-RB
CG13288-RC
CG10483-RB
CG10486-RC
ncRNA:asRNA:CR45167-RA
coverage
L
R
B pUC57-RMCE-target (7750 bp)
6016
6791
attB
attB
HRl
mEGFP
mRuby
blas[r]
HRr
r
l
coverage
C SR9rg cell line genome
L
R
l
attB
attB
HRl
mEGFP
mRuby
blas[r]
HRr
r
L
R
L
l
r
R
D
SR9
SR9rg
SR9rg>pst_E1n
H2O
L - R
L - l
r - R
P - R
3000-
1500-
1000-
500-
250-
E
MtnAP
cas9
copiaP
hygror
3xFlag
nls
nls
S2R+
SR9
CuSO4:
-
+
-
+
*
